# Ferrous-glutathione coupling mediates ferroptosis and frailty in *Caenorhabditis elegans*

**DOI:** 10.1101/594408

**Authors:** Nicole L. Jenkins, Simon A. James, Agus Salim, Fransisca Sumardy, Terence P. Speed, Marcus Conrad, Des R. Richardson, Ashley I. Bush, Gawain McColl

## Abstract

All eukaryotes require iron. Replication, detoxification, and a cancer-protective form of regulated cell death termed *ferroptosis*^1^, all depend on iron metabolism. Ferrous iron accumulates over adult lifetime in the *Caenorhabditis elegans* model of ageing^2^. Here we show that glutathione depletion is coupled to ferrous iron elevation in these animals, and that both occur in late life to prime cells for ferroptosis. We demonstrate that blocking ferroptosis, either by inhibition of lipid peroxidation or by limiting iron retention, mitigates age-related cell death and markedly increases lifespan and healthspan in *C. elegans*. Temporal scaling of lifespan is not evident when ferroptosis is inhibited, consistent with this cell death process acting at specific life phases to induce organismal frailty, rather than contributing to a constant ageing rate. Because excess age-related iron elevation in somatic tissue, particularly in brain^3–5^, is thought to contribute to degenerative disease^6, 7^, our data indicate that post-developmental interventions to limit ferroptosis may promote healthy ageing.

---

Ferroptosis is a regulated cell death program genetically and biochemically distinct from apoptosis, necrosis and autophagic cell death. It kills malignant cells but may also be inappropriately activated in ischemic injury and neurodegeneration^8–11^. This cell death mechanism is executed by (phospho)lipid hydroperoxides induced by either iron-dependent lipoxygenases, or by an iron-catalyzed spontaneous peroxyl radical-mediated chain reaction (autoxidation). Under homeostatic conditions the ferroptotic signal is terminated by glutathione peroxide-4 (GPx4), a phospholipid hydroperoxidase that needs glutathione as a cofactor. While the signaling that regulates ferroptosis has been studied in depth^12–14^, the role of iron load in this death signal is poorly resolved^15^.

Redox cycling between Fe^2+^ and Fe^3+^ can contribute to cellular stress. This is mitigated by a range of storage and chaperone pathways to ensure that the labile iron pool is kept to a minimum^6^. In *Caenorhabditis elegans* the emergence of labile ferrous iron with age correlates with genetic effects that accelerate ageing^2, 16^ and could be a lifespan hazard^17^. Excess iron supply has been shown to shorten lifespan in *C. elegans*^18, 19^, however variable results have been reported with iron chelation. The iron chelator deferiprone was reported not to impact *C. elegans* lifespan^20^, but this study was limited by indirect measures of iron load, use of only a single dose of deferiprone, and small sample size. In contrast, use of calcium-ethylenediaminetetraacetic acid (CaEDTA), a non-specific chelator that does not redox-silence iron, caused a minor (undisclosed) increase in lifespan^21^. Whether selective targeting of ferrous iron burden can impact on ageing and lifespan is unknown.

We contemplated whether the developmental dependence on iron for reproduction and cellular biochemistry could represent an ancient and conserved liability in late life. The load of tissue iron increases needlessly in ageing nematodes, mammals and humans^2–5^. This must tax regulatory systems that prevent abnormal redox cycling of iron, such as the Fe^2+^-glutathione complexes considered the dominant form of iron in the cellular labile iron pool^22^. We hypothesized that age-dependent elevation of labile iron, coupled with a reduction of glutathione levels conspire to lower the threshold for ferroptotic signaling, increasing the vulnerability of aged animals and implying that disruption to the iron-glutathione axis is fundamental to natural ageing and death. To test this, we investigated the vulnerability to ferroptosis of ageing nematodes upon the natural loss of glutathione during lifespan. We examined the effects of inhibiting ferroptosis in *C. elegans* using two distinct treatments: a potent quenching agent for lipid peroxidation (autoxidation)^23^, as well as a small lipophilic iron chelator^24^ that prevents the initiation and amplification of lipid peroxide signals. Our analysis of these interventions provides mechanistic insight into the influence of the Fe^2+^-glutathione couple on the occurrence of natural ferroptosis across lifespan, at the cellular and organismal level.

## Results

### Glutathione depletion vulnerability

Glutathione is not only the dominant coordinating ligand for cytosolic ferrous iron^22^ but is also the substrate used by glutathione peroxidase-4 (GPX4) to clear the lipid peroxides that induce ferroptotic cell death^25–27^. Deletion of four *C. elegans* homologs of GPX4 decreases lifespan^28^, but whether ferroptosis is responsible is unknown. We tested whether acute depletion of glutathione can initiate ferroptosis in adult *C. elegans* using diethyl maleate (DEM), which conjugates glutathione^29, 30^. We found that DEM induced death in 4-day old worms (at the end of their reproductive phase) in a dose- and time-dependent manner, with ≈50% lethality occurring after 24-hour exposure to 10 mM DEM (**Figure 1A**) associated with ≈50% depletion of glutathione (**Figure 1B**). We also found that total glutathione levels steadily decrease with normal ageing, approaching ≈50% on Day 10 of the levels on Day 1 (**Figure 1C, Supplemental Table S1**), and that *C. elegans* become disproportionately more vulnerable to DEM lethality as they enter the midlife stage (**Figure 1D**).

**Figure 1.**
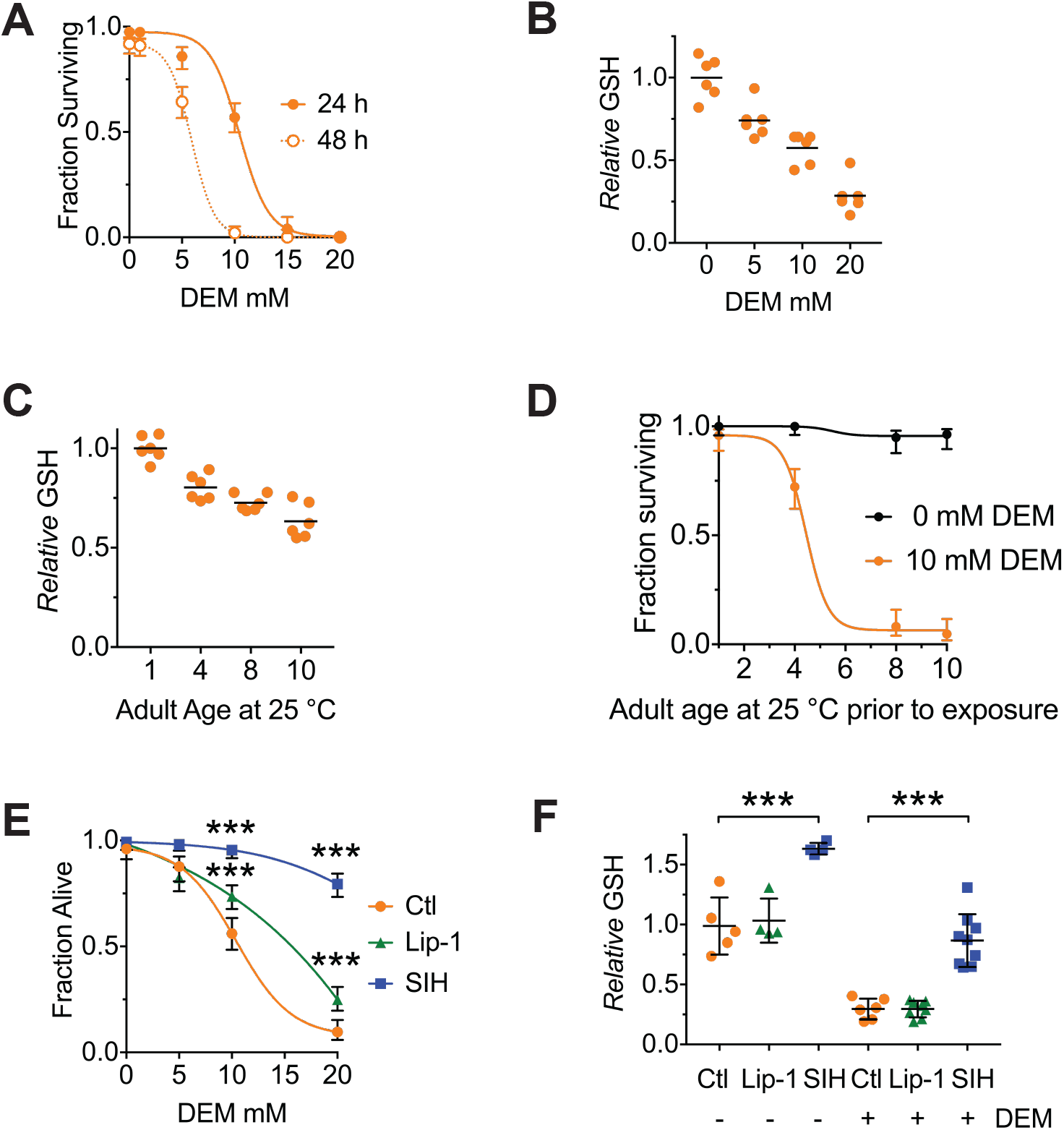
Both Lip-1 and SIH protect against toxicity from glutathione depletion. Treatment with DEM represents an acute stress that reduces glutathione levels and causes death, with older animals being more susceptible. **(A)** Survival curves of adults following either 24- or 48-hour exposure to increasing doses of DEM. Treatment begun on Day 4 of adulthood. Shown are proportions ± 95% confidence intervals^46^, with a sigmoidal curve fitted. **(B)** Total glutathione (GSH) decrease following 6 hours of DEM exposure. Day 4 adults used, with results normalized to the GSH levels in worms not exposed to DEM. Plotted are 9 independent replicates, with each estimated derived from 50 adults per measure). Linear regression R^2^= 0.98, *p* = 0.01 **(C)** Total GSH levels decrease with increased adult age in *C. elegans*. Each point is derived from independent replicates of 50 adults, with black lines marking the mean value. Results are normalized to the GSH levels in Day 1 worms (ANOVA: F (3, 20) = 32.96, *p*< 0.0001; see **Supplemental Table S1** for pairwise comparisons). **(D)** Aged *C. elegans* adults become progressively more sensitive to GSH depletion by DEM. Shown are proportions ± 95% confidence intervals, with a sigmoidal curve fitted. **(E)** Both Lip and SIH treatment protect against lethality from DEM derived glutathione depletion. Day 4 adults, with values representing pooled data from four independent experiments ± 95% confidence intervals, each with a fitted sigmoidal curve. Pairwise comparisons at 10 and 20mM DEM were performed using Fisher’s exact test; *** denotes *p*<0.001. **(F)** Total glutathione levels are preserved following SIH pretreatment, but not by Lip-1. Day 4 adults were exposed to DEM (10 mM) for 6 hours, and total glutathione (GSH) assayed. Each point is derived from independent replicates of 50 adults, with black lines marking the mean value ± SD. (ANOVA: F (5, 30) = 50.97, *p*< 0.0001; see **Supplemental Table S2** for pairwise comparisons). *** denotes p<0.001

We tested whether lethality associated with glutathione depletion was caused by ferroptosis. We examined the treatment of *C. elegans* with the selective ferroptosis inhibitor, liproxstatin (Lip-1, 200 µM) ^1^. We also targeted the accumulation of late life iron^2, 16^, that catalyses (phospho)lipid hydroperoxide propagation, using salicylaldehyde isonicotinoyl hydrazone (SIH, 250 µM), a lipophilic acylhydrazone that scavenges intracellular iron and mobilizes it for extracellular clearance^24^. Importantly, unlike chelators such as CaEDTA, SIH binds iron in a redox-silent manner^31^. For both interventions, *C. elegans* were treated from early adulthood (late L4) onwards to eliminate any potential developmental effects.

DEM toxicity in 4-day old worms was rescued by both Lip-1 and SIH (**Figure 1E, Supplemental Table S2**), with more marked protection by SIH. This is consistent with ferroptosis contributing to the death mechanism. Therefore, the fall in glutathione with ageing (**Figure 1C, Table S1**) would be expected to interact synergistically with the concomitant rise in labile iron^2, 16^ to increase the risk of ferroptosis. We found that this age-dependent rise in iron itself contributes to the fall in glutathione, since pretreatment of the worms with SIH from L4 prevented the age-dependent decrease in glutathione when assayed on Day 4 of adult life (**Figure 1F**). Furthermore, SIH mitigated the glutathione depletion induced by DEM in Day 4 animals (**Figure 1F**), demonstrating that cytosolic iron synergizes the depletion of glutathione initiated by DEM. While Lip-1 alleviated the lethality of DEM (**Figure 1E**), it did not prevent the fall in glutathione that was induced by ageing (as assayed on Day 4) or by DEM (**Figure 1F**). Thus, Lip-1 inhibition of ferroptosis in *C. elegans* occurs downstream of glutathione depletion, consistent with its effect in rescuing ferroptosis in cultured cells^32^.

### Individual cell ferroptosis heralds organismal demise

A feature of ferroptosis is the propagation of cell death in a paracrine manner mediated by uncertain signals that might include the toxic lipid peroxidation end-products 4-hydroxynonenal (4-HNE) and malondialdehyde (MDA)^33, 34^. Compared to strong oxidants like the hydroxyl radical, 4-HNE and MDA are relatively stable and able react with macromolecules, such as proteins distal to the site of origin. To determine whether individual cell death precedes organismal death in our model of ageing, we used propidium iodide to visualize dead cells *in vivo* after DEM treatment and during ageing. Propidium iodide is a fluorescent intercalating agent that binds to DNA, but cannot cross the membrane of live cells, making it possible to identify the nuclei of recently dead or dying cells, as shown in **Figure 2A**. Examination of aged cohorts, or young animals treated with DEM, indicated that cell death (particularly death of intestinal cells) preceded organismal death in both 4 day old (**Figure 2B**) and 8 day old (**Figure 2C**) adults, and was significantly attenuated by both Lip-1 or SIH. Thus, the animal dies cell by cell, rather than in a single event, and this progressive degeneration is likely to contribute to the frailty phenotype.

**Figure 2.**
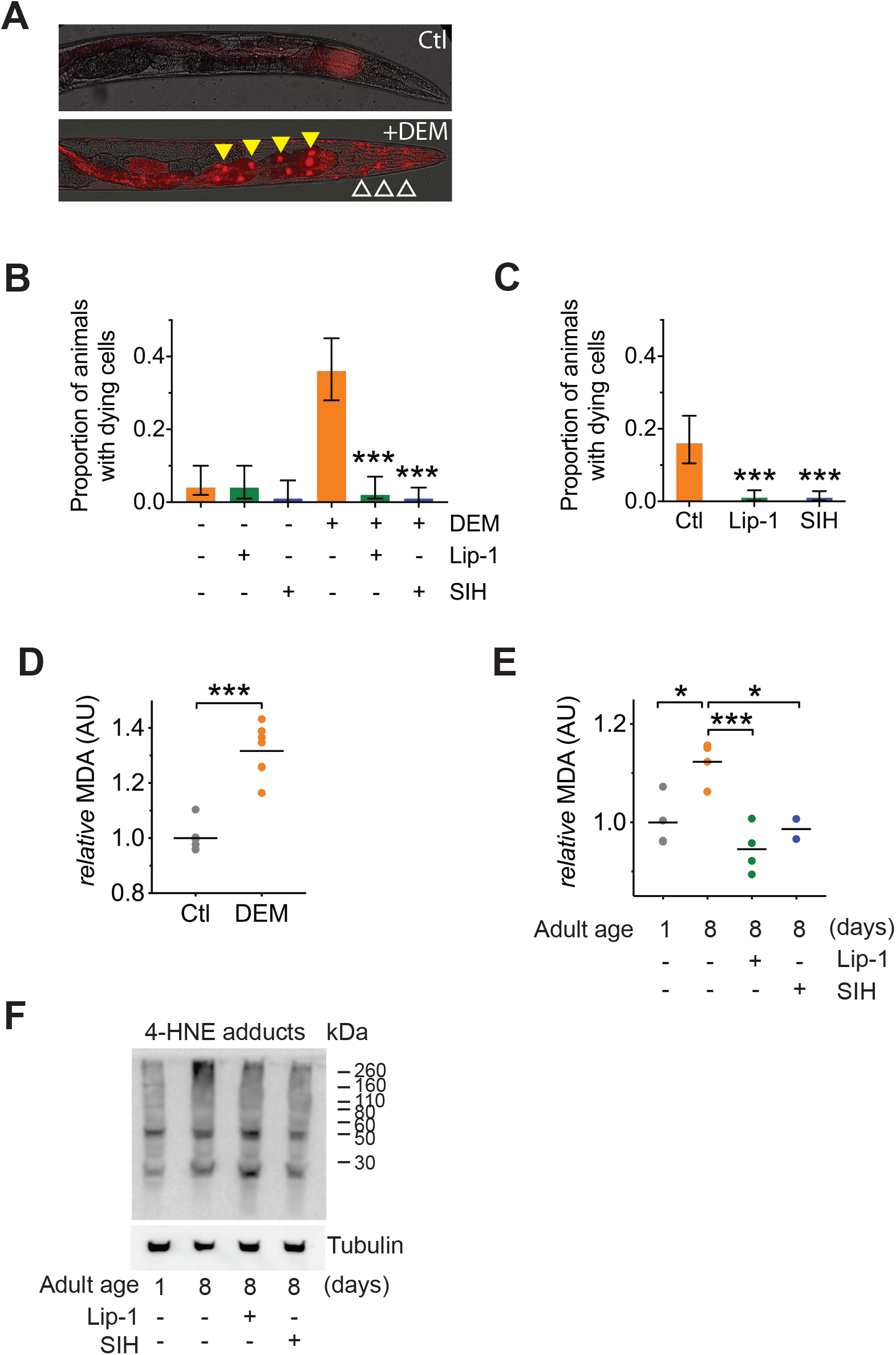
Both Lip-1 and SIH inhibit cell death and protect against lipid peroxidation. Measures of lipid peroxidation and cell death show an increase with age and reduction by both Lip-1 and SIH treatment. **(A)** Representative propidium iodide fluorescence (red) overlay of bright field micrograph depicting dead intestinal cells (marked by nuclear signal, yellow triangles) within a live Day 4 adult treated with DEM. Smaller fluorescent puncta were also observed, consistent with neuronal cell nuclei (white unfilled triangles). Untreated Day 4 adult control animals (Ctl) showed no cell death. **(B)** Proportion of live animals at Day 4 (± 95% confidence intervals) of adulthood showing dead cell fluorescence (propidium iodide) ± exposure to 10 mM DEM for 24h. Cohorts of animals included: co-treatment with a vehicle control (-DEM, Ctl, *n* =102; +DEM Ctl *n* =117), Lip-1 (-DEM, Lip-1, *n* =84; +DEM Ctl *n* =106) or SIH (-DEM, Ctl, *n* =89; +DEM Ctl *n* =129). Lip-1 and SIH both markedly reduced the proportion of animals with dead cells after DEM treatment (z-test: Ctl vs Lip-1 Z=6.37 *** p<0.001; Ctl vs SIH Z=7.24, *** p<0.001). **(C)** Proportion of live animals at Day 8 (± 95% confidence intervals) of adulthood showing dead cell fluorescence (propidium iodide). Co-treatment with Lip-1 (*n* =234) or SIH (*n* =308) markedly reduced the proportion of animals with dying cells compared to vehicle control (Ctl, *n* =119; z-test: Ctl vs Lip-1 Z=5.67 *** p<0.001; Ctl vs SIH Z=6.28, *** p<0.001). **(D)** Levels of the lipid peroxidation end product malondialdehyde (MDA) increases in *C. elegans* following acute glutathione depletion by 20 mM DEM exposure for 6h. MDA levels are shown as values normalized against the mean of untreated Day 4 Adults (Ctl) for independent samples. (Ctl vs +DEM, unpaired 2-tailed t-test *** p<0.001) **(E)** Malondialdehyde (MDA) increases in aged *C. elegans* (Day 1 vs Day 8 adults, ANOVA * p<0.05). Treatment with either Lip-1 (Day 8 vs Day 8 +Lip-1 adults, ANOVA *** p<0.001) or SIH (Day 8 vs Day 8 +SIH adults, ANOVA * p<0.05) reduces levels of MDA. Data represent independent samples with values normalized against the mean of untreated Day 1 Adults. **(F)** Representative immunoblot against 4-HNE protein adducts comparing Day 1 control and Day 8 control adults and aged adults treated with Lip-1 and SIH with corresponding tubulin blot below (representative of triplicate experiments). The relative intensity of the bands show an age-related increase that is ameliorated by Lip-1 and SIH.

To estimate changes in lipid peroxidation, we assayed MDA via the thiobarbituric acid reactive substance assay. As expected, acute glutathione depletion by DEM exposure caused a marked increase in the relative amounts of MDA (**Figure 2D**). We also observed an aged-related increase in MDA, consistent with an age-related increase in ferroptotic signalling in *C. elegans*, that was ameliorated by both Lip-1 and SIH treatment (**Figure 2E**). Consistent with the MDA results, we also found a concomitant qualitative increase in 4-HNE protein adducts with age that was suppressed by both Lip-1 and SIH treatments (**Figure 2F**).

### Changes in iron quantity, speciation and cytoplasmic fraction

Lowering cellular iron suppresses ferroptosis, but the peroxyl radical trapping ferroptosis inhibitors, such as Lip-1, are not expected to change iron levels. We examined the impact of SIH and Lip-1 interventions on iron levels over lifespan using synchrotron-based X-ray fluorescence microscopy^2, 35^ to measure both iron concentration (presented as areal density, pg µm^−2^) and total (pg per worm) iron (**Figure 3A**; **Supplemental Figure S1**). Both total iron and areal density increased with age in control animals (**Figure 3B & C; Supplemental Tables S3&S4**), as expected^2^. SIH dramatically reduced the areal density of iron (and reduced variance) with ageing (**Figure 3C**; **Supplemental Table S3&S4**), but Lip-1 did not alter iron density. Notably, by Day 8, animals treated with SIH contained total iron load on par with the untreated control group (**Figure 3C**; **Supplemental Tables S5&S6**), as the lower areal density was offset by an increase in body size of SIH-treated worms compared to age matched controls. These results highlight how bulk measures of total iron or measurements by inference^20^ can be confounded by changes in the animal morphology when exploring ageing interventions.

**Figure 3.**
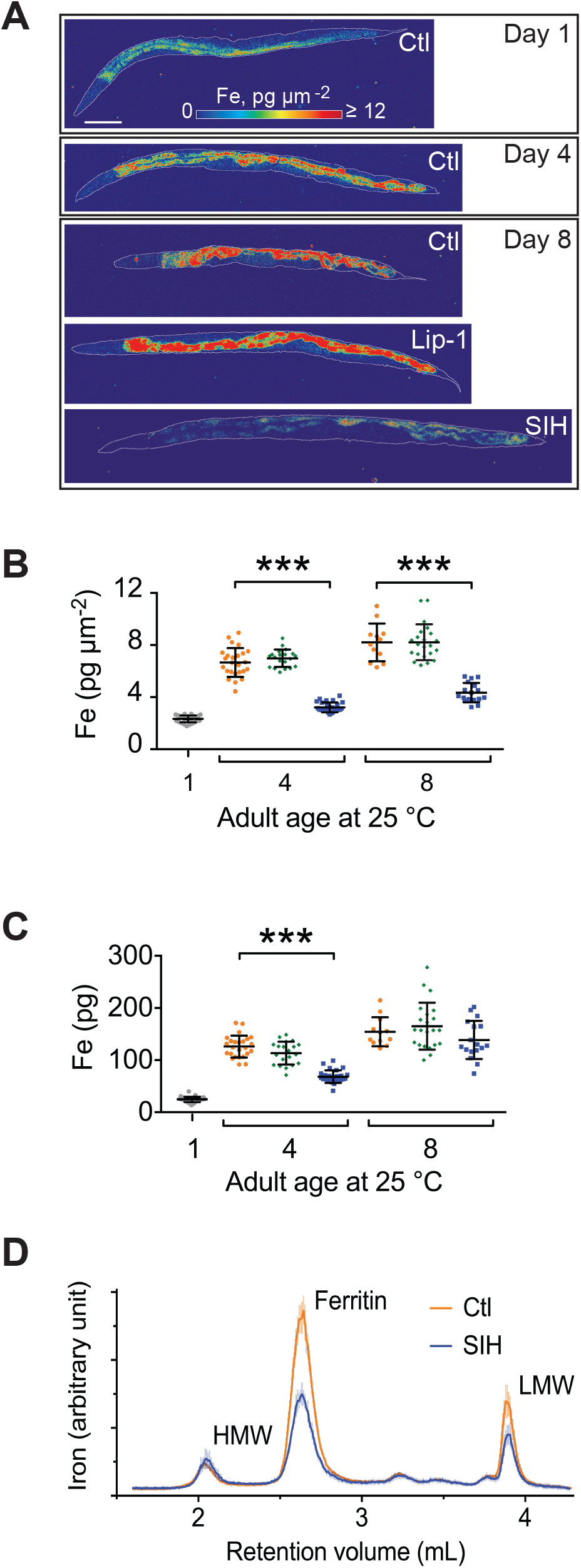
Effects of Lip-1 and SIH on iron levels and distribution in *C. elegans*. In all panels, vehicle control (0.5% v/v DMSO, Ctl) treated worms are shown in orange, Lip-1 treated (200 µM Lip-1) are green and SIH treated (250 µM SIH) are blue. **(A)** Representative X-ray fluorescence microscopy maps of tissue iron (Fe) reported as areal density (pg µm^−2^) for a first day adult (Day 1) and animals treated for four days (Day 4) and eight days (Day 8) with vehicle control (Ctl), Lip-1 or SIH at 25 °C. Scale bar = 50 µm. **(B)** Plot of mean areal density for iron (pg µm^−2^) for all treatment cohorts aged at 25 °C. The starting population (Day 1) is shown in grey. The control cohort (orange) shows an age related increase in total iron (as previously observed^2^). The Lip-1 group (green) has similar iron levels across each age, whereas the SIH cohort (blue) has markedly less total iron (ANOVA: F(6,148)=171.3, *p* < 0.0001; see **Supplemental Table S3** for sample summary and **Table S4** for pair-wise comparisons). Each data point represents a value from a single *C. elegans* adult, with mean ± SD, *** *p*<0.001. **(C)** Plot of total body iron (pg) for treated *C. elegans* cohorts aged at 25 °C. Each data point represents a value from a single *C. elegans* adult, with mean ± SD. All treatments have increased total iron across age with SIH treated (blue) worms retaining significantly less iron than control (red) and Lip-1 (green) treated worms at Day 4 (ANOVA: F(6,148)=97.3, *** *p* < 0.0001; see **Supplemental Table S5** for sample summary and **Table S6** for pair-wise comparisons) **(D)** Native, size-exclusion chromatography of iron-macromolecular complexes from 10 day old adults treated with vehicle control (Ctl, shown in orange) or SIH treated cohorts (shown in blue). The means ± SD, from three independent biological replicates, are plotted. The three major peaks include unaltered high molecular weight complexes (HMW, >1 MDa, ∼2.2 mL retention volume), ferritin bound iron (∼2.7 mL retention volume; previously identified as FTN-2^2^; area under the peak decreased by ∼53% relative to Ctl) and low molecular weight iron complexes (LMW, <30 kDa, ∼3.9 mL retention volume, decreased ∼47% relative to Ctl).

We had previously determined age-related changes to the *C. elegans* iron-proteome, characterized on size exclusion chromatography by three major peaks: a high molecular weight peak (HMW, >1 MDa), ferritin, and a low MW peak (LMW, 600 Da) that may contain labile iron. With ageing, iron redistributes in *C. elegans* out of the ferritin peak (where it is sequestered in redox-silent storage reserves) and accumulates in the HMW and LMW peaks^2^. The chromatographic profile of aged *C. elegans* (10 days post adulthood) treated with SIH (**Figure 3D**) revealed decreased iron associated with the LMW peak (normalized peak area approximately 40%). Ferritin-bound iron was also similarly decreased by SIH (normalized peak area approximately 50%), but iron bound within HMW species was unaffected. The age-related changes in LMW iron are consistent with increased labile iron, which is withdrawn as the substrate for ferroptosis by SIH treatment.

### Fe^2+^ increase with ageing is normalized by liproxstatin and SIH

X-ray absorption near edge structure (XANES) spectroscopy, using fluorescence detection for visualization, directly assesses the *in vivo* coordination environments of metal ions in biological specimens (*φ*XANES)^16^. The centroid of the XANES pre-edge feature reflects the relative abundance of ferrous [Fe^2+^] and ferric [Fe^3+^] species^36^. Since Fe^2+^ in the labile iron pool is the specific substrate for ferroptosis, and rises with ageing in *C. elegans*^2^, we investigated the impact of our interventions using *φ*XANES^16^. This synchrotron-based spectroscopy allowed us to evaluate steady state iron speciation (Fe^2+^/ Fe^3+^) in a specific region (anterior intestinal, **Figure 4A**; **Supplemental Figure S2**) of intact, cryogenically-stabilized control, Lip-1 and SIH-treated worms. We found that the age-related increase in the Fe^2+^ fraction was normalized to that of a young animal by both Lip-1 and SIH treatments (**Figure 4B & C; Supplemental Figure S3A & B; Supplemental Table S7**).

**Figure 4.**
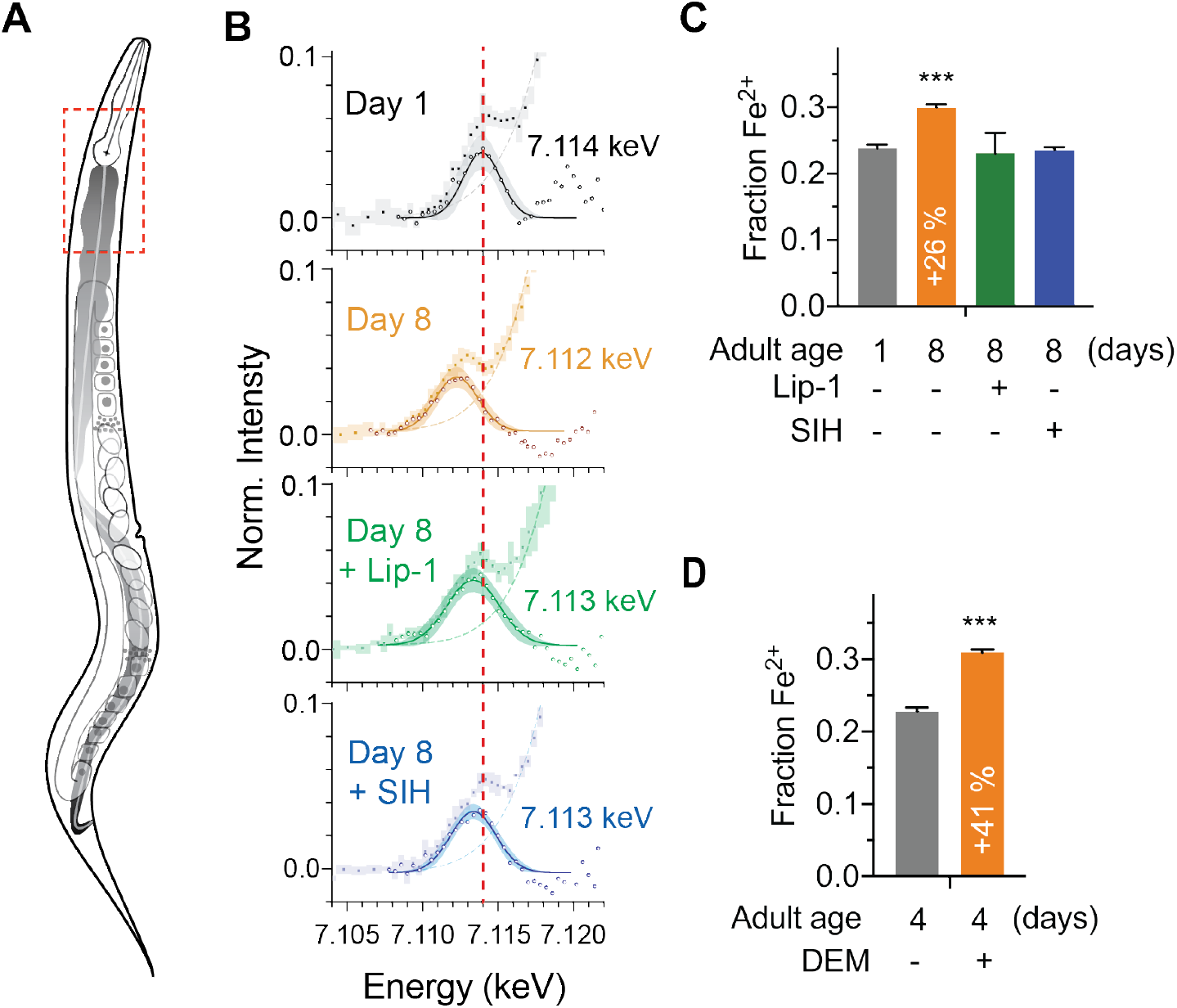
Effects of ageing, glutathione depletion, SIH and Lip-1 on pro-ferroptotic Fe^2+^ levels. *φ*XANES evaluation of the Fe^2+^ fraction in vivo within intact animals. **(A)** Schematic highlighting the anatomy of an adult hermaphrodite *C. elegans* with the intestine shaded in grey. Dashed box is indicative of the region of animals selected for *φ*XANES. **(B)** Fe^2+^ synchrotron microscopy. *φ*XANES imaging allowed extraction of the normalised Fe K-edge XANES spectra (coloured circles) from the anterior intestine of Day 1 (*n* = 6) and Day 8 control (*n* = 4), SIH treated (*n* = 4) and Lip-1 treated (*n* = 5) worms. Averaged spectra for each group are shown along with 95 % confidence intervals (shading). The pre-edge region, following subtraction of the rising edge (dashed line), highlights changes in the intensity and position of the 1s → 3d transition. The extracted data (empty circles) and fitted Gaussian (solid lines; shading represents the 95% CI) are superimposed to determine the centroid values for the pre-edge peak, from which the Fe^2+^ fraction is derived. Changes in the first derivative of the Fe K-edge XANES (**Supplemental Figure S3**) reflect variation in the intensity of the 1s → 4s and 1s → 4p transitions. The relative intensity of these features was used to estimate the proportion of Fe^2+^ iron in the specimens. For reference, the red line through all spectra denotes the centroid of the Day 1 adults at 7.114 keV. **(C)** The proportional change in fractional Fe^2+^ contribution for spectrum in the aged (Day 1 versus Day 8 adults) and treated (Lip-1 and SIH, from panel B) specimens is indicated, along with 95% confidence interval. Changes in the first derivative of the Fe K-edge XANES (**Supplemental Figure S3**) was used to infer variation in the intensity of the 1s → 4s and 1s → 4p transitions and the relative intensity of these features was then used to estimate the proportion of Fe^2+^ iron. **(D)** The proportional change in fractional Fe^2+^ contribution for Day 4 adults treated with (*n*= 4) and without acute glutathione depletion via DEM (*n*= 4) is indicated, along with 95% confidence interval.

Higher levels of pro-ferroptotic Fe^2+^ might be compounded by a loss of glutathione. So, we also assessed changes in fractional Fe^2+^ induced by lethal glutathione depletion by DEM. jXANES of 4 day old wild type worms treated with DEM identified a marked increase in the Fe^2+^ fraction (**Figure 4D; Supplemental Figure S3C & D; Supplemental Table S7**), revealing the upper limit for tolerable Fe^2+^ fraction being about 0.3 of the total iron (**Figure 4D**). These results help to contextualize the observed increase in Fe^2+^ during normal ageing also being about 0.3 of the total iron (**Figure 4C**), which was normalized to ≈0.2 by Lip-1 or SIH intervention.

### Lifespan effects of ferroptosis inhibition or blocking iron accumulation

Since Fe^2+^ accumulates with ageing and contributes to *C. elegans* frailty by executing cells before organismal death, we hypothesized that ferroptosis directly impacts on lifespan and may represent an underlying process that contributes to organismal ageing. We found that treatment of *C. elegans* with Lip-1 markedly extended lifespan [**Figure 5A&B**; average ∼70% increase in median lifespan (8 independent replicates; *p*<0.002), **Supplemental Table S8**]. Dose response is shown in **Supplemental Figures S4A & B**. An alternative ferroptosis inhibitor, ferrostatin^1^, was also examined, producing a significant but more modest median lifespan extension (**Supplemental Figure S4C**). Targeting the accumulation of late life iron using SIH also resulted in a marked increase in median lifespan [**Figure 5A&B**; average ∼100% median increase (8 independent replicates; *p*<0.0001), **Supplemental Table S8**]. Dose response is shown in **Supplemental Figure S4D**. Exposing *C. elegans* to 250 µM SIH as an iron complex (Fe(SIH)_2_NO_3_) neutralized the benefits of SIH on lifespan (**Supplemental Figure S4E**), confirming that the rescue mechanism required SIH being free to ligate iron.

**Figure 5.**
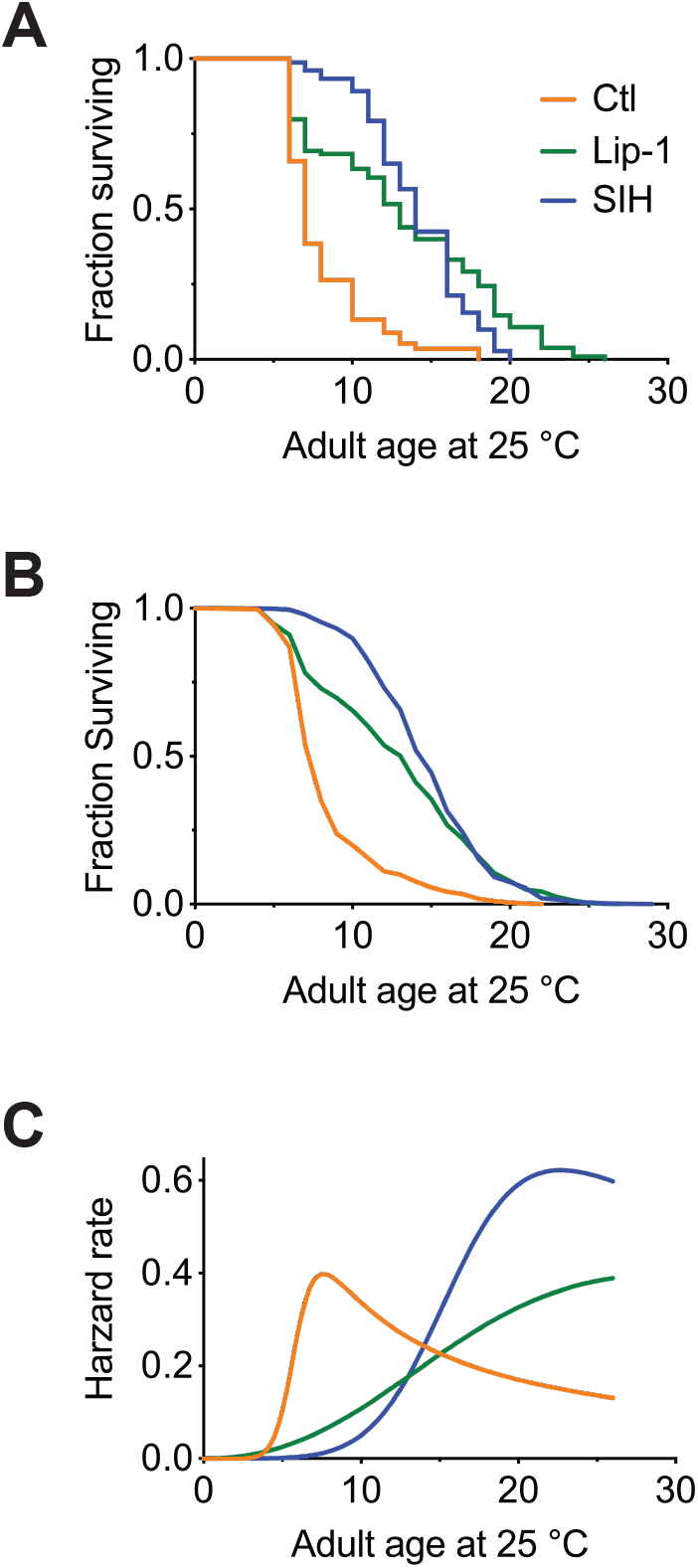
Inhibiting ferroptosis extends *Caenorhabditis elegans* lifespan. Treatment with both Lip-1 and SIH extend lifespan. **(A)** Representative Kaplan-Meier survival curve from *C. elegans* treated with vehicle control (Ctl, median survival 7 days, death event *n* = 88); Lip-1 (median survival 13 days, Log-rank test *p* < 0.001, *n* = 103) and SIH (median survival 14 days, *p* < 0.001, *n* = 71) at 25 °C. **(B)** Survival curve derived from pooled data from all eight replicate experiments (Ctl *n*= 709, Lip-1 *n*= 809, and SIH *n*= 720; see **Supplemental Table S8**) at 25 °C. **(C)** Plot of hazard (mortality) rate against age at 25 °C, derived from meta-analysis of pooled data (presented in b). Both SIH and Lip-1 alter mortality rates relative to control populations and are also distinct from each other. (see **Supplemental Tables S13-14**).

### Lifespan increases are not due to temporal scaling

Lip-1 and SIH had distinct and *unprecedented* effects on ageing, as shown by the lifespan curves in **Figure 5**. Treatment with Lip-1 primarily altered late life survival, while SIH extended mid-life with a squaring of the survival curve. Interventions that increase lifespan in *C. elegans* are not uncommon, but Stroustrup *et al*. recently demonstrated the great majority of longevity interventions *e.g*. dietary and temperature alteration, oxidative stress, and genetic disruptions of the insulin/IGF-1 pathway (*e.g*. *daf-2* and *daf-16*), heat shock factor *hsf-1*, or hypoxia-inducible factor *hif-1*, each alter lifespan by *temporal scaling* - an apparent stretching or shrinking of time^37^. For an intervention to extend lifespan by temporal scaling it must alter, to the same extent throughout adult life, all physiological determinants of the risk of death. In effect temporal scaling arises when the risk of death is modulated by an intervention acting solely on the rate constant associated with a single stochastic process.

Combining the replicate data from 8 independent experiments (**Figure 5B**), we assessed whether Lip-1 and SIH treatment effects can be explained by the temporal scaling model of accelerated failure time (AFT). We found that the lifespan increases were not consistent with the temporal scaling model (*p*<0.01; **Supplemental Tables S9-S14** and **Figures S5-S10**), so the interventions must ***target previously unrecognized ageing mechanisms***. For SIH treatment, the risk of death (hazard) in early adulthood was greatly reduced compared to control populations but rose precipitously in late life (**Figure 5C**). Lip-1 markedly reduced the rate of mortality in the post-reproductive period (late-life) with early life mortality closer to that seen in untreated populations. These findings are consistent with ferroptotic cell death limiting lifespan in late life rather than being a global regulator (*e.g.* insulin/IGF-1 pathway) of ageing. This raises the possibility of targeted intervention with minimal or no metabolic cost.

The sample size in our experiment is much smaller than the lifespan machine experiment undertaken by Stroustrup *et al*.^37^ yet the data against temporal rescaling were significant. To minimize the likelihood that our findings are due to either intrinsic bias in our experiment or inflation of effect size (the *Winner’s curse phenomenon*) we also examined the effect of temperature on lifespan intervention. Stroustrup *et al*.^37^ reported that changing temperature results in simple temporal rescaling of lifespans; our data corroborated this result, and showed that SIH still extended lifespan by a similar dimension at both 20°C and 25 °C (**Supplemental Tables S15-S17, Supplemental Figure S11**).

Our results indicate that while iron accumulation may impact many processes that influence ageing rate, ferroptosis inhibition predominantly reduces frailty rather than slows a global rate of ageing. Notably, Stroustrup *et al*. identified only two other instances among the many lifespan interventions tested in *C. elegans* that modulated lifespan outside a temporal scaling model, namely altered feeding behaviour (*eat-2* mutants) and mitochondrial dysfunction (*nuo-6* mutants)^37^. Both these mutants express marked developmental variability and reduced fitness.

### Preventing ferroptosis improves fitness and healthspan

Interventions that increase lifespan in *C. elegans* often do so at the detriment of fitness and healthspan. Adult body size can inform on fitness; reduced size may reflect a trade-off between longevity and fitness, as typically seen under dietary restriction where the cost of increased longevity can be lowered size, fertility and movement^38^. Distinctly, SIH-treated animals grew substantially larger. Following one day of treatment all animals were of similar body length (**Figure 6A & B**). After 4 days and 8 days of intervention, adult SIH-treated animals were significantly longer compared to similarly aged controls (*e.g.* control 1440 ± 123 µm *versus* SIH 1696 ± 64 µm, means ± SD on Day 8, *p* < 0.001). In addition, SIH induced an increase in body volume between Days 1 and 4, but not thereafter (**Figure 6C**). SIH-treated worms grew to greater volume than both control and Lip-1 treated worms at Day 4, indicating that preventing iron accumulation can improve animal robustness (for all comparisons see **Supplemental Tables S18-21**). Lip-1 had no effect on length or volume.

**Figure 6.**
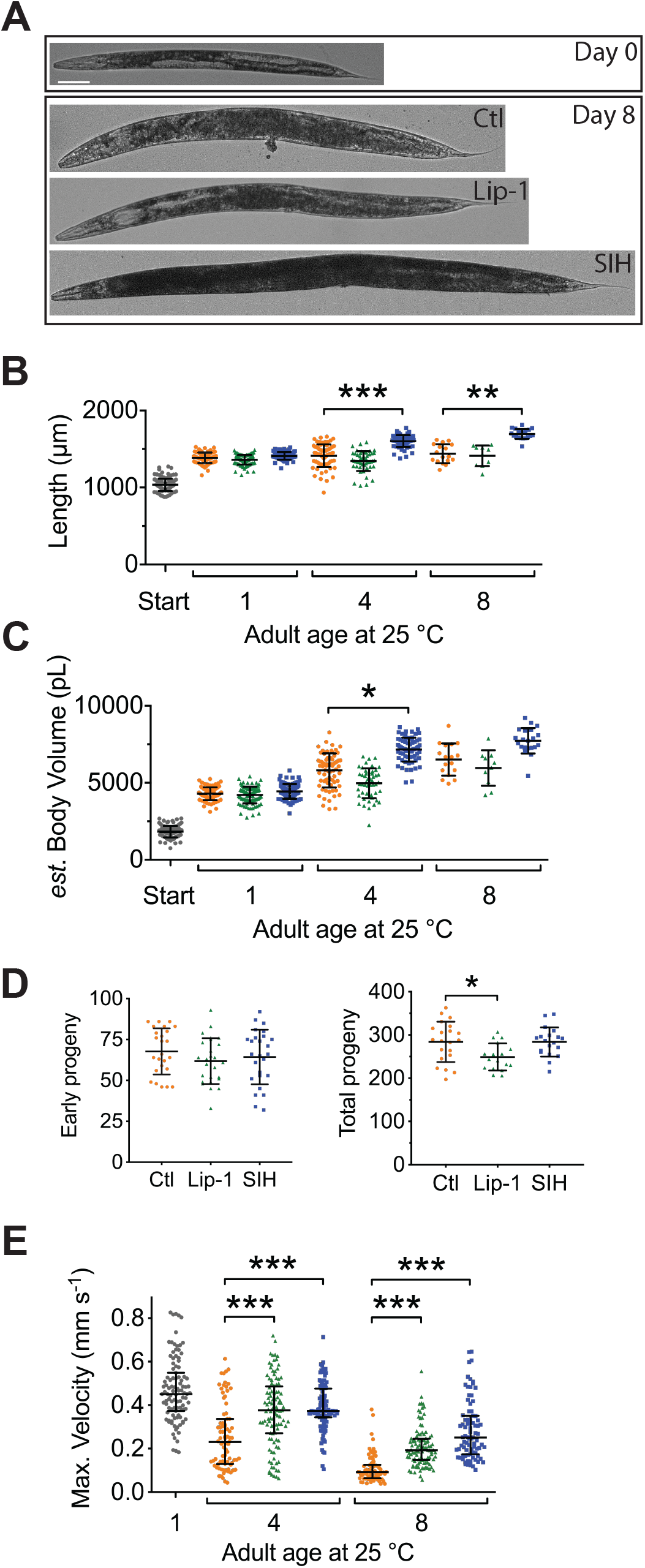
Lip-1 and SIH blocks frailty in *C. elegans*. In all panels, vehicle control (0.5% v/v DMSO, Ctl) treated worms are shown in orange, Lip-1 treated (200 µM Lip-1) are green and SIH treated (250 µM SIH) are blue. Significant differences between treatments are highlighted where * indicates *p*<0.05, ** indicates *p*<0.01 and *** indicates *p*<0.001. **(A)** Micrograph of an adult *C. elegans* on the first day of treatment (Day 0) and animals treated with Ctl, Lip-1 or SIH for eight days at 25 °C (Day 8). Scale bar = 50 µm. **(B)** Estimates of adult body length (in µm), showing SIH treated animals (blue) have longer average body length compared to age matched control (orange)or Lip-1 (green) treated populations (Kruskal-Wallis ANOVA: H(10)= 432.6, *p* < 0.0001; see **Supplemental Table S18** for sample summary and S19 for pair-wise comparisons). **Start** (grey) represents the beginning population of L4/young adults grown from egg at 25 °C for 48 hours prior to transfer to treatment plates. Each point represents an individual worm, with mean and error bars representing standard deviation (SD) **(C)** Estimated adult body volume (in pL), showing increased body volume with adult age for all groups (Kruskal-Wallis ANOVA: H(10)= 489, *p* < 0.0001; see **Supplemental Table S20** for sample summary and **Table S21** for pair-wise comparisons), with SIH treated animals having even greater body volume. Each point represents an individual worm, with mean ± SD. **(D)** Early fertility (first 24 hours) and total reproductive output are unaltered when vehicle control (Ctl) treated cohorts are compared to Lip-1 or SIH treated animals at 25 °C. Each data point represents an estimate from a single *C. elegans* adult, with mean ± SD (ANOVA: Early fertility F(2,74)=0.996, p=0.37; Total fertility F(2,57)= 4.89, *p*=0.011). **(E)** Estimates of maximum velocity achieved by aged and treated cohorts of *C. elegans*. Treatment with either Lip-1 or SIH attenuates the age-related decline in maximum velocity (Kruskal-Wallis ANOVA: H(7)= 298.5, *p* < 0.0001; see **Supplemental Table S22** for sample summary and **Table S23** for pair-wise comparisons). Each data point represents an estimate from a single *C. elegans* adult, with median ± interquartile range. Equivalent analyses of mean velocity and total distance travelled (and how these data correlate) are shown in **Supplemental Figures S12-14** and **Supplemental Tables S24-27**).

We also examined whether the interventions altered early and total reproductive output when worms were treated from early adulthood/late L4 (as used in the lifespan experiments). Early fertility (first 24 hours) was not altered by either SIH or Lip-1 treatment (**Figure 6D**; *p*>0.4). Lip-1 treatment resulted in a small decrease in lifetime reproductive output (**Figure 6D**; *p*<0.05), but SIH had no effect. Early fertility in *C. elegans* is paramount with respect to Darwinian fitness^39, 40^, so the reduction in lifetime fertility with Lip-1 treatment is consistent with a mild deleterious effect in early adulthood.

The effects of both interventions on movement parameters were assessed, since peak motile velocity has been previously demonstrated to correlate strongly with *C. elegans* healthspan and longevity^41^. As expected, control animals showed a steady decline in maximum velocity as they aged (**Figure 6E**). Treatment with SIH or Lip-1 markedly improved the maximum velocity of ageing animals (**Figure 6E**), with increases also in distance travelled and mean velocity (**Supplemental Tables S22-27** and **Figures S12-13**).

## Discussion

Our findings indicate that late-life dysregulation of glutathione and iron may trigger ferroptosis in *C. elegans* and that activation of this cell death signal limits lifespan. We previously determined that as *C. elegans* age, not only does intracellular iron accumulate but the capacity to safely sequester iron in the iron-storage protein ferritin fails. This contributes to the increased cellular fraction of Fe^2+^ that we observed here, leading to increased oxidative load^2, 16^, and providing the specific substrate for ferroptosis^42^. The observed depletion of glutathione during ageing further lowers the threshold for ferroptosis^1^.

Thus, in normal ageing, the decrease in GSH couples with an age-related increase in free Fe^2+^ to multiply the likelihood of ferroptosis, leading to death when combined changes in iron and GSH reach a critical threshold (**Figure 7**). We find that SIH and Lip-1 increase lifespan by prohibiting ferroptotic death rather than inhibiting ageing rate. We find also that late-life ferroptosis is a prominent contributor to age-related frailty. The healthspan benefits of inhibiting ferroptosis confirm that healthspan improvement need not always require a change in global ageing but can result from preventing a cause of frailty, raising an exciting conceptual prospect for therapeutic intervention.

**Figure 7.**
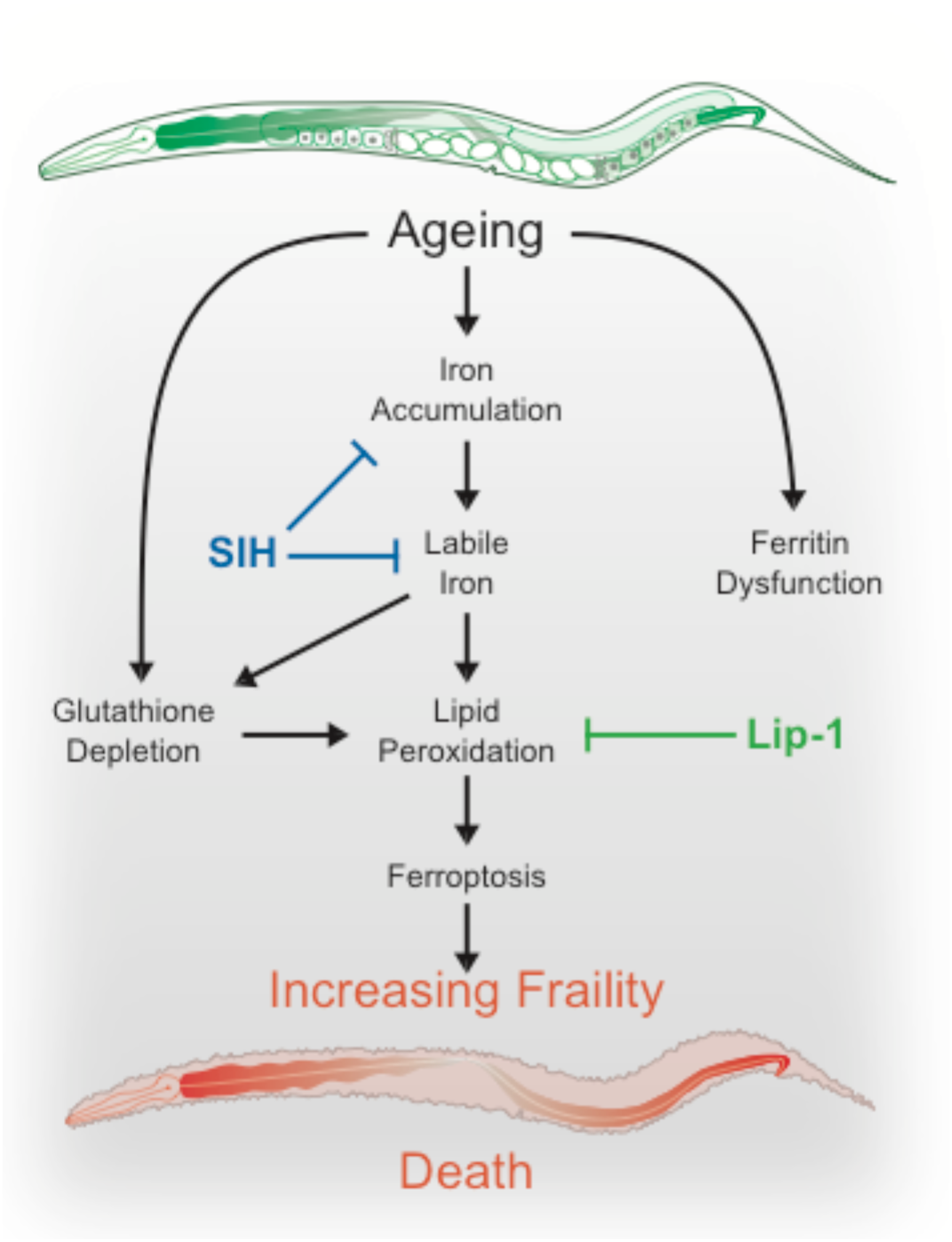
Schematic overview. During normal ageing iron unnecessary accumulates. The safe storage of surplus iron in ferritin begins to fail in late life, causing a corresponding elevation of reactive, ‘labile’ iron. In combination with falling glutathione levels there is increased risk of ferroptotic cell death, via lipid peroxidation signals. These cell death events increase frailty and ultimately shorten organism lifespan. These pharmacological interventions potentially represent targets to improve late life vigor and fitness.

Iron is critical to a growing organism, yet unnecessary retention might predispose towards increased frailty later in life^7, 43^. A previous study found that iron supplementation of *C. elegans* shortened lifespan but did not measure *in vivo* iron levels or the fraction of Fe^2+^ ^20^. Our measures of chronic versus acute changes in the fraction of Fe^2+^ help define the organismal limits of buffering capacity at 0.3 (or 30%). Iron deficiency causes deficits in major developmental pathways, but our interventions were delivered in adulthood to avoid any potential interference with development^7^. This revealed that limiting iron retention post-development was not only tolerable, but improved health and life history traits.

SIH and Lip-1 alter ageing at specific life phases and, unlike most known lifespan interventions, do not slow the rate of ageing ^37^. They act in related but different ways to extend lifespan, with differing impacts on hazard rate and distinct effects on iron levels, life-history traits and acute glutathione depletion. Importantly, both interventions increase both lifespan and healthspan without apparent major fitness trade-offs, in contrast to those previously reported in long-lived mutants that do slow the rate of ageing^39, 40^.

Ageing is the principal risk factor for many major human diseases including cancer and dementia. An ancient biochemical dependence on iron may have established an inevitable liability in late life. Needless iron elevation in somatic tissue has been described in many organisms from drosophila to rodents to humans, particularly in brain^3–5^ and might be a universal feature of ageing. Notably, there is no excretion mechanism for systemic iron in animals^44, 45^, highlighting that while iron is limiting for development, there has been no evolutionary pressure to regulate its accumulation in post-reproductive life. Ferroptosis plays an important role in kerbing cancer but becomes inappropriately activated in ischemia and neurodegeneration, where its inhibition holds therapeutic promise^8–11^. Future studies could test the hypothesis that the cancer-protective benefit of ferroptosis involves the reciprocal acceleration of ageing.

## Acknowledgments

We thank Abdel Belaidi for comments on the manuscript, Nicholas Stroustrup and Walter Fontana for providing their raw data to enable validation of our temporal scaling analysis and acknowledge the Australian Synchrotron. This study was supported by grants from the Australian Research Council to AIB and GM (DP130100357 and DP180101248), University of Melbourne Research Grant Support Scheme and Miller Foundation to GM, and the Victorian Government’s Operational Infrastructure Support Program. We thank the Caenorhabditis Genetics Center (CGC) supported by the US National Institutes of Health - Office of Research Infrastructure Programs (P40 OD010440) for providing *C. elegans* strains.

## Author Contributions

NLJ, AIB and GM planned the study. NLJ and GM performed the experiments with assistance from SAJ and FS. NLJ, SAJ, AS, TPS and GM analysed the data. DRR and MC provided resources. NLJ, AIB and GM wrote the manuscript with contributions from all authors.

## METHODS

### Strains

Wild type (strain N2) and the temperature sensitive-sterile strain TJ1060: *spe-9*(*hc88*); *fer-15*(*b26*) were obtained from the *Caenorhabditis* Genetics Center. The wild type strain was maintained at 20 °C on standard nematode growth media (NGM) ^47^ and aged at 20 °C or 25 °C as required. TJ1060 was maintained at 16 °C and also aged at 20 °C or 25 °C as required. TJ1060 was predominately used to remove the inconvenience of progeny production and can be regarded as a proxy for wild type.

### Compounds

Compounds used in this study include:

- Diethyl maleate (DEM) obtained from Sigma-Aldrich.

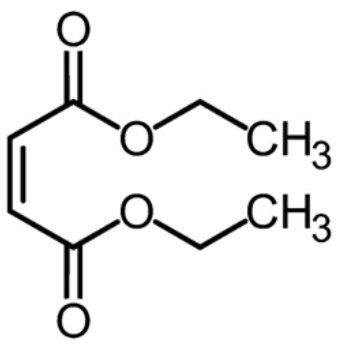
- Liproxstatin (Lip-1; N-[(3-chlorophenyl) methyl]-spiro[piperidine-4,2’(1’H)-quinoxalin]-3’- amine) obtained from the laboratory of Marcus Conrad (initially) and subsequently ApexBio Tech LLC.

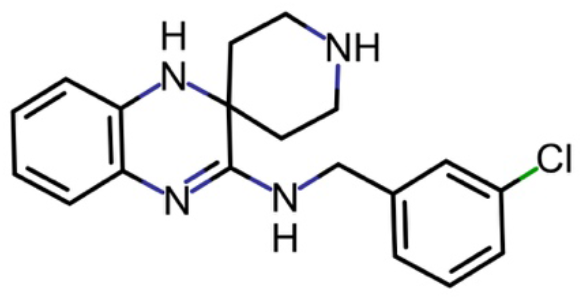
- Salicylaldehyde isonicotinoyl hydrazone (SIH) obtained from the laboratory of Des Richardson (University of Sydney).

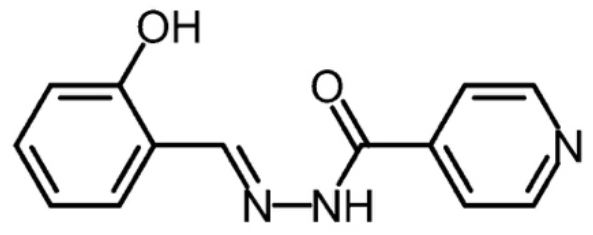
- SIH precomplexed with iron as Fe(SIH)_2_NO_3_.

### Glutathione depletion

Diethyl maleate (DEM; Sigma-Aldrich) was added to neat DMSO and added to molten NGM at 55 °C to a final concentration of 5, 10, 15 or 20 mM DEM and 0.5 % v/v DMSO. Plates were seeded with OP50 and used within 24 hours. As above, data was collected at 25 (±1) °C using the temperature sensitive-sterile strain TJ1060. A synchronous population was obtained by transferring egg-laying adults to fresh plates at 16 °C for 2-3 hours. The adults were removed and the plates with eggs then transferred to 25 °C to ensure sterility. After 48 hours at 25 °C, when worms were at the late L4/young adult stage, 25-35 nematodes were transferred to fresh plates containing either vehicle control, 250 µM SIH, or 200 µM Lip-1. Worms were aged at 25 °C for a further 4 days and then transferred to DEM plates. Survival, determined by touch-provoked movement, was scored at 24 and 48 hours after exposure to DEM.

Ageing studies were also undertaken to determine changes with age of both survival after DEM exposure and basal glutathione levels. Initial populations were obtained as describe above, with worms aged on standard NGA plates.

### Quantification of total glutathione

Measurement of total glutathione per worm was based on established protocols and is based on a kinetic spectrophotometric assay using the reaction between GSH and 5,5′-dithio-bis (2-nitrobenzoic acid) (DTNB) measured at 412 nm ^48, 49^. All reagents were freshly prepared prior to the assay and for each estimate 50 adults were collected in 200 µL of S-basal ^47^ in 1.7 ml microfuge tubes. Animals were washed twice in S-basal, pelleted via centrifugation and total volume reduced to 20 µL. A 50 µL aliquot of Extraction Buffer was added, then the samples were frozen in Liquid N_2_ and store at −80 °C until required. Extraction buffer consisted of 6 mg/mL 5-sulfosalicylic acid dehydrate, 0.1 % v/v Triton X-100 and Complete, EDTA-free Proteinase inhibitor cocktail (Roche) in KPE buffer (0.1 M potassium phosphate buffer and 5 mM EDTA at pH 7.5).

Samples were homogenized with a Bioruptor Next Gen (Diagenode) bath sonicator, set on HIGH and cooled to 4 °C, using 10 cycles of 10 seconds ON and 10 seconds OFF. Supernatant was collected following a 14K x *g* spin at 4 °C. Assays were performed in 96 well microplates (clear polystyrene, flat-bottomed, Greiner bio-one), in a total volume of 200 µL per well. To each well was added 50 µL of lysate supernatant, 50 µL of milli-Q H_2_O and then 100 µL of GA buffer (NADPH 400 µM, glutathione reductase 1 U/mL and 0.3 mM DTNB in KPE buffer diluent). Reactions were incubated for 1-2 min at room temperature and then absorbance measured at 412 nm for 10 min with 1 min interval using a Powerwave plate spectrophotometer (BioTek). The rate of change in absorbance per minute is linearly proportional to the total concentration of GSH. Total GSH in the samples was interpolated from using linear regression from a standard curve of known GSH concentrations (0 to 1 µM) run in tandem. Within experiment results are presented as relative glutathione levels, where results are normalized to the mean of the starting population.

### Lipid peroxidation

Measurement of malondialdehyde (MDA) was performed using a Thiobarbituric acid reactive substances (TBARS) assay kit (10009055, Caymen Chemical) as per manufacturer instructions using reduced reaction volumes of 1 mL. For *C. elegans* samples with acute glutathione depletion, Day 1 adults were treated with and without 20 mM DEM for 6h at 25 °C prior to collection. For ageing, animals were aged at 25 °C and treated with Lip-1 or SIH as previously described. Replicate samples were collect, washed twice in S-basal, pelleted by centrifugation. Following removal of excess buffer samples (∼40 µL) were frozen in liquid-N_2_ and stored at −80 °C until needed. Samples were then homogenized via a Bioruptor bath sonicator (Diagenode, set on ‘high power’ with 10 cycles of 10s pulses with a 10s pause between pulses, at 4 °C), then centrifuged at 21,500 x*g* at 4 °C for 30 min and the supernatant retained. The concentration of protein was determined using a BCA assay kit (Bio-Rad) and equivalent aliquots of 20-25 µg total protein used for subsequent measurements.

Analysis of Hydroxynonenal (4-HNE) protein adducts was also used as a proxy for lipid peroxidation. Duplicate samples of 50 and 200 worms were collected and washed twice in S-basal, pelleted by centrifugation and the supernatant discarded. These samples (∼30 µL) were frozen in liquid-N_2_ and stored at −80 °C until needed. To each sample an 10 µL 4x Bolt LDS sample buffer (Invitrogen) and 3 µL TCEP (Invitrogen) was added and the sample heated to 95 °C for 10 min. Lysates were loaded onto NuPAGE™ 4-12% Bis-Tris acrylamide gels (1.0 mm, 10-well, Invitrogen), electrophoresed with MES running buffer and then transferred onto 0.45 µm PVDF membrane by electroblot using a Mini Blot module (Invitrogen). 4-HNE protein adducts were detected by an anti 4-HNE protein adduct antibody (1:2000, AB5605, Millipore) in Tris-buffer saline with 5% skim milk, and ECL (GE Healthcare). The membranes were stripped using a 1x ReBlot Strong Antibody Stripping Solution (Merck) for 15 min, reprobed for tubulin using an anti-Tubulin antibody (1:10,000, T6074, Sigma-Aldrich).

### Visualization of cell death

The red-fluorescent propidium iodide (PI), was used to visualize dead cells within live *C. elegans* after DEM treatment and during ageing. Populations were incubated for 24 h at 25 °C with PI (a 10 µL volume of 0.25 mg/mL solution added to the bacterial lawn on 50mm NGM plates) prior to the described age or with concurrent exposure to 10 mM DEM (as described above) and PI. For ageing experiments, animals were visualized at Day 6 and Day 8. Cohorts of live animals (*i.e.* showing spontaneous or touch-provoked movement) were isolated and mounted under glass coverslips on 2% agarose pads without anesthetic. Imaging were captures with on a Leica DMI3000B inverted microscope, DsRed filter set and a DFC 3000G digital.

### Liquid chromatography-inductively coupled plasma mass spectrometry

Liquid chromatography was performed using established protocols^2^. Briefly, samples of aged *C. elegans* were lysed using a Bioruptor Next Gen (Diagenode) bath sonicator set on HIGH and cooled to 4 °C using 10 cycles of 10 sec ON and 10 sec OFF, in a 1:1 volume ratio of Tris-buffered saline (pH 8.0) with added proteinase inhibitors (EDTA-free; Roche). Sample homogenization was confirmed by microscopic inspection. Lysates were then centrifuged for 15 min at 175,000 *g* at 4 °C. The supernatant was removed and total protein concentration in the soluble fraction was determined using a NanoDrop UV spectrometer (Thermo Fisher Scientific) before being transferred to standard chromatography vials with polypropylene inserts (Agilent Technologies) and kept at 4 °C on a Peltier cooler for analysis. Size exclusion chromatography-inductively coupled plasma-mass spectrometry was performed using an Agilent Technologies 1100 Series liquid chromatography system with a BioSEC 5 SEC column (5 μm particle size, 300 Å pore size, I.D. 4.6 mm, Agilent Technologies) and 7700x Series ICP-MS as previously described^50^. A buffer of 200 mM NH_4_NO_3_ was used for all separations at a flow rate of 0.4 mL min^−1^. A total of 50 µg of soluble protein was loaded onto the column by manually adjusting the injection volume for each sample. Mass-to-charge ratios (*m/z*) for phosphorus (31) and iron (56) were monitored in time resolved analysis mode.

Plots of the mean (± standard deviation) of three independent biological replicates are shown. Integration of the three major peaks was performed using Prism (ver. 7 for Mac OS X, Graphpad).

### X-ray Fluorescence Microscopy

#### Sample preparation - Elemental mapping

Specimens were prepared for XFM using previously described protocols^51, 52^. Briefly, adult *C. elegans* were removed from NGM, washed four times in excess S-basal (0.1 M NaCl; 0.05 M KHPO_4_ at pH 6.0), briefly in ice-cold 18 MΩ resistant de-ionized H_2_O (Millipore) and twice in ice-cold CH_3_COONH_4_ (1.5 % w/v). Samples were transferred onto 0.5 µm-thick silicon nitride (Si_3_N_4_) window (Silson), excess buffer wicked away and then the slide was frozen in liquid nitrogen (N_2_)-chilled liquid propane using a KF-80 plunge freezer (Leica Microsystems). The samples were lyophilised overnight at −40 °C and stored under low vacuum until required.

#### Elemental mapping

The distribution of metals was mapped at the X-ray Fluorescence Microscopy beamline at the Australian Synchrotron ^53^ using the Maia detector system ^54^. The distribution of elements with atomic number < 37 were mapped using an incident beam of 15.6 keV X-rays. This incident energy allowed clear separation of X-ray fluorescence (XRF) peaks from the relatively intense elastic and inelastic scatter. The incident beam (∼1.71 ×10^9^ photons s^−1^) was focussed to approximately 2 × 2 μm^2^(H × V, FWHM) in the sample plane and the specimen was continuously scanned through focus (1 mm sec^−1^). The resulting XRF was binned in 0.8 μm intervals in both the horizontal and vertical giving virtual pixels spanning 0.64 μm^2^ of the specimen probed with a dwell time of 8 μsec. XRF intensity was normalized to the incident beam flux monitored with a nitrogen filled ionization chamber with a 27 cm path length placed upstream of the focusing optics. Three single-element thin metal foils of known areal density (Mn 18.9 μg cm^−2^, Fe 50.1 μg cm^−2^ and Pt 42.2 μg cm^−2^, Micromatter, Canada) were used to calibrate the relationship between fluorescence flux at the detector and elemental abundance. Dynamic Analysis, as implemented in GeoPIXE 7.3 (CSIRO), was used to deconvolve the full XRF spectra at each pixel in the scan region to produce quantitative elemental maps ^55^. This procedure includes a correction for an assumed specimen composition and thickness, in this case 30 μm of cellulose. Though unlikely to exactly match the actual sample characteristics, deviations from these assumptions are not significant for the results presented in this study as the effects of beam attenuation and self-absorption on calcium and iron XRF are negligible for a dried specimen of this type and size ^56^.

#### Elemental quantification and image analysis

Analysis of elemental XRF maps was performed using a combination of tools native to GeoPIXE and ImageJ ^57^. Incident photons inelastically scattered (Compton scatter) from the sample detail the extent and internal structure of individual *C. elegans*. The differential scattering power of the specimens and substrate allowed individual animals (or parts thereof) to be identified as regions of interest (ROI; **Supplemental Figure S1**) facilitating analysis of elemental content on a ‘per worm’ basis. This segmentation of each elemental map was achieved using the histogram of pixel intensities from Compton maps to locate the clusters within the image. ROIs composed of < 10,000 pixels were deemed to be so small that their elemental content was not reflective of the elemental content of whole animals and so these were excluded from the analysis. The ‘non-worm’ region of each scan was used to calculate the value specimen elemental content was distinguishable from background noise, *i.e.* the critical value as defined by ^58^. The background corrected elemental maps were used to establish the areal densities and the total mass of each element associated with individual ROIs.

#### Sample preparation - *φ*XANES Imaging

Adult *C. elegans* were removed from NGM, washed four times in excess ice-cold S-basal (0.1 M NaCl; 0.05 M KHPO_4_ at pH 6.0). Samples were transferred onto 0.5 µm-thick silicon nitride (Si_3_N_4_) window (Silson), excess buffer wicked away and then the slide was frozen *in situ* under a laminar stream of 100 °K dry nitrogen (N_2_) gas.

#### *φ*XANES Imaging

The beam energy was selected using a Si(311) double-crystal monochromator with a resolution of ∼0.5 eV. *φ*XANES imaging was achieved by recording Fe XRF at 106 incident energies spanning the Fe K-edge (7112 eV). Measurement energy interval was commensurate with anticipated structure in the XANES: 

|  |  |  |  |  |
| --- | --- | --- | --- | --- |
| 7000 eV to 7100 eV: | 5 | × | 20.0 eV | steps |
| 7100 eV to 7105 eV: | 5 | × | 1.0 eV | steps |
| 7105 eV to 7135 eV: | 75 | × | 0.4 eV | steps |
| 7135 eV to 7165 eV: | 15 | × | 2.0 eV | steps |
| 7165 eV to 7405 eV: | 1 | × | 240.0 eV | steps |
| 7405 eV to 7455 eV: | 5 | × | 5.0 eV | steps |

As for XFM, *φ*XANES measurements used a beam spot ∼2 × 2 μm but data was recorded using continuous scanning at 0.2 mm sec^−1^ (binned at 2 μm intervals). Transit time through each virtual pixel was 10 ms and the incident X-ray intensity at 7455 eV was ∼1.67 ×10^10^ photons s^−1^. These imaging parameters gave a total dose associated with the *φ*XANES measurement estimated at ∼5 MGy. This value is commensurate with doses typically delivered during bulk X-ray absorption spectroscopy.

#### *φ*XANES analysis

The XANES spectra from an iron foil (50.1 μg cm^−2^, Micromatter Canada) was measured to monitor the energy calibration of the beamline. The maxima of the first peak in the derivative spectra of the iron foil was subsequently defined as 7112.0 eV. The energy stability of beamline has been determined at < 0.25 eV over 24 hrs making energy drift over the course of a scan negligible. Consistency of the measured edge positions in conjunction with stability of beam position and flux recorded in ion chambers upstream the specimen position provide confidence that energy stability was high through the duration of the experiment. Small position drifts were aligned by cross-correlation of the calcium map which remains essentially constant throughout the energy series.

XANES probes the density of states on the absorbing atom and reveals electronic and structural details of coordination environment. The aligned *φ*XANES image series is stack of images, one per incident energy allowing the XANES of individual cells to be assessed. Previous work has shown that the distribution of calcium is a useful marker for the position of *C. elegans* intestinal cells and we used this information to identify regions of interest in the *φ*XANES stack corresponding to anterior intestinal cells. Anterior intestinal cells were chosen due to their consistent and robust iron content^51^; see **Figure S2** for the ROIs and iron maps used for the *φ*XANES analysis.

As all points on the specimen represent a heterogenous mixture of iron binding species the resulting XANES spectra are admixtures with contributions from all of these components. The technical particulars of the XFM beamline (being primarily designed for elemental mapping) are not optimised for high resolution spectroscopy and our XANES spectra are relatively sparse. For iron K-edge XANES the abrupt increase in absorption coefficient at the critical threshold obscures the presence of 1s → 4s and 1s → 4p electronic transitions. Berry *et al* ^59^ demonstrated that the relative intensity of these transitions provides the proportional contribution of Fe^2+^ and Fe^3+^ to the XANES and can be assessed by interrogating the first derivative of the XANES spectra.

#### Lifespan determination

Lifespan was measured using established protocols ^2,60^. SIH was dissolved in neat dimethyl sulfoxide (DMSO; Sigma-Aldrich) then added to the molten NGM at 55 °C (to a final concentration of 250 µM SIH in 0.5 % v/v DMSO). Lip-1 was dissolved in neat DMSO then added to the molten NGM at 55 °C (to a final concentration of 200 µM Lip-1 in 0.5 % v/v DMSO). Media containing equivalent vehicle alone (0.5 % v/v DMSO) was used for comparison. Standard overnight culture of the *Escherichia coli* (*E. coli*) strain OP50 was used as the food source.

Lifespan data was collected at 25 (±1) °C using the temperature sensitive-sterile strain TJ1060 [*spe-9*(*hc88*); *fer-15*(*b26*)]. A synchronous population was obtained by transferring egg-laying adults to fresh plates at 16 °C for 2-3 hours. The adults were removed and the plates with eggs then transferred to 25 °C to ensure sterility. After 48 hours at 25 °C, when worms were at the late L4/young adult stage, 25-35 nematodes were transferred to fresh plates containing either vehicle control, 250 µM SIH, or 200 µM Lip-1. All plates were coded to allowing blinding of the experimenter to the treatment regime during scoring. Nematodes were scored for survival at one to three-day intervals and transferred to freshly prepared plates as needed (2-5 days).

*C. elegans* are bacteriophores and the *E. coli* (OP50) monoxenic diet can colonize the pharynx and intestine, resulting in death. Consequently, antibiotics are known to extend *C. elegans* lifespan ^61^. In addition, iron chelating compounds, such as EDTA have been reported to have antibiotic properties. We performed a *disk diffusion test* ^62^ on both Lip-1 and SIH and observed no evidence for inhibition of *E. coli* (strain OP50) growth). Furthermore, an additive effect on median lifespan extension was seen when SIH and the antibiotic ampicillin were co-administered to *C. elegans* (**Supplemental Figure S15**), consistent with independent effects on lifespan.

It is well documented that differences are observed between independent measures of lifespan, with micro-environmental factors such as minor temperature fluctuations potentially resulting differences in median and maximum lifespan between replicates ^63^. After determining the optimal doses of 250 µM SIH and 200 µM Lip-1, respectively, cohorts of nematodes were compared in 8 independent replicates. As the number of worms measured is known to influence the likelihood of accurately observing differences in lifespan ^64^, the starting populations for all treatments within experiments were in excess of 70 individuals. The median and maximum lifespans observed of control and treated populations for these 8 replicates are shown in **Table S8.** As can be seen in this table, the median lifespan of treated populations was always greater than that of control populations, however the magnitude of the difference varied between experiments, with the median lifespan of control populations ranging from 7 to 9 days.

#### Body size analysis

A developmentally synchronous population, derived from eggs laid over a 2-hour window, were cultured on NGA media at 25 °C for 48 h, and then as young adult worms were transferred onto three treatment plates for an additional 24 h. The treatment plates included NGA with 0.5 % (v/v) DMSO (vehicle control, Ctl), 250 µM SIH, or 200 µM Lip-1 (as described above).

Cohorts of approximately 100 animals were transferred into a 1.5 ml centrifuge tube containing 400 µL S-basal. Following a brief centrifugation excess S-basal was removed leaving the animals suspended in 50 µL. Animals were euthanised and straightened by a 15 second exposure to 60 °C (using a heated water bath). Samples were then mounted between glass slides and a cover slip and immediately imaged. Micrographs were collected using a Leica M80 stereomicroscope and Leica DFC290 HD 3 MP) digital camera. Pixel sizes were defined using a calibrated 25 µm grid slide (Microbrightfield, Inc). Size and shape metrics were extracted from brightfield images were analysed using the WormSizer plugin ^65^ for ImageJ.

#### Fertility analysis

Wild type (N2) adults (4-day post egg lay) were transferred to fresh plates for 30 minutes at 20 °C to establish a developmentally synchronous population. Adult nematodes were then removed, and eggs were then transferred to 25 °C. As with the survival analyses, after 48 hours at 25 °C, when worms were at the late L4/young adult stage individual nematodes were transferred to plates containing vehicle control, 250 µM SIH, or 200 µM Lip-1. After 24 hours, adult worms were transferred to fresh plates and transferred daily until the end of the fertile period. After allowing progeny to develop for 2 days at 20 °C, they were then counted to determine daily and total fertility. Early fertility is determined by the number of progeny laid in the first 24-hour period.

#### Movement

A developmentally synchronous population, derived from eggs laid over a 2-hour window, were cultured on NGA media at 25 °C for 48 h, and then as young adult worms were transferred onto three treatment plates for an additional 24h. The treatment plates included NGM + 0.5 % (v/v) DMSO (vehicle control, Ctl), NGM + 250 µM SIH, and NGM + 200 µM Lip-1 (as described above).

Single worms were transferred to a 55 mm NGA assay plate devoid of a bacterial lawn, without a lid, and left to recover from the transfer for 2 minutes. Movement of the adults was then recorded using a stereomicroscope (Leica M80) with transmitted illumination from below. A 30 second video recording was captured using a 3 MP DFC290 HD digital camera (Leica Microsystems) at a rate of 30 frames per second. Pixel length was calibrated using a 25 µm grid slide (Microbrightfield, Inc). Recorded series were analysed using the wrMTrck plugin^66^ for ImageJ (www.phage.dk/plugins) and Fiji ^67^ (a distribution of ImageJ).

The maximum velocity achieved was expressed as mm per second (as derived from the distance between displaced centroids per second). Additional metrics of movement were determined including mean velocity (mm s^−1^) and (total) distance travelled (mm). These variables were collated in Prism (v7.0a GraphPad Software) and presented as a scatter plot with medians and interquartile range.

### Quantification and Statistical Analysis

#### Areal density and total iron analysis

Areal iron and total body iron data were assessed for normality using a D’Agostino & Pearson test (see **Supplemental Tables S3 and S5**). Based on this analysis a one-way ANOVA was performed followed by a Sidak’s multiple comparisons test (as implemented by PRISM; see **Supplemental Tables S4 and S6**).

#### Standard lifespan analysis

Kaplan–Meier survival curves were generated and compared via non-parametric log rank tests (Prism v7.0a, GraphPad Software).

#### Testing for Departure from Temporal Rescaling

Following the recently published results of Stroustrup *et al*. we determined whether the results observed with both the SIH and Lip-1 interventions were due to temporal scaling of ageing. A modified Kolmogorov-Smirnov (K-S) test was applied to **the residuals from a replicate-specific accelerated failure time (AFT) model fitted according to the Buckley-James method that uses a nonparametric baseline hazard function**. The function *bj* in R package *rms* was used to fit the replicate-specific model with interventions as categorical independent variables. We used the same approach for testing whether the temperature difference results in simple temporal rescaling, with the only difference being using temperature rather than intervention as categorical independent variable in the AFT model. Full details of these analyses are included in the online Supplementary information.

#### Characterizing Departure from Temporal Rescaling

Parametric survival models with Weibull baseline hazards and Gamma frailty were fitted to replicate-specific data using the R package *flexsurv*. A likelihood ratio test was used to compare models that assume simple temporal rescaling to models that allow varying degrees of departure from temporal rescaling. The best model for each replicate was selected using a likelihood ratio test and the goodness of fit (GOF) of the best model is evaluated using a chi-square GOF test. To combine data across different replicates, we performed fixed-effect and random-effect meta-analyses for each parameter in the best model (**Supplemental Table S13)**. Briefly, the fixed-effect meta-analysis estimates were derived using Inverse Variance Weighting (IVW) in which the estimates from each replicate were weighted by the inverse of their variance estimates. The meta-analysis estimates were then calculated simply as the weighted average of estimates from all replicates. The fixed-effect meta-analysis assumes that there is insignificant variation between the estimates of the same parameter across different replicates. The random-effect meta-analysis also derives the estimates by assigning weights to estimates from each replicate, but in this case the weights take into account the variation of estimates across replicates.

The fixed-effects and random-effects meta-analysis estimates are quite similar and in **Supplemental Figure S9** we can see that the meta-analysis estimates provide the best fit to SIH data and worst for Lip-1 data. Since there is significant between-replicate variation for the majority of the parameters, it is not surprising that the when the meta-analysis estimates are applied to the real data, a chi-square goodness of fit reveals significant lack of fit (*χ*^2^(3) = 237.0 for control worms, *χ*^2^(5) = 258.0 for Lip-1 and *χ*^2^(3) = for SIH, all p-values < 0.001).

One notable pattern from **Table S13** is that for nearly all replicates, there is more heterogeneity due to unobserved factors among the control worms, as indicated by the negative Dlog(s^2^) parameter estimates for Lip-1 and SIH data. This heterogeneity is also reflected in a de-acceleration of the hazard function for control worms (**Supplemental Figures S9c-d**) beyond 7-8 days. This de-acceleration of the hazard function is the main contributor to the crossing behaviour we observe when comparing the survival functions (**Supplemental Figure S10**), and it is what causes a violation of the simple temporal rescaling assumption.

#### Survival during GSH depletion

For survival with increasing DEM dose response and protection by compounds (Lip-1 and SIH), data was plotted as fraction of animal alive with upper and lower 95% confidence interval, using the Wilson ‘score’ method ^68^ using asymptotic variance ^46^ and fitted with a sigmoidal curve (Prism). Pairwise comparisons of treated groups versus control at each concentration of DEM was determined using the N-1 chi-squared test ^69, 70^.

#### Fertility

Differences in fertility (*i.e.* early and total reproductive output) were assessed using an ordinary one-way analysis of variance (ANOVA), followed by a Tukey’s multiple comparison test (as implemented by Prism v7.0a, GraphPad Software).

#### Body length and volume analysis

Data of estimated adult body length and volume were initially assessed for normality using a D’Agostino & Pearson test (see **Supplemental Tables S18 and S20**). Based on this analysis a nonparametric Kruskal-Wallis Analysis of Variance (ANOVA) was performed followed by a Dunn–Šidák test for multiple comparisons (as implemented by Prism v7.0a, GraphPad Software; **Supplemental Tables S19 and S21**).

#### Movement Analysis

Data of estimated maximum velocity were initially assessed for normality (see **Supplemental Table S22**). Based on this analysis a nonparametric Kruskal-Wallis ANOVA was performed followed by a Dunn–Šidák test for multiple comparisons (as implemented by PRISM; **Supplemental Table S23**). Mean velocity and total distance travelled were also determined (**Supplemental Figures S12-13**). Results summaries and comparisons between treatments are shown in **Supplemental Tables S24-27**. The data for the three movement parameters were combined across treatments and ages to determine the relationship between the estimated parameters, all were found to be positively correlated (**Supplemental Figure S14**).

#### Cell death analysis

Differences between the proportion of live animals with fluorescently labelled nuclei in control versus Lip-1 and SIH treatment, either aged or exposed to DEM, were compared using a z-test.

#### Type I error for statistical hypothesis testing

Unless otherwise stated, all statistical tests are conducted with type I error set at 0.05.

## SUPPLEMENTARY INFORMATION

### Glutathione determination

#### Glutathione and aging

There was a significant reduction in glutathione levels with increased adult age in *C. elegans* (**Figure 1c**). For comparisons between age and treatment groups an Ordinary one-way ANOVA was performed, followed by Tukey’s multiple comparisons test (ANOVA: F (3, 20) = 32.96, *p*< 0.0001). The results of the pairwise comparisons, corrected for multiple comparisons, are shown in **Table S1**.

**Table S1:** Summary of glutathione level comparisons between ages

| Tukey's multiple comparisons test | Mean Diff. | 95.00% CI of diff. | Significant? | Adjusted *p* Value |
| --- | --- | --- | --- | --- |
| Day 1 vs. Day 4 | 0.1964 | 0.08439 to 0.3085 | Yes | 0.0003 |
| Day1 vs. Day 8 | 0.2738 | 0.1618 to 0.3858 | Yes | <0.0001 |
| Day 1 vs. Day 10 | 0.3667 | 0.2546 to 0.4787 | Yes | <0.0001 |
| Day 4 vs. Day 8 | 0.07738 | -0.03466 to 0.1894 | No | 0.2992 |
| Day 4 vs. Day 10 | 0.1702 | 0.0582 to 0.2823 | Yes | 0.0015 |
| Day 8 vs. Day 10 | 0.09286 | -0.01918 to 0.2049 | No | 0.1423 |

#### Glutathione depletion

There was a significant reduction in glutathione levels after treatment with of 4 Day old adults with DEM (**Figure 1f**). Pre-treatment with SIH protected against this reduction, with this treatment also resulting in a higher basal level of glutathione. For comparisons between treatment groups an Ordinary one-way ANOVA was performed, followed by Tukey’s multiple comparisons test (ANOVA: F (5, 30)= 50.97, *p*< 0.0001). The results of the pairwise comparisons, corrected for multiple comparisons, are shown in **Table S2.**

**Table S2:** Summary of glutathione level comparisons after DEM exposure with pre-treatment

| Tukey's multiple comparisons test | Mean Diff. | 95 % CI of diff. | Significant ? | Adjusted *p* Value |
| --- | --- | --- | --- | --- |
| Day 4 Control vs Day 4 Lip-1 | -0.04508 | -0.3588 to 0.2686 | No | 0.9997 |
| Day 4 Control vs Day 4 SIH | -0.6447 | -0.9584 to -0.331 | Yes | <0.0001 |
| Day 4 Ctl vs. Day 4 Ctl + DEM | 0.6921 | 0.409 to 0.9752 | Yes | <0.0001 |
| Day 4 Lip-1 vs. Day 4 Lip-1 + DEM | 0.7379 | 0.4515 to 1.024 | Yes | <0.0001 |
| Day 4 SIH vs. Day 4 SIH + DEM | 0.7667 | 0.4857 to 1.048 | Yes | <0.0001 |
| Day 4 Ctl + DEM vs. Day 4 Lip-1 + DEM | 0.0006989 | -0.2518 to 0.2532 | No | >0.9999 |
| Day 4 Ctl + DEM vs. Day 4 Lip-1 + DEM | -0.5701 | -0.8166 to -0.3237 | Yes | <0.0001 |

### Supplemental Analysis of X-ray Fluorescence Mapping

**Figure S1:** The following figures show the ROIs associated with each iron map (ROI; white = overlapped/fractional, green = whole/minimally overlapped, red = excluded from analysis). Shown are the masks used to identify and analyse the iron elemental maps of TJ1060 populations at different adult ages ± Lip-1 or SIH at 25 °C.

Figure S1A: Day 1 adults (i.e. first day of adulthood)
Figure S1B: Day 4 Control
Figure S1C: Day 4 + Lip-1
Figure S1D: Day 4 + SIH
Figure S1E: Day 8 Control
Figure S1F: Day 8 + Lip-1
Figure S1G: Day 8 SIH

**Figure S1A.**
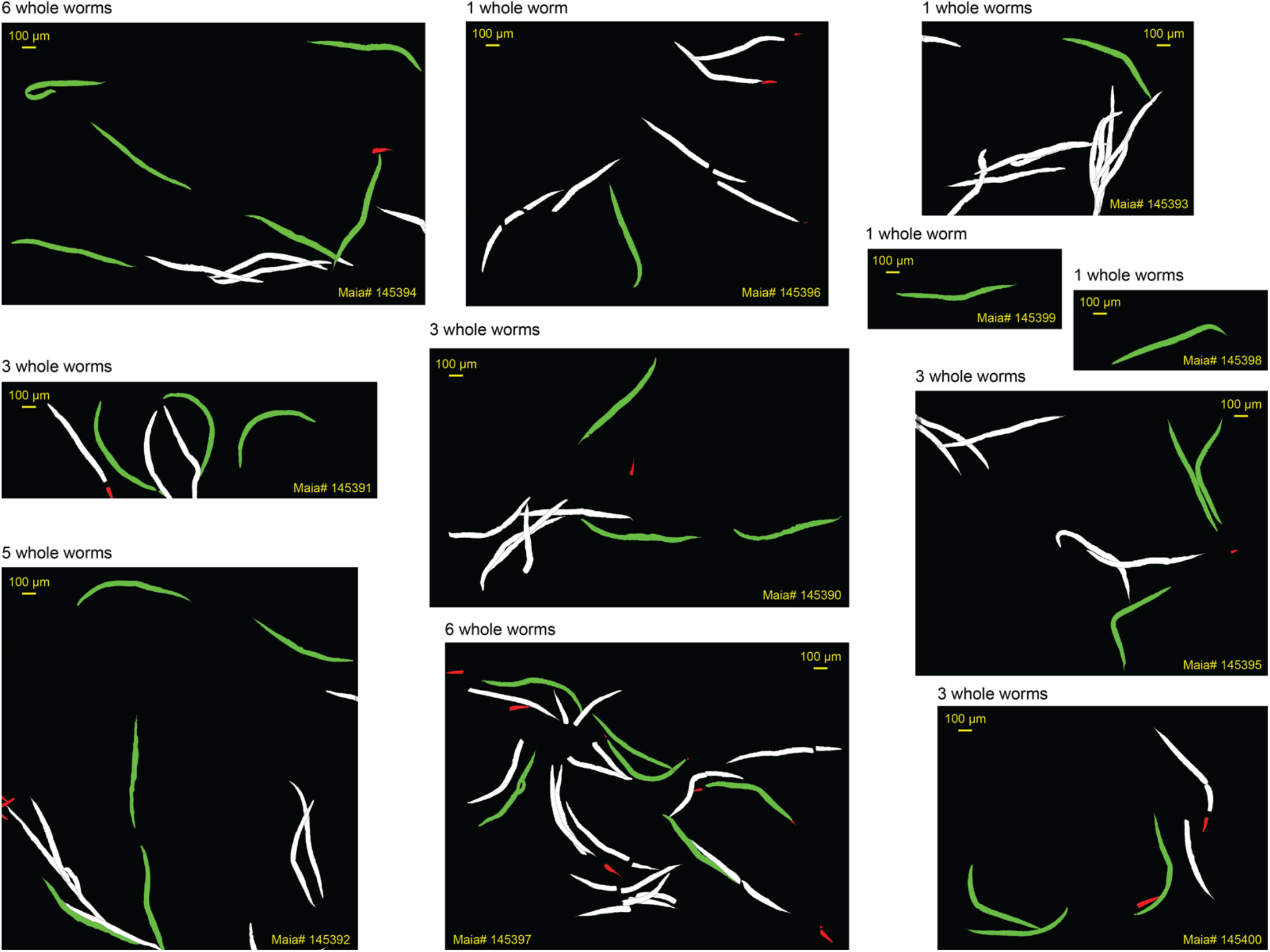
Masks for 1 day old adults (starting population)

**Figure S1B.**
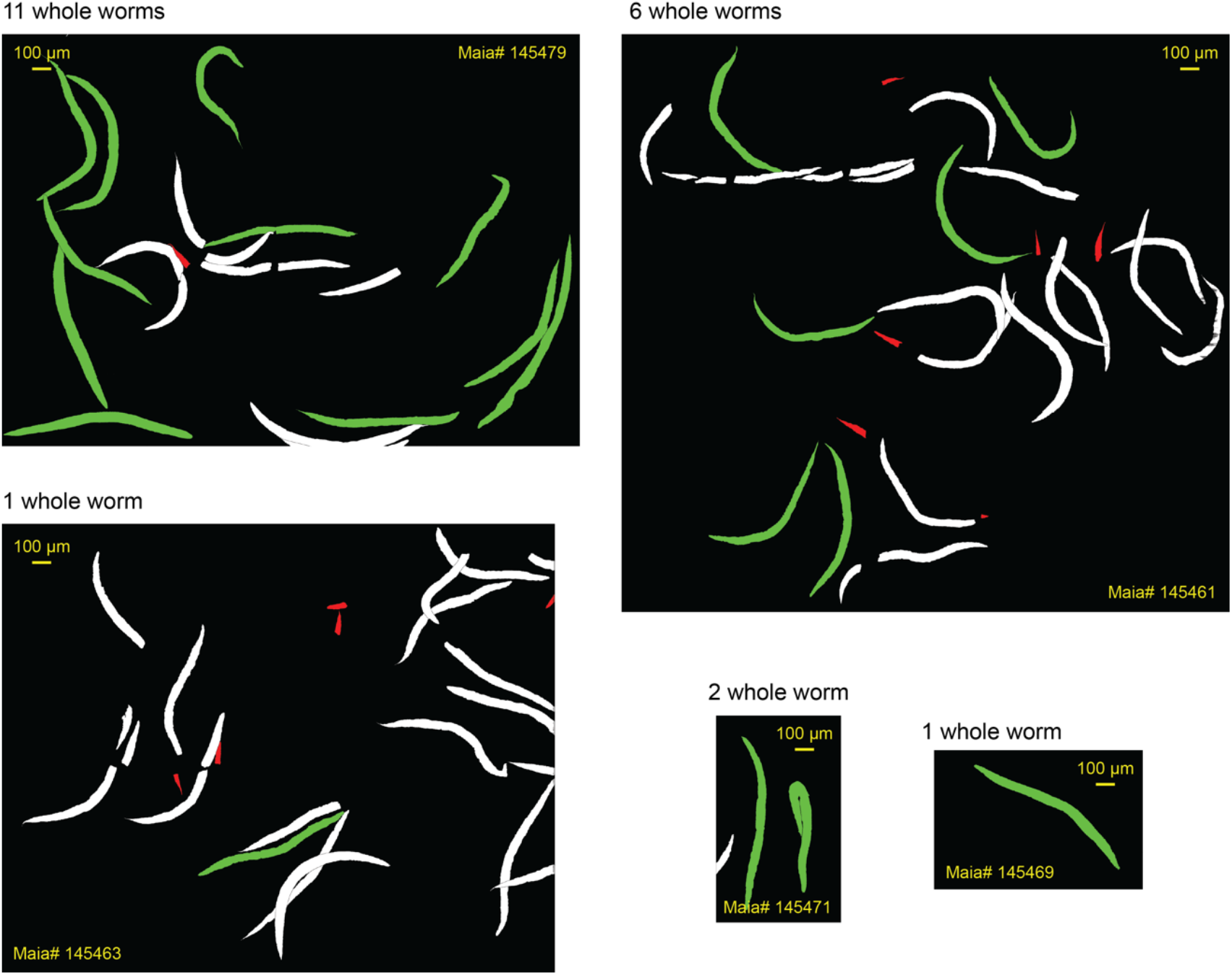
Masks for 4 day old Control adults.

**Figure S1C.**
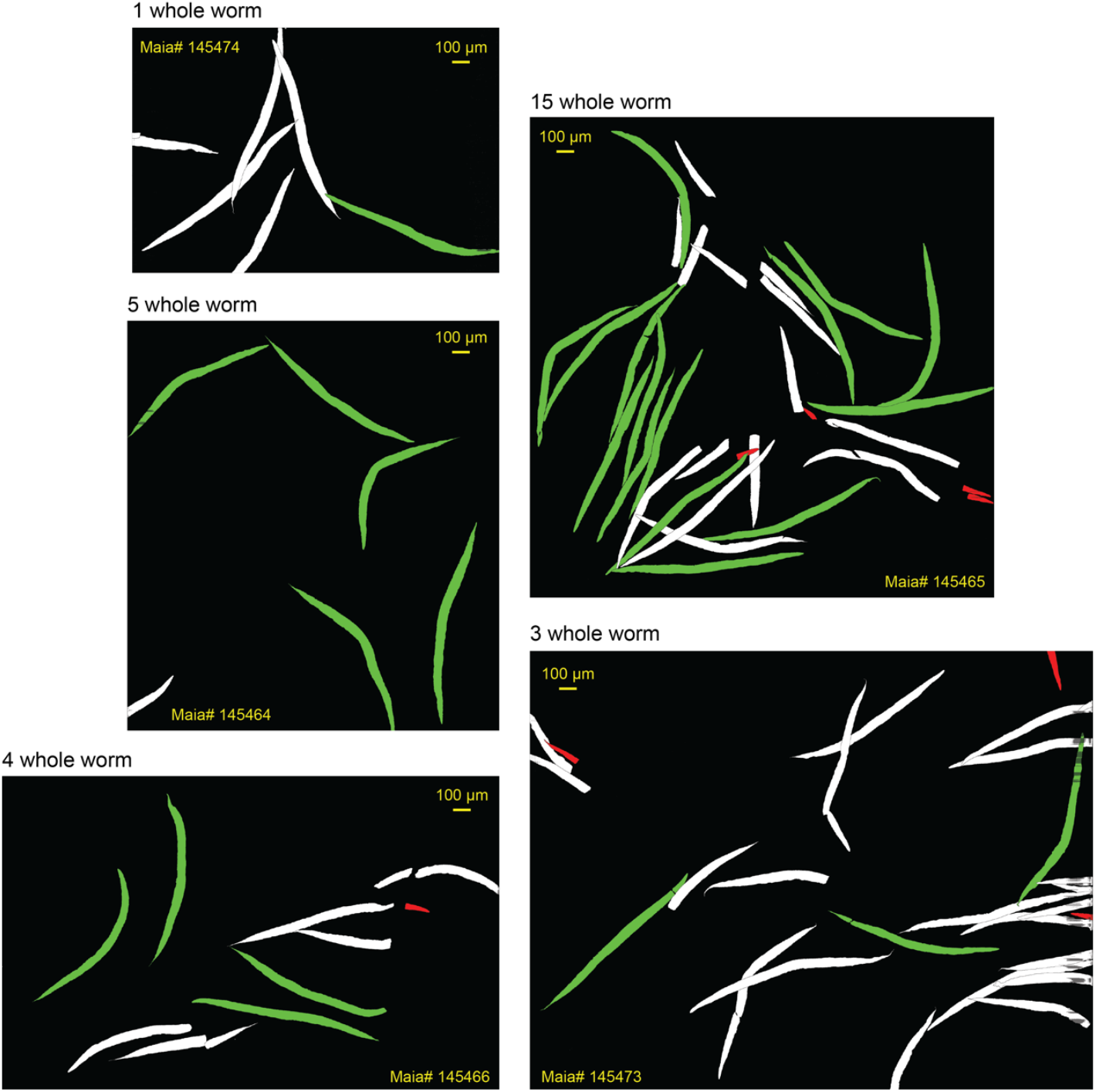
Masks for 4 day old SIH treated adults.

**Figure S1D.**
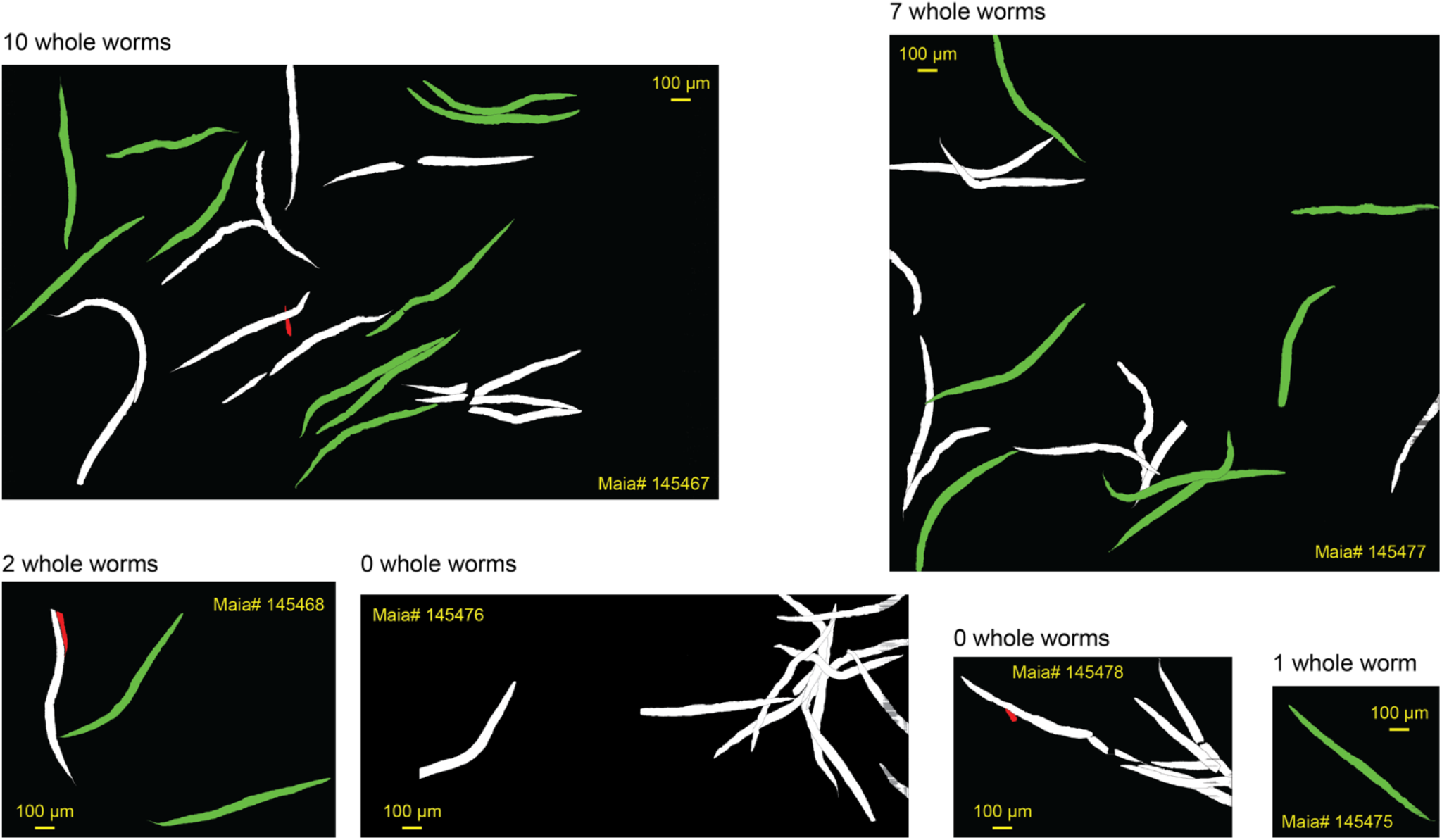
Masks for 4 day old Lip-1 treated adults.

**Figure S1E.**
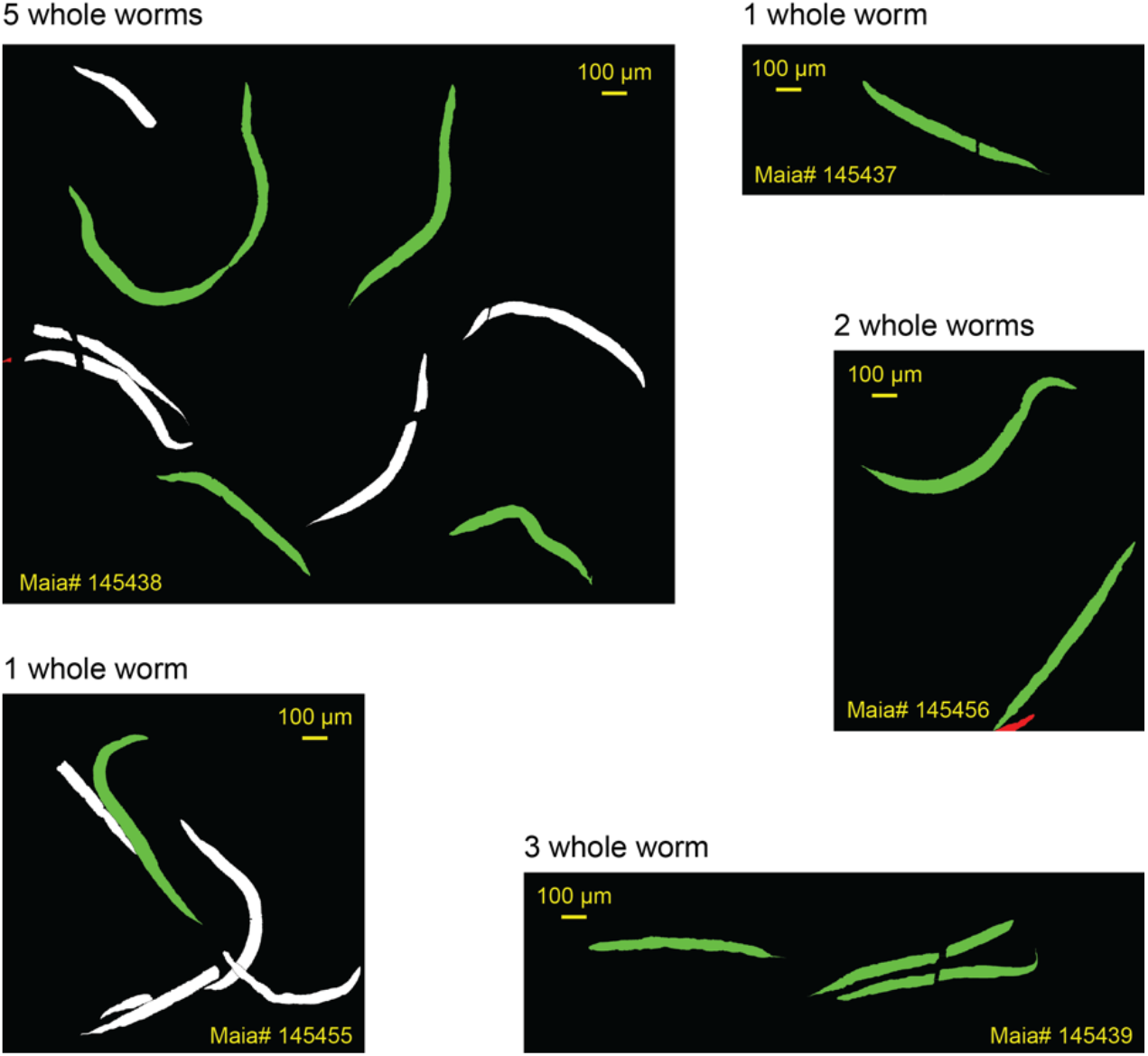
Masks for 8 day old Control adults.

**Figure S1F.**
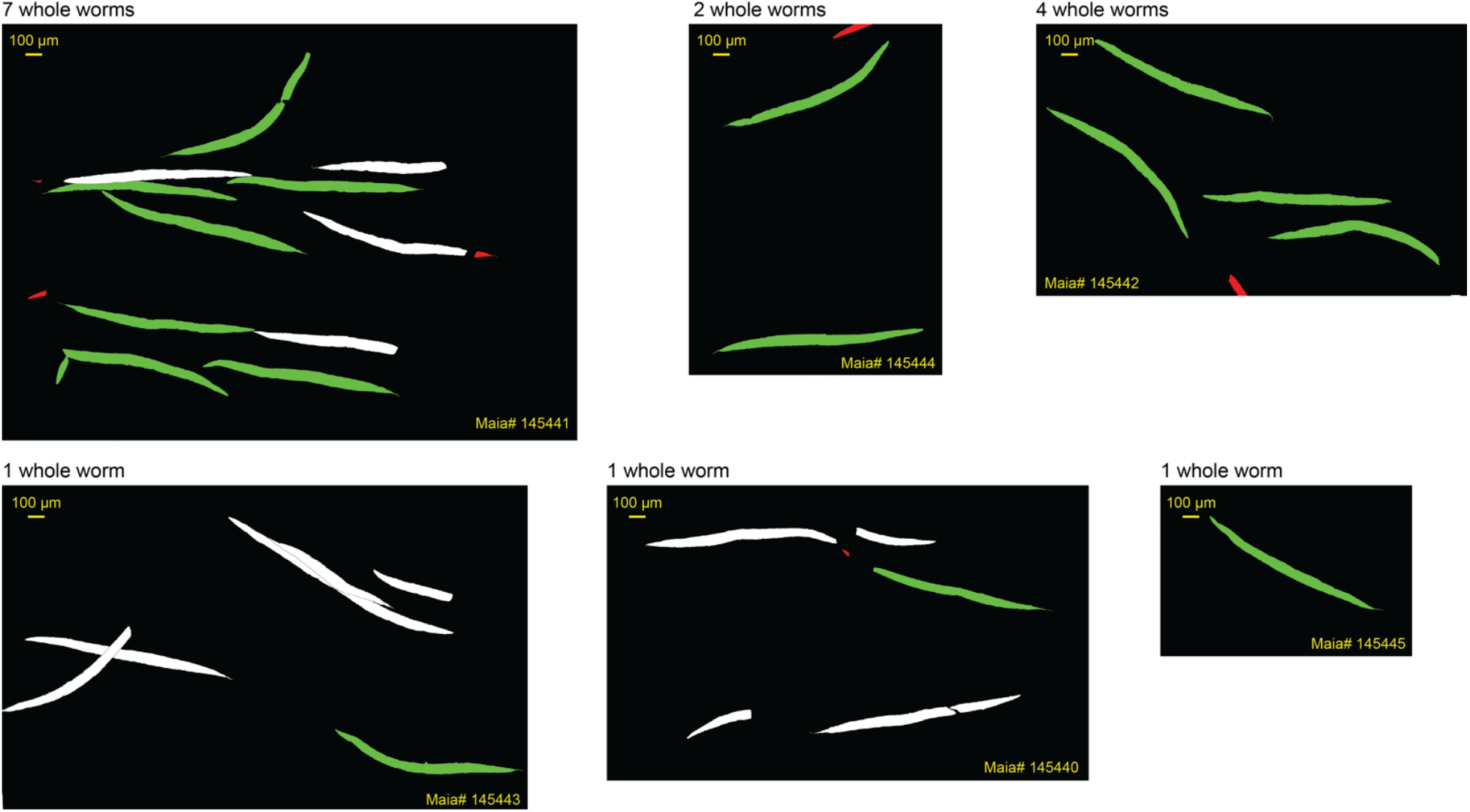
Masks for 8 day old 250 µM SIH treated adults.

**Figure S1G.**
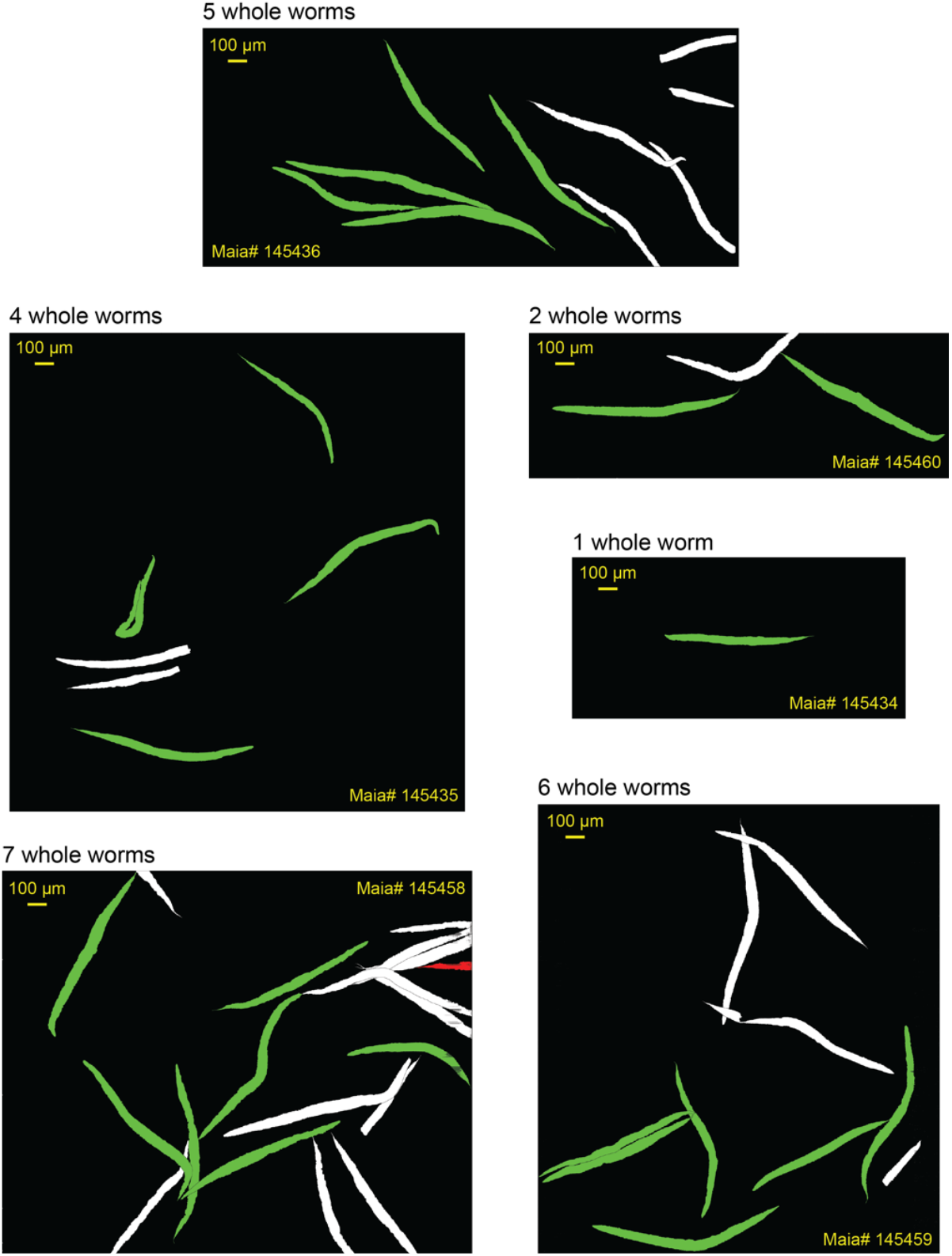
Masks for 8 day old 200 µM Lip-1 treated adults.

### Analysis of Iron from X-ray Fluorescence Microscopy

#### Total Mean areal density of iron analysis

Summary statistics and tests for normality of areal density for iron (pg µm^−2^) are included in **Table S3**. All the total iron data sets were normally distributed, as indicated below.

**Table S3:**
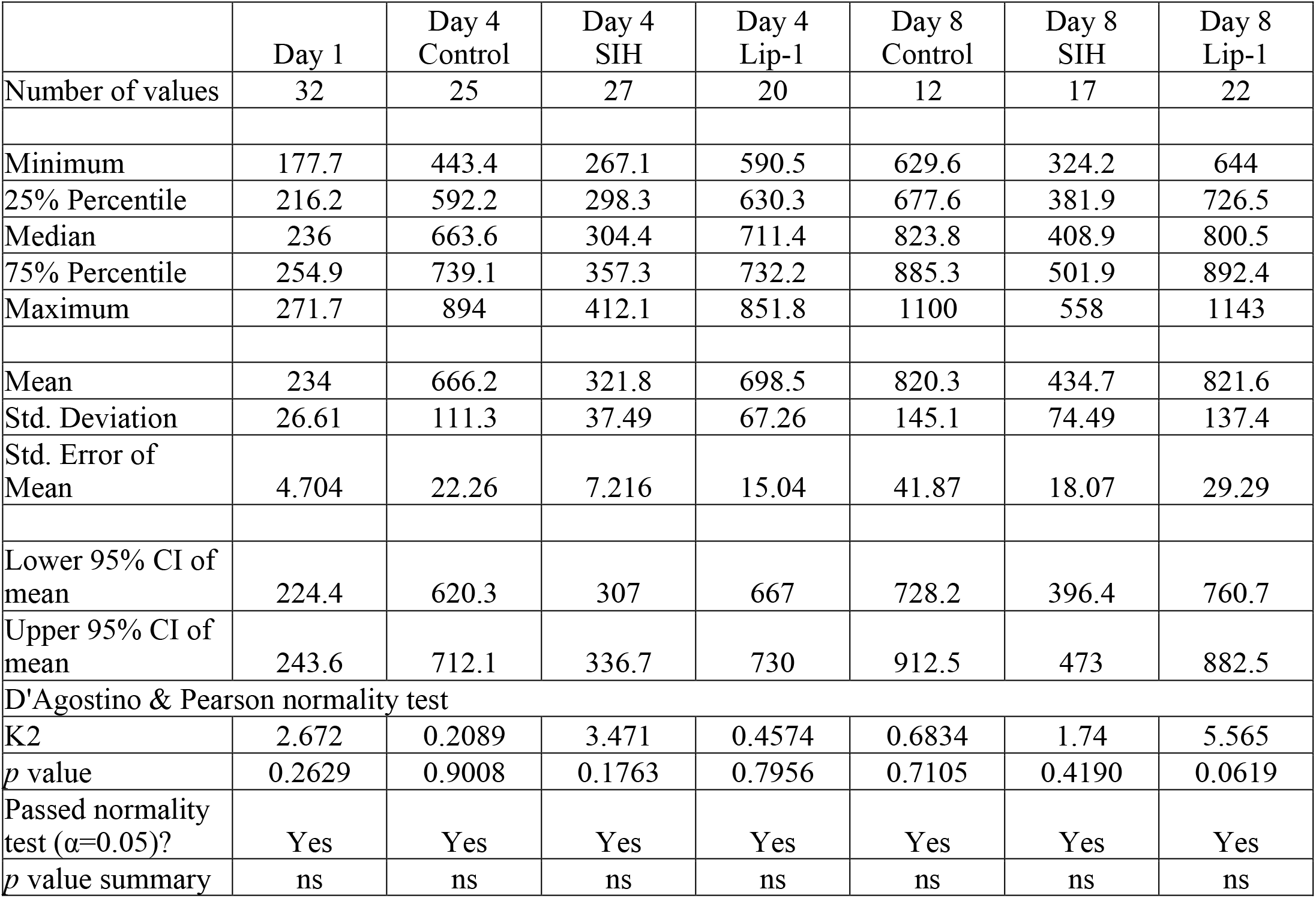
Summary of areal density iron results between treatments and ages

|  | Day 1 | Day 4 Control | Day 4 SIH | Day 4 Lip-1 | Day 8 Control | Day 8 SIH | Day 8 Lip-1 |
| --- | --- | --- | --- | --- | --- | --- | --- |
| Number of values | 32 | 25 | 27 | 20 | 12 | 17 | 22 |
| Minimum | 177.7 | 443.4 | 267.1 | 590.5 | 629.6 | 324.2 | 644 |
| 25% Percentile | 216.2 | 592.2 | 298.3 | 630.3 | 677.6 | 381.9 | 726.5 |
| Median | 236 | 663.6 | 304.4 | 711.4 | 823.8 | 408.9 | 800.5 |
| 75% Percentile | 254.9 | 739.1 | 357.3 | 732.2 | 885.3 | 501.9 | 892.4 |
| Maximum | 271.7 | 894 | 412.1 | 851.8 | 1100 | 558 | 1143 |
| Mean | 234 | 666.2 | 321.8 | 698.5 | 820.3 | 434.7 | 821.6 |
| Std. Deviation | 26.61 | 111.3 | 37.49 | 67.26 | 145.1 | 74.49 | 137.4 |
| Std. Error of Mean | 4.704 | 22.26 | 7.216 | 15.04 | 41.87 | 18.07 | 29.29 |
| Lower 95% CI of mean | 224.4 | 620.3 | 307 | 667 | 728.2 | 396.4 | 760.7 |
| Upper 95% CI of mean | 243.6 | 712.1 | 336.7 | 730 | 912.5 | 473 | 882.5 |
| D'Agostino & Pearson normality test |  |  |  |  |  |  |  |
| K2 | 2.672 | 0.2089 | 3.471 | 0.4574 | 0.6834 | 1.74 | 5.565 |
| p value | 0.2629 | 0.9008 | 0.1763 | 0.7956 | 0.7105 | 0.4190 | 0.0619 |
| Passed normality test ($\alpha=0.05$)? | Yes | Yes | Yes | Yes | Yes | Yes | Yes |
| p value summary | ns | ns | ns | ns | ns | ns | ns |

There was a significant difference between mean areal density of iron (F (6, 148) = 171.3, *p* < 0.0001) amongst the groups measured. Comparisons between age and treatment groups an Ordinary one-way ANOVA was performed, followed by Sidak’s multiple comparisons test. The results of the pairwise comparisons, corrected for multiple comparisons, are shown in **Table S4**.

**Table S4:** Summary of areal density of iron comparisons between ages and treatments

| Sidak's multiple comparisons test | Mean Diff. | 95.00% CI of diff. | Significant? | Adjusted p value |
| --- | --- | --- | --- | --- |
| Day 1 Control vs. Day 4 Control | -432.2 | -499.3 to -365.1 | Yes | <0.0001 |
| Day 1 Control vs. Day 8 Control | -586.3 | -671.5 to -501.2 | Yes | <0.0001 |
| Day 4 Control vs. Day 8 Control | -154.1 | -242.4 to -65.85 | Yes | <0.0001 |
| Day 4 Control vs. Day 4 SIH | 344.4 | 274.6 to 414.1 | Yes | <0.0001 |
| Day 4 Control vs. Day 4 Lip-1 | -32.28 | -107.7 to 43.15 | No | 0.9226 |
| Day 8 Control vs. Day 8 SIH | 385.6 | 290.9 to 480.4 | Yes | <0.0001 |
| Day 8 Control vs. Day 8 Lip-1 | -1.229 | -91.45 to 89 | No | >0.9999 |
| Day 4 SIH vs. Day 8 SIH | -112.9 | -190.7 to -35.02 | Yes | 0.0006 |
| Day 4 Lip-1 vs. Day 8 Lip-1 | -123.1 | -200.8 to -45.42 | Yes | 0.0001 |
| Day 4 SIH vs. Day 4 Lip-1 | -376.6 | -450.8 to -302.5 | Yes | <0.0001 |
| Day 8 SIH vs. Day 8 Lip-1 | -389.9 | -469 to -304.8 | Yes | <0.0001 |

#### Total body iron analysis

Summary statistics and tests for normality of total body iron (pg) are included in **Table S5**. All the total iron data sets were normally distributed, as indicated below.

**Table S5:** Summary of total body iron results between treatments and ages

|  | Day 1 | Day 4<br>Control | Day 4<br>SIH | Day 4<br>Lip-1 | Day 8<br>Control | Day 8<br>SIH | Day 8<br>Lip-1 |
| --- | --- | --- | --- | --- | --- | --- | --- |
| Number of values | 32 | 25 | 27 | 20 | 12 | 17 | 22 |
| Minimum | 13.48 | 91.73 | 41.07 | 71.51 | 122.8 | 74.16 | 100 |
| 25% Percentile | 22.06 | 109.2 | 61.28 | 91.34 | 134.3 | 117.5 | 128.9 |
| Median | 25.36 | 124.4 | 66.78 | 114.6 | 146.6 | 127.5 | 157.2 |
| 75% Percentile | 27.75 | 138.6 | 71.8 | 129.5 | 173.2 | 169.5 | 189.6 |
| Maximum | 39.6 | 171.1 | 98.92 | 148.6 | 214.2 | 202 | 277.9 |
| Mean | 24.79 | 126.1 | 68.37 | 113.3 | 154.4 | 138.6 | 165.1 |
| Std. Deviation | 5.166 | 21.05 | 12.1 | 21.91 | 27.94 | 36.58 | 44.93 |
| Std. Error of Mean | 0.9133 | 4.211 | 2.328 | 4.9 | 8.065 | 8.873 | 9.579 |
| Lower 95% CI of mean | 22.93 | 117.4 | 63.59 | 103.1 | 136.7 | 119.8 | 145.2 |
| Upper 95% CI of mean | 26.65 | 134.8 | 73.16 | 123.6 | 172.2 | 157.4 | 185 |
| D'Agostino & Pearson normality test |  |  |  |  |  |  |  |
| K2 | 2.708 | 0.8963 | 4.252 | 1.75 | 3.123 | 0.6397 | 4.128 |
| *p* value | 0.2582 | 0.6388 | 0.1193 | 0.4169 | 0.2099 | 0.7262 | 0.1270 |
| Passed normality test<br>($\alpha=0.05$)? | Yes | Yes | Yes | Yes | Yes | Yes | Yes |
| *p* value summary | ns | ns | ns | ns | ns | ns | ns |

There was a significant difference between total body iron (F(6,148)=97.3, *p* < 0.0001) amongst the groups measured. Comparisons between age and treatment groups an Ordinary one-way ANOVA was performed, followed by Sidak’s multiple comparisons test. The results of the pairwise comparisons, corrected for multiple comparisons, are shown in **Table S6**.

**Table S6:** Summary of total body iron between ages and treatments.

| Sidak's multiple comparisons test | Mean Diff. | 95.00% CI of diff. | Significant? | Adjusted $p$ Value |
| --- | --- | --- | --- | --- |
| Day 1 Control vs. Day 4 Control | -101.3 | -120.9 to -81.67 | Yes | <0.0001 |
| Day 1 Control vs. Day 8 Control | -129.6 | -154.5 to -104.8 | Yes | <0.0001 |
| Day 4 Control vs. Day 8 Control | -28.35 | -54.16 to -2.538 | Yes | 0.0211 |
| Day 4 Control vs. Day 4 SIH | 57.71 | 37.31 to 78.11 | Yes | <0.0001 |
| Day 4 Control vs. Day 4 Lip-1 | 12.74 | -9.306 to 34.79 | No | 0.6818 |
| Day 8 Control vs. Day 8 SIH | 15.8 | -11.91 to 43.51 | No | 0.6991 |
| Day 8 Control vs. Day 8 Lip-1 | -10.7 | -37.07 to 15.68 | No | 0.9550 |
| Day 4 SIH vs. Day 8 SIH | -70.26 | -93.01 to -47.5 | Yes | <0.0001 |
| Day 4 Lip-1 vs. Day 8 Lip-1 | -51.79 | -74.49 to -29.08 | Yes | <0.0001 |
| Day 4 SIH vs. Day 4 Lip-1 | -44.97 | -66.65 to -23.29 | Yes | <0.0001 |
| Day 8 SIH vs. Day 8 Lip-1 | -26.49 | -50.23 to -2.763 | Yes | 0.0178 |

### Analysis of Iron from X-ray Absorption Near Edge Spectroscopy (*φ*-XANES)

**Figure S2:**
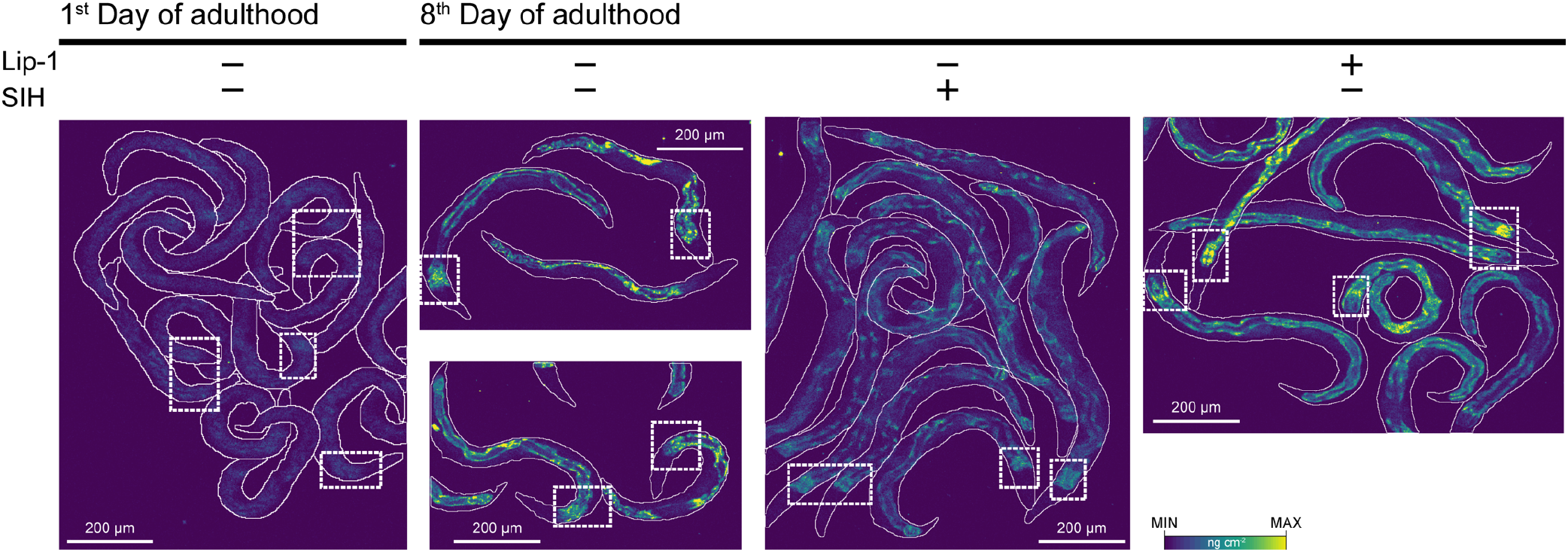
XFM maps of Fe and regions-of-interests for *φ*-XANES analysis (dashed boxes).

Summary of pooled spectra for young (*n*=6), aged (*n*=4) TJ1060 animals and aged animals with animals with SIH (*n*=4) or Lip-1 treatment (*n*=5) (*i.e.* the mean for all pixels in all ROIs within each group) from scanning the iron K-edge, including features present in the pre-edge (∼7.112 keV) and rising edge (∼7.124 keV) are shown in **Fig S3A**, with their corresponding first-derivatives (**Fig S3B**). Similarly, pooled spectra and first derivatives from 4 day old adult wild types treated with (*n*=4) and without (*n*=4) DEM is shown in **Fig S3C-D**.

**Figure S3:**
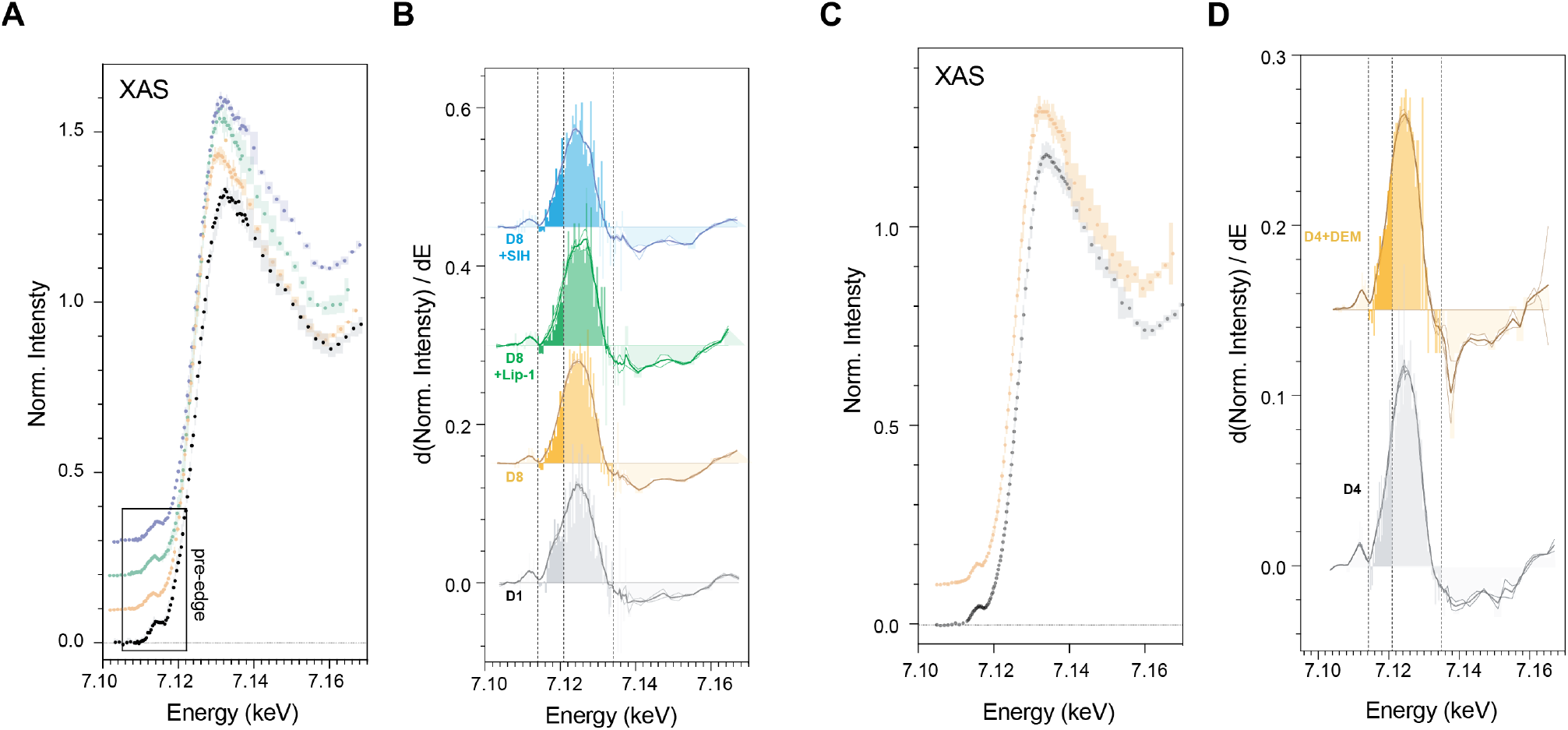
Changes in the first derivative of the Fe K-edge XANES reflect variation in the intensity of the 1s → 4s and 1s → 4p transitions. **A**) Iron K-edge *φ*XANES spectra (circles) determined by integrating 105 XFM micrographs acquired at energies spanning the iron absorption edge (7000 - 7455 eV). Shaded region represents 95 % CI; spectra offset for clarity. Shown are spectra for Day 1 adults, Day 8 adults, Day 8+Lip and Day 8+SIH groups. **B**) The corresponding first derivative for spectra shown in **A**; spectra offset in the vertical for clarity. Darker shading (7.114 – 7.121 keV) marks the area under the curve attributed the 1s → 4s transition. The lighter shading (7.121 – 7.132 keV) marks the are under the curve attributed to the 1s → 4p transition. **C**) Pooled spectra for Day 4 adults treated with and without DEM, and D) their matching first derivatives, as described above.

#### Estimation of Fe^2+^

There was a significant difference between the fractional Fe^2+^/Fe(total) estimates, determined by non-overlapping 95% CI, between aged TJ1060 animals (**Table S7**). Treatment with Lip-1 or SIH restored the Fe^2+^/Fe(total) estimate. Similarly, treatment of wild type (N2) animals with DEM markedly increased Fe^2+^/Fe(total).

**Table S7:** Summary of the estimated Fe^2+^/Fe(total) for each treatment group.

| | Group | Mean $\text{Fe}^{2+}/\text{Fe}(\text{total})$ | 95% CI |
| --- | --- | --- | --- |
| **TJ1060** | Day 1 | 0.239 | 0.233-0.244 |
|  | Day 8 Control | 0.300 | 0.295-0.305 |
|  | Day 8 Lip-1 | 0.231 | 0.198-0.262 |
|  | Day 8 SIH | 0.236 | 0.232-0.240 |
| **N2** | Day 4 | 0.228 | 0.223-0.233 |
|  | Day 4 + DEM | 0.310 | 0.305-0.314 |

### Lifespan analysis

**Table S8:** Summary of survival data from 8 **independent** replicate experiments. Median and maximum lifespan figures are days of adulthood at 25 (±1) °C. Censored individuals are those that were lost, primarily due to crawling off the side of the plate. Median lifespan was initially compared using a Log-rank (Mantel-Cox) test. * p<0.0001; † p=0.0013

| Replicate | control |  |  |  | 250 µM SIH |  |  |  |  | 200 µM Lip-1 |  |  |  |  |
| --- | --- | --- | --- | --- | --- | --- | --- | --- | --- | --- | --- | --- | --- | --- |
|  | death events | censored | median | max. | death events | censored | median | max. | % median ↑ | death events | censored | median | max. | % median ↑ |
| 1 | 88 | 3 | 7 | 17 | 71 | 4 | 14* | 19 | 100 | 103 | 1 | 13* | 25 | 86 |
| 2 | 61 | 10 | 8 | 16 | 96 | 7 | 14* | 19 | 75 | 88 | 6 | 11† | 20 | 38 |
| 3 | 109 | 3 | 9 | 21 | 81 | 7 | 16* | 23 | 78 | 108 | 2 | 14* | 24 | 56 |
| 4 | 99 | 1 | 7 | 19 | 111 | 10 | 16* | 20 | 129 | 147 | 1 | 14* | 24 | 100 |
| 5 | 91 | 16 | 9 | 19 | 93 | 9 | 14* | 18 | 56 | 86 | 7 | 14* | 26 | 56 |
| 6 | 97 | 11 | 8 | 16 | 105 | 10 | 14* | 24 | 75 | 112 | 6 | 12* | 25 | 50 |
| 7 | 93 | 2 | 7 | 21 | 90 | 29 | 17* | 29 | 143 | 80 | 6 | 15* | 27 | 114 |
| 8 | 72 | 5 | 8 | 22 | 73 | 13 | 20* | 26 | 150 | 85 | 5 | 14* | 26 | 75 |
| Total | 709 | 51 |  |  | 720 | 89 |  |  |  | 809 | 34 |  |  |  |
| Mean |  |  | 8 | 19 |  |  | 16 | 22 | +101% |  |  | 13 | 25 | +72% |

**Figure S4:**
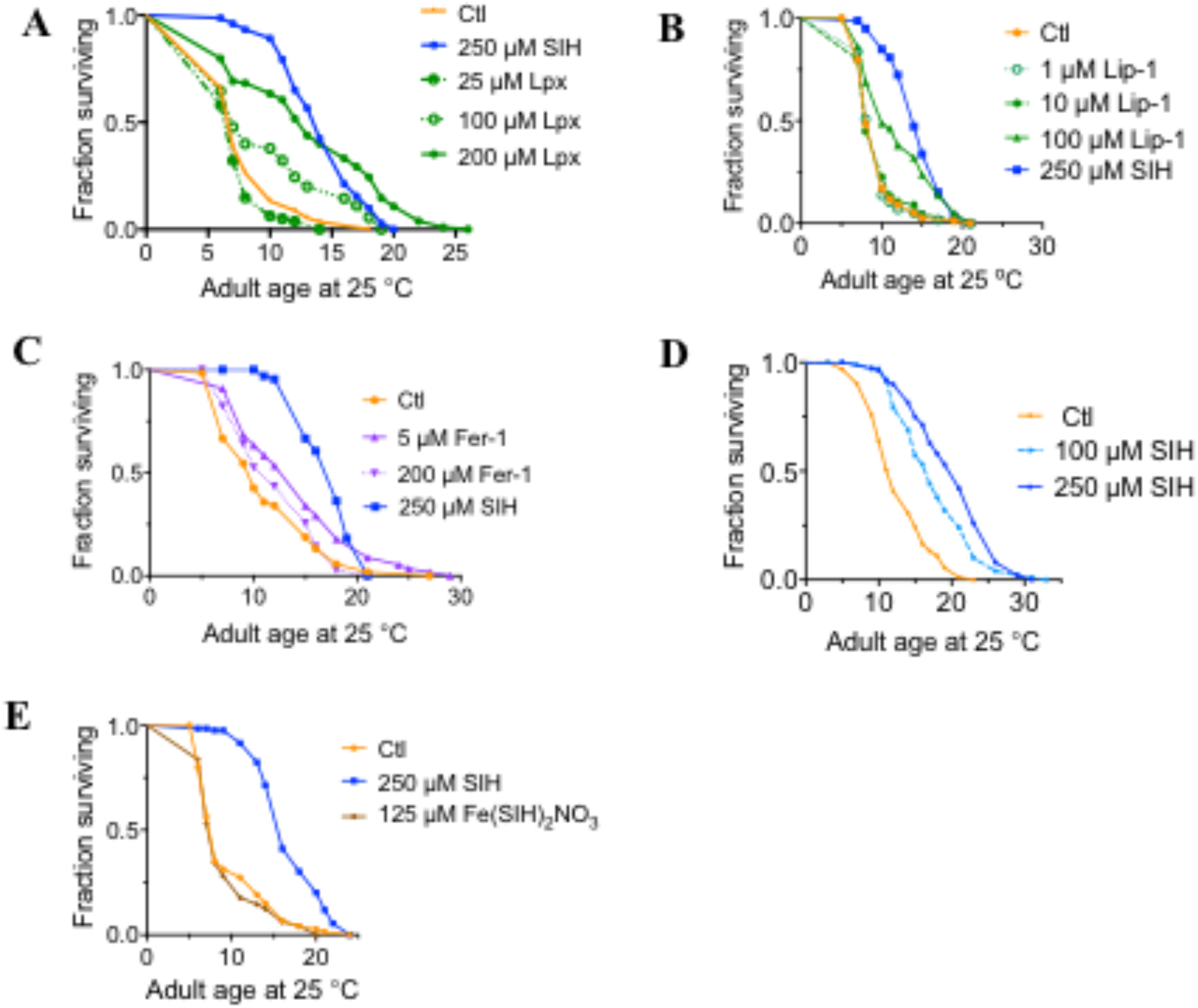
Dose response of SIH, Liproxstatin-1 and Ferrostatin-1. **A:** Median survival (Log-rank (Mantel-Cox) test: Control=6 days; 25 µM Lip-1=6 days (*p*=0.07 ns); 100 µM Lip-1=6 days (*p*=0.006); 200 µM Lip-1=12 days (*p*<0.0001); 250 µM SIH=13 days (*p*<0.0001) **B:** Median survival (Log-rank (Mantel-Cox) test: Control=7 days; 1 µM Lip-1=9 days (*p*=0.90 ns); 10 µM Lip-1=7 days (*p*=0.78 ns); 100 µM Lip-1=9 days (*p*<0.0001); 250 µM SIH=13 days (*p*<0.0001) **C:** Median survival (Log-rank (Mantel-Cox) test: Control=10 days; 5 µM Fer-1=15 days (*p*=0.0047); 200 µM Fer-1=12 days (*p*=0.35 ns); 250 µM SIH=18 days (*p*<0.0001) **D:** Median survival (Log-rank (Mantel-Cox) test: Control=12 days; 100 µM SIH=17 days (*p*<0.0001); 250 µM SIH=21 days (*p*<0.0001) **E:** Median survival (Log-rank (Mantel-Cox) test: Control=8 days; 125μM Fe(SIH)2NO3, equimolar SIH=8 days (*p*=0.51 ns); 250 µM SIH=16 days (*p*<0.0001)

### Testing Departure from Temporal Rescaling

Following the recently published results of Stroustrup *et al*. ^37^ we determined whether the results observed with both the SIH and Liproxstatin interventions were due to temporal scaling of aging.

Using the modified Kolmogorov-Smirnov (K-S) test ^71^ we examined whether the treatment effects can be reasonably modelled using the Accelerated Failure Time (AFT) model to determine whether we can reasonably assume that the treatment effect manifests in temporal rescaling. To control for inter-replicate differences, the **test is conducted on the residuals a replicate-specific AFT model with the Buckley-James method** ^72^ **using the nonparametric baseline hazards form**. The function *bj* in R package *rms* was used to fit the models. The null hypothesis for the two-sample K-S test is that the simple temporal rescaling holds and the residuals for the two treatment groups under comparison come from the same distribution.

Using the modified Kolmogorov-Smirnov (K-S) test, we examined whether the treatment effects can be reasonably modelled using the AFT model to determine whether we can reasonably assume that the treatment effect manifests in temporal rescaling. To control for inter-replicate differences, the **test is conducted on the residuals from replicate-specific AFT model with the Buckley-James method that uses nonparametric baseline hazards form**. The function *bj* in R package *rms* was used to fit the models. The null hypothesis for the two-sample K-S test is that the simple temporal rescaling holds and the residuals for the two treatment groups under comparison come from the same distribution.

Since the R function can only take right-censored data and our lifespan data are interval-censored, we use the mid-point of the interval to assign the time of event. Treating interval-censored data as right-censored is expected to underestimate the variability in the statistical estimates ^73^ which in turn will produce an optimistic (smaller than it should be) *p*-value. To reduce the likelihood of false rejection of the null hypothesis merely because of the optimistic *p*-value, we chose a more stringent Type I error (0.01) than the usual 0.05 when conducting the K-S test.

**Table S9:** *p*-values of KS test on Residuals of nonparametric AFT models

| Replicate | Control vs Lip-1 | Control vs SIH | Lip-1 vs SIH |
| --- | --- | --- | --- |
| 1 | $9 \times 10^{-5}$ | $2 \times 10^{-3}$ | $1 \times 10^{-15}$ |
| 2 | $2 \times 10^{-2}$ | $4 \times 10^{-3}$ | $8 \times 10^{-5}$ |
| 3 | $7 \times 10^{-3}$ | $9 \times 10^{-4}$ | $4 \times 10^{-11}$ |
| 4 | $4 \times 10^{-13}$ | $3 \times 10^{-15}$ | $2 \times 10^{-31}$ |
| 5 | $2 \times 10^{-4}$ | $1 \times 10^{-4}$ | $7 \times 10^{-28}$ |
| 6 | $1 \times 10^{-9}$ | $3 \times 10^{-5}$ | $2 \times 10^{-5}$ |
| 7 | $3 \times 10^{-6}$ | $4 \times 10^{-14}$ | $1 \times 10^{-7}$ |
| 8 | $1 \times 10^{-6}$ | $4 \times 10^{-6}$ | $1 \times 10^{-8}$ |

As can be seen from **Table S9**, the effect of Lip-1 and SIH treatment relative to control always deviates away from simple temporal rescaling (all *p*-values < 10^−2^) with the exception of Lip-1 in replicate 2. **Figure S5** shows graphically why the AFT assumption is not reasonable since the survival curves of residuals from the AFT models show ‘crossing’ behavior. If the simple temporal rescaling assumption is reasonable, we would expect the survival curves for the different treatments to be very similar to each other. The observed crossing of the curves is primarily caused by the de-acceleration in the survival function for control worms.

When all the replicates are combined, and the modified KS test were performed on the residuals of the AFT models with the best parametric form, we found that the *p*-value for comparing Control *vs* Lip-1, Control *vs* SIH and Lip-1 *vs* SIH are 2 x 10^−24^, 1 x 10^−24^ and 3 x 10^−37^ respectively. These results indicate that failure to control for inter-replicate differences would lead to even stronger evidence of departure from simple temporal rescaling.

**Figure S5:**
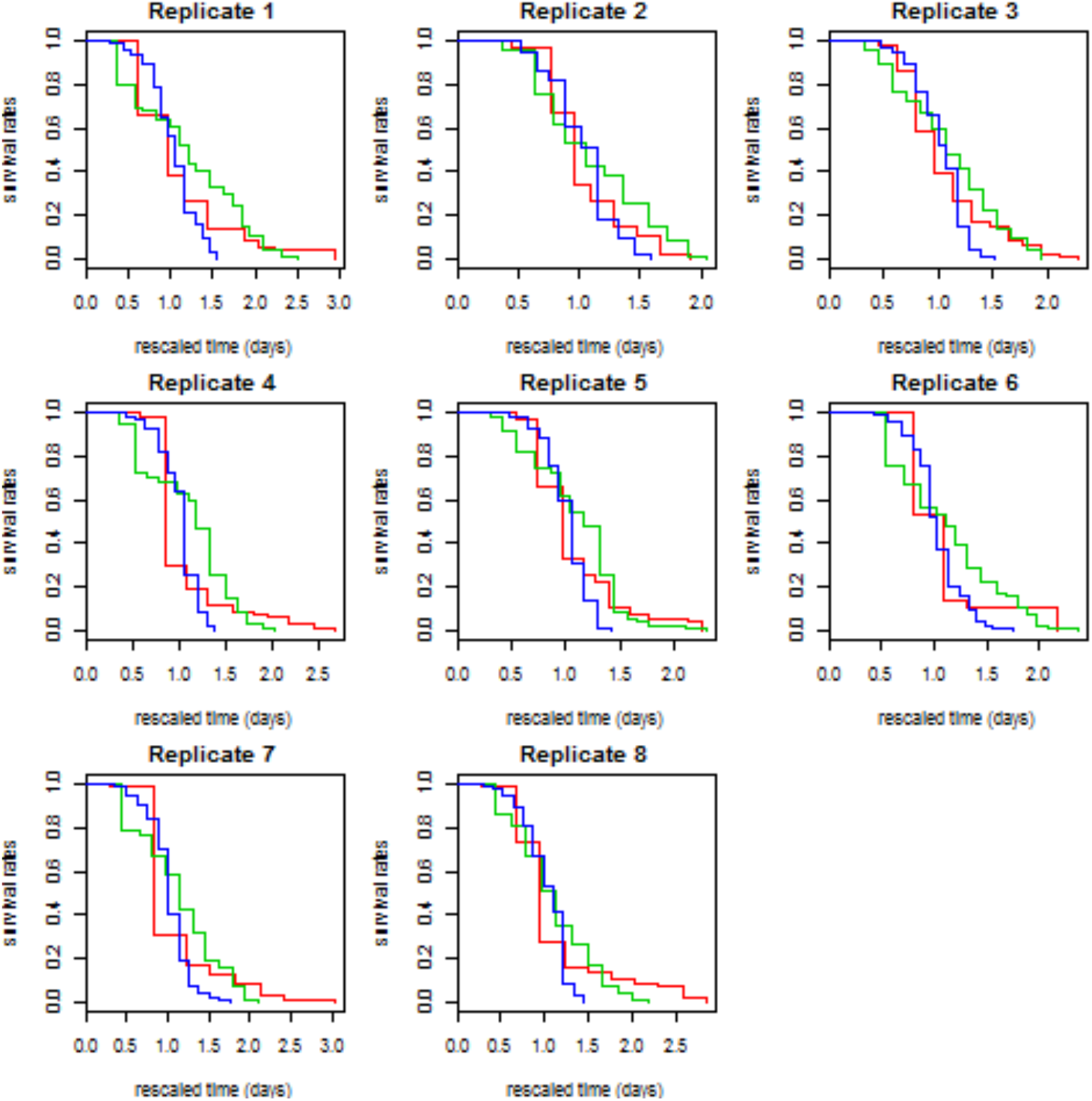
The residuals of AFT model with nonparametric hazards form for different replicates (red = control, green = Lip-1, blue = SIH). The x-axis has been rescaled to remove the temporal rescaling effect. Under the null hypothesis, we expect the residuals for the different groups to have the same survival curves.

### Determining AFT models with the best baseline hazard form

In order to investigate the possible reasons for departure from simple temporal rescaling, we used parametric survival models which require specification of a parametric baseline hazard form. To minimize the risk of model misspecification, we identified the most appropriate baseline hazard form for each replicate using the Bayesian Information Criterion (BIC), with the best parametric form chosen as the model that minimizes the BIC.

The following parametric baseline hazards were fitted:

Gompertz, Gompertz with Frailty, Weibull, Weibull with Frailty, Log-normal and Log-logistic. The mathematical formulae for each parametric form are detailed below:

Gompertz: h(t|a,b) = (a/b)exp(t/b)

Gompertz with frailty: h(t|a,b,σ) = (a/b)exp(t/b)/[1 + σ^2^a exp((t/b) − 1)]

Weibull: h(t|α,β) = (α/β)(t/β)^α−1^

Weibull with frailty: h(t|α,β,σ) = (α/)(t/β)^α−1^/[1 + σ^2^(t/β)^α^]

Log-normal: h(t|µ,σ) = ϕ((log t − µ)/σ)/σt[1 − Φ ((log t − µ)/σ)]

Log-logistic: h(t|α,β) = (α/β)(t/β)^α−1^[1 + (t/β)^α^]

Here *ϕ* and Φ denote the probability density function (PDF) and cumulative distribution function (CDF), respectively, of the standard normal distribution; *µ* and *σ* denote the mean and standard deviation (in the case of the log-normal, the mean and standard deviation of the logarithm of *x*); *λ*, *α*, and *a* are shape parameters; *β* and *b* are scale parameters. In the case of frailty, individual hazards *h_i_*(*t*) are related to a baseline hazard by a random factor *Z* that follows a Gamma distribution with mean 1 and variance *σ*^2^.

Bayesian Information Criterion (BIC) is used to determine the best parametric form of the hazards; with better fit indicated by lower BIC value. All computations are done using flexsurv R package, taking into account that events are interval censored to account for the fact that we do not observe the exact event time and only know that events occurred within an interval (a,b).

**Figure S6:**
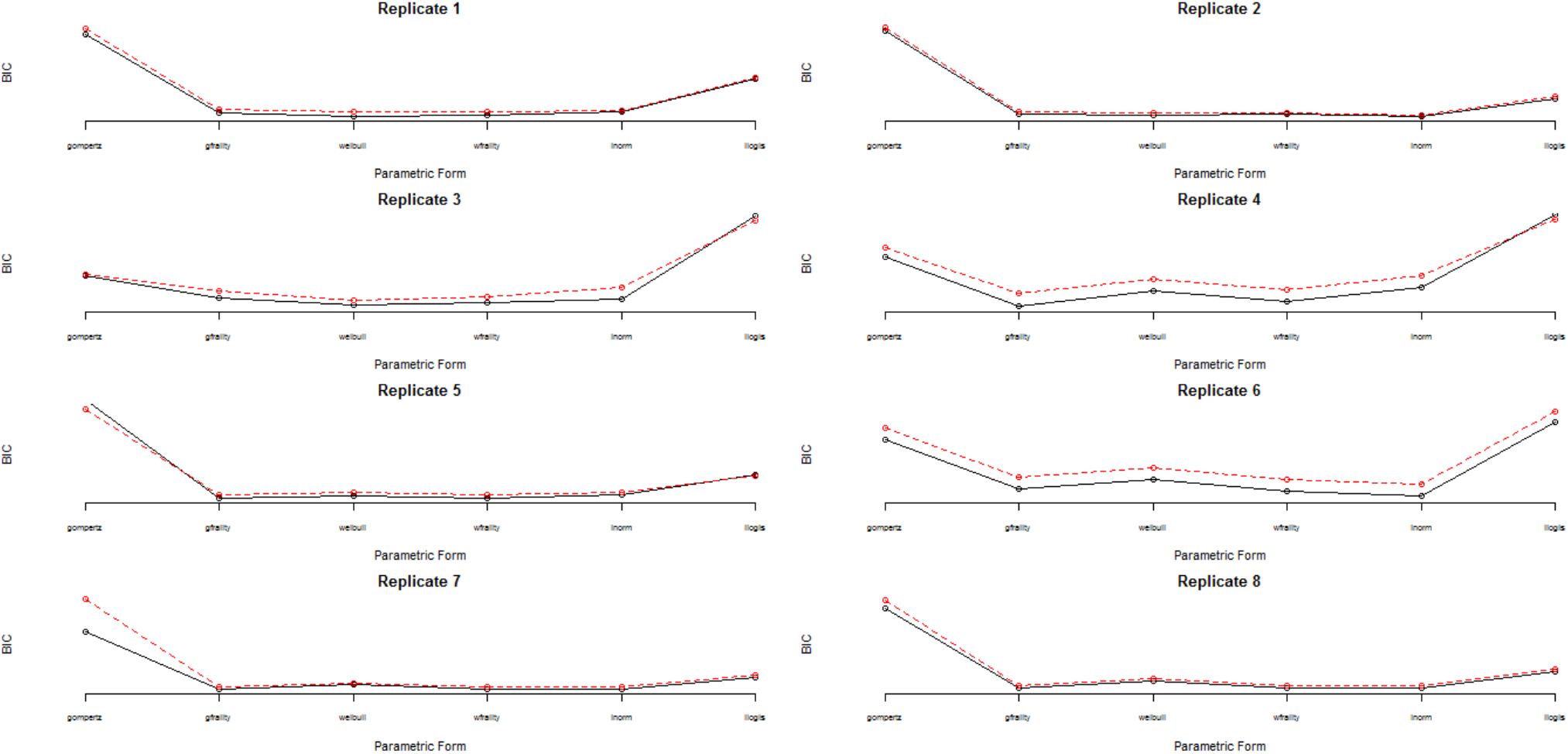
Bayesian Information Criterion (BIC) for AFT models with different parametric baseline hazard forms

From **Figure S6**, we can see that Gompertz baseline hazard form does not fit the data well, except when frailty is used. Weibull baseline hazard fits some replicates quite well and the fit is further improved when frailty is assumed. In fact, Weibull with frailty provides the best parametric baseline hazards form for nearly all the replicates, followed closely by the log-normal models.

### Possible Causes of Departure from Temporal Rescaling

Stroustrup *et al.* ^37^ show that unobserved heterogeneity (*e.g.* due to heterogeneity in the temperature the worms were exposed to) could cause de-acceleration and further, when the degree of heterogeneity is different between treatments, this could give rise to apparent departure from temporal rescaling. We investigated whether there is significant difference in the degree of heterogeneity by comparing two models for each replicate: **(M1)** model with Weibull frailty (Weibull hazard, Gamma frailty) where the degree of heterogeneity (represented by parameter s^2^ and *a*) is assumed to be the same for all three treatments, **(M2)** where the parameter s^2^ is allowed to be different but parameter *a* fixed across treatments and **(M3)** where the parameter s^2^ and a *are* allowed to be different across treatments. We compared the three models based on their BIC values and also performed likelihood ratio tests, comparing M1 *vs* M2 and M1 *vs* M3.

**Note that only M1 can be classified as an AFT model while M2 and M3 are not AFT models**, as the treatment effects also manifest in the other parameters apart from the location (shift) parameter.

**Table S10** shows that both M3 and M2 provide better fit than M1 for all replicates as indicated by small likelihood ratio test (LRT) p-values, with M3 providing more convincing p-values.

**Table S10:**
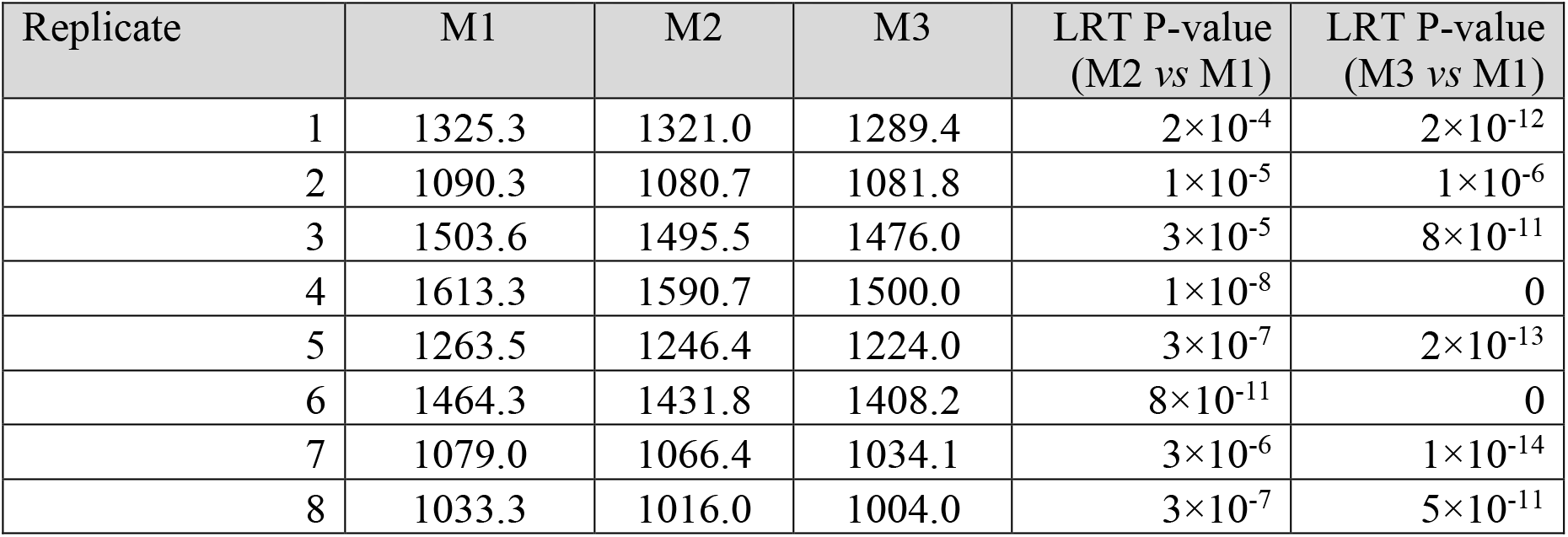
BIC values for AFT model with Weibull frailty baseline hazards (M1), non-AFT model with Weibull frailty baseline hazards and treatment-dependent heterogeneity levels σ^2^(M2) and non-AFT model with Weibull frailty baseline hazards and treatment-dependent shape parameter (a) and heterogeneity levels σ^2^(M3)

| Replicate | M1 | M2 | M3 | LRT P-value<br>(M2 vs M1) | LRT P-value<br>(M3 vs M1) |
| --- | --- | --- | --- | --- | --- |
| 1 | 1325.3 | 1321.0 | 1289.4 | $2 \times 10^{-4}$ | $2 \times 10^{-12}$ |
| 2 | 1090.3 | 1080.7 | 1081.8 | $1 \times 10^{-5}$ | $1 \times 10^{-6}$ |
| 3 | 1503.6 | 1495.5 | 1476.0 | $3 \times 10^{-5}$ | $8 \times 10^{-11}$ |
| 4 | 1613.3 | 1590.7 | 1500.0 | $1 \times 10^{-8}$ | 0 |
| 5 | 1263.5 | 1246.4 | 1224.0 | $3 \times 10^{-7}$ | $2 \times 10^{-13}$ |
| 6 | 1464.3 | 1431.8 | 1408.2 | $8 \times 10^{-11}$ | 0 |
| 7 | 1079.0 | 1066.4 | 1034.1 | $3 \times 10^{-6}$ | $1 \times 10^{-14}$ |
| 8 | 1033.3 | 1016.0 | 1004.0 | $3 \times 10^{-7}$ | $5 \times 10^{-11}$ |

The fitted survival curves for M3 model in each replicate are compared to the observed curves in **Figure S7**. To investigate whether M3 provides an adequate fit to the data, for each replicate, we performed a chi-square goodness of fit test, comparing the observed survival curve to the fitted curve for each treatment group. The results are presented in **Table S11**. While the controls and SIH are always well-fitted by the Weibull frailty models (all *p*-values >0.01), the Lip-1 data from replicates 1, 4, 5 and 7 are not adequately fitted by the Weibull frailty model. The lack of fit for replicate 4 in particular is mainly caused by the estimated survival underestimating the observed counterparts in the middle-section between 7 and 15 days and overestimation on the tails.

**Table S11:** Chi-square Goodness of Fit *p*-value for M3 (non-AFT model with treatment-dependent shape and heterogeneity parameters)

| Replicate | Control | Lip-1 | SIH |
| --- | --- | --- | --- |
| 1 | 0.4 | $4 \times 10^{-4}$ | 0.8 |
| 2 | 0.08 | 0.07 | 0.1 |
| 3 | 0.05 | 0.08 | 0.1 |
| 4 | 0.05 | $1 \times 10^{-12}$ | 0.5 |
| 5 | 0.4 | $2 \times 10^{-5}$ | 0.8 |
| 6 | 0.001 | 0.01 | 0.2 |
| 7 | 0.2 | 0.001 | 0.4 |
| 8 | 0.1 | 0.07 | 0.07 |

**Figure S7:**
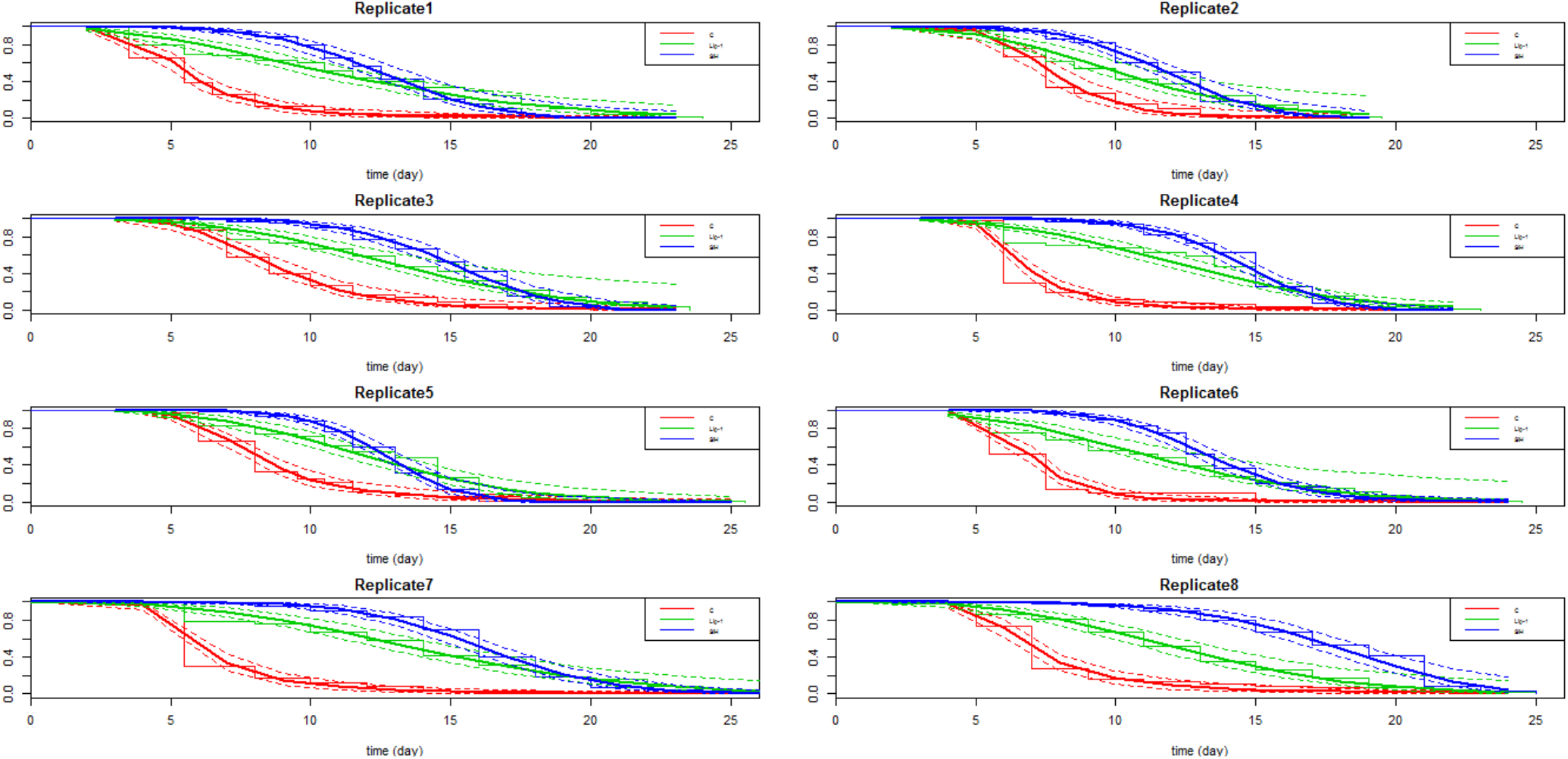
The observed and fitted survival curves based on the best non-AFT model for each replicate

### Combining Replicates

#### Models for Combining Similar Replicates

We tried to identify the most parsimonious model that can best fit the combined data from all replicates. If some of the parameters are quite similar across replicates, we can fit a simpler model than the saturated model where all parameters are allowed to be different across replicates. A range of models are fitted and the best simple model for the combined data is Model 6 (M6) with the same treatment-dependent shapes and heterogeneity parameters across replicates but replicate-specific parameters for scale parameter of the control worms and temporal rescaling parameters. The BIC value for this model is smaller than that for the saturated model (15079 *vs* 15347) and the LR test statistic is 79.2 (df = 90) with *p*-value = 0.78, indicating that based on LR test the combined model (M6) does not provide worse fit to the data. The need for replicate-specific scale parameters and temporal rescaling parameters corroborates the evidence in **Figure S7** which showed these parameters as having considerable variations across replicates.

Goodness-of-fit (GOF) test at replicate-level based on model M6 (**Table S12**) shows that this model provides more or less the same level of fit to the replicate-specific model (**Table S11**), with replicates showing good fit before still showing good fit now. The fitted survival curves (based on M6) are given in **Figure S8**.

**Table S12:** Chi-square Goodness of Fit Statistics (*p*-values) for M6 (the best parsimonious model according to BIC)

| Replicate | Control | Lip-1 | SIH |
| --- | --- | --- | --- |
| 1 | $6 \times 10^{-1}$ | $10^{-6}$ | $5 \times 10^{-1}$ |
| 2 | $2 \times 10^{-1}$ | $5 \times 10^{-2}$ | $4 \times 10^{-2}$ |
| 3 | $9 \times 10^{-3}$ | $10^{-1}$ | $2 \times 10^{-1}$ |
| 4 | $10^{-2}$ | $4 \times 10^{-11}$ | $2 \times 10^{-1}$ |
| 5 | $5 \times 10^{-1}$ | $4 \times 10^{-5}$ | $4 \times 10^{-1}$ |
| 6 | $3 \times 10^{-7}$ | $10^{-2}$ | $4 \times 10^{-2}$ |
| 7 | $3 \times 10^{-1}$ | $3 \times 10^{-3}$ | $5 \times 10^{-1}$ |
| 8 | $2 \times 10^{-1}$ | $2 \times 10^{-1}$ | $7 \times 10^{-2}$ |

**Figure S8:**
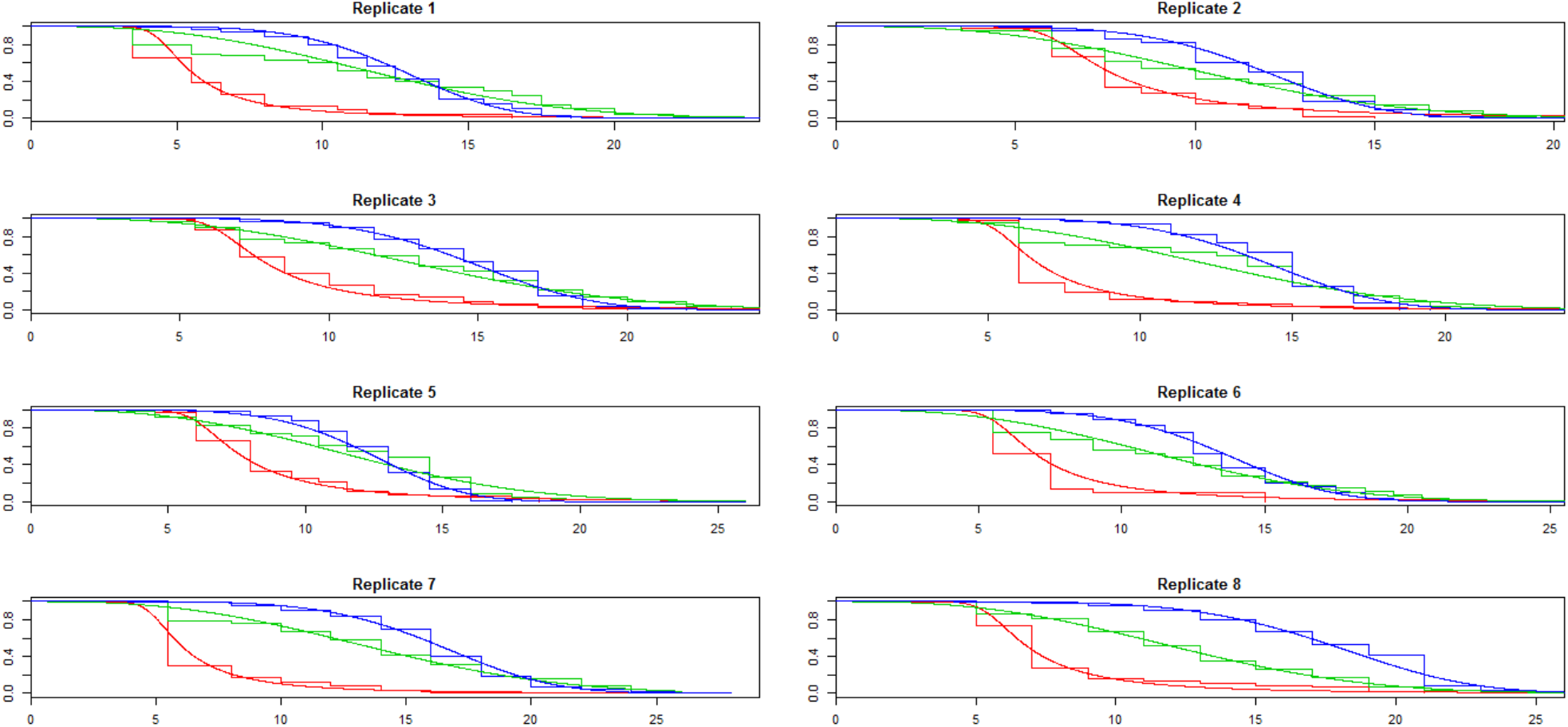
The observed and fitted survival curves based on model the most parsimonious model M6

### Meta-Analysis

In **Table S13**, we present the parameter estimates from the best non-AFT model with Weibull frailty hazards for each replicate and also fixed-effect and random-effect meta-analysis estimates for each parameter of the model. Briefly, the fixed-effect meta-analysis estimates were derived using Inverse Variance Weighting (IVW) in which the estimates from each replicate were weighted by the inverse of the variance estimates. The meta-analysis estimates were then calculated simply as the weighted average of estimates from all replicates. The fixed-effect meta-analysis assumes there is insignificant variation in the estimates of the same parameter across different replicates. The random-effect meta-analysis derived the estimates by also assigning weights to estimates from each replicate, but the weights take into account the variation of estimates across replicates.

**Figure S9:**
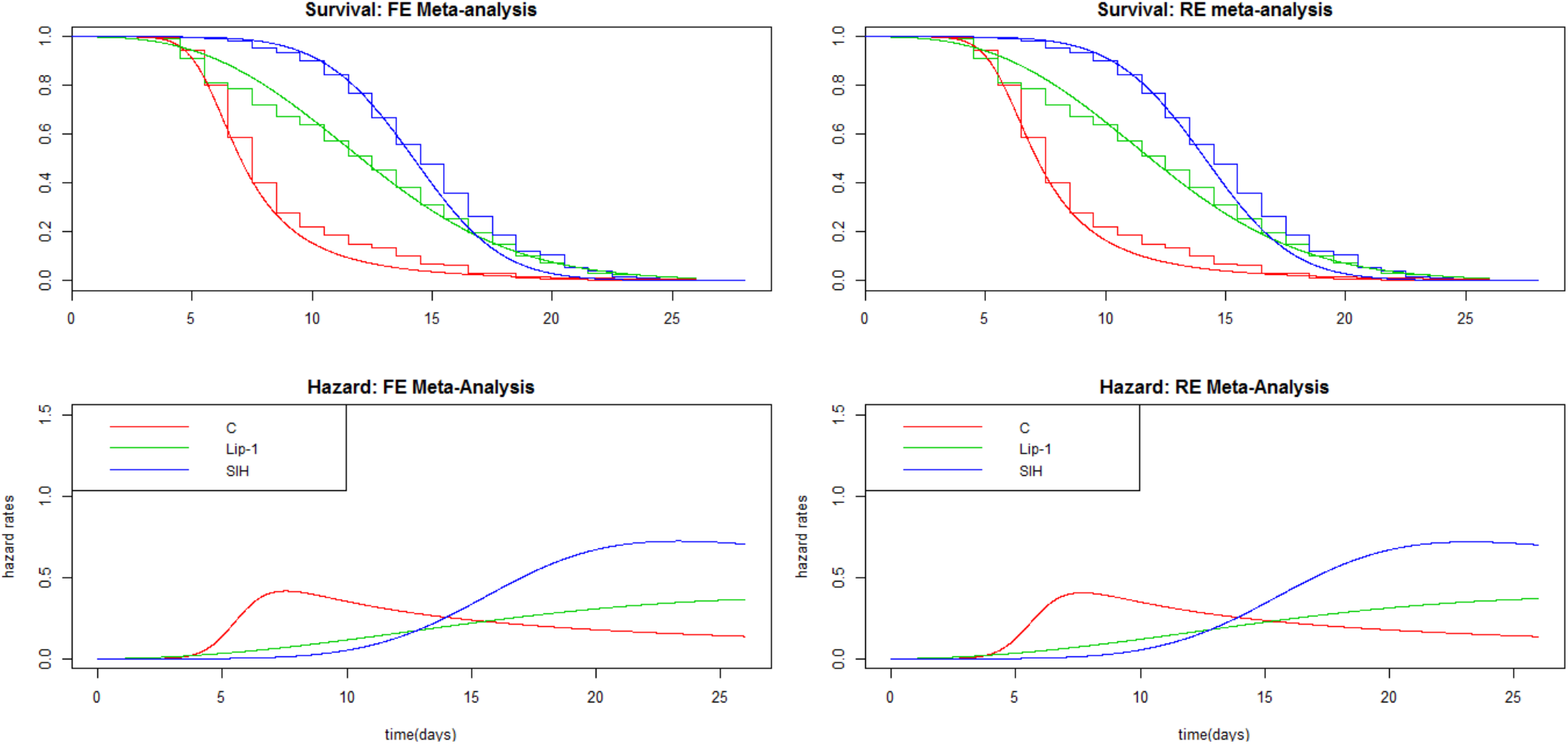
Meta-analysis Estimates of Survival and Hazard Functions. FE=fixed error, RE=random error.

**Table S13:**
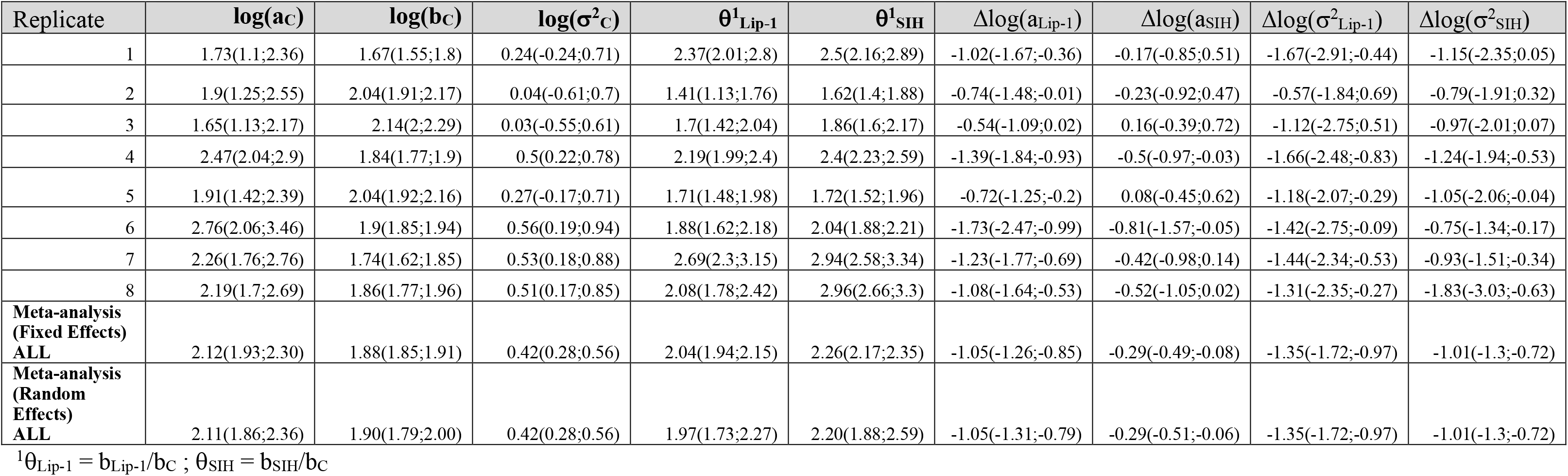
Parameter Estimates and their 95% confidence intervals from non-AFT Weibull-Frailty Models with treatment-dependent shape and heterogeneity parameters

| Replicate | $\log(ac)$ | $\log(bc)$ | $\log(\sigma^2_c)$ | $\theta^1_{Lip-1}$ | $\theta^1_{SIH}$ | $\Delta\log(a_{Lip-1})$ | $\Delta\log(a_{SIH})$ | $\Delta\log(\sigma^2_{Lip-1})$ | $\Delta\log(\sigma^2_{SIH})$ |
| --- | --- | --- | --- | --- | --- | --- | --- | --- | --- |
| 1 | 1.73(1.1;2.36) | 1.67(1.55;1.8) | 0.24(-0.24;0.71) | 2.37(2.01;2.8) | 2.5(2.16;2.89) | -1.02(-1.67;-0.36) | -0.17(-0.85;0.51) | -1.67(-2.91;-0.44) | -1.15(-2.35;0.05) |
| 2 | 1.9(1.25;2.55) | 2.04(1.91;2.17) | 0.04(-0.61;0.7) | 1.41(1.13;1.76) | 1.62(1.4;1.88) | -0.74(-1.48;-0.01) | -0.23(-0.92;0.47) | -0.57(-1.84;0.69) | -0.79(-1.91;0.32) |
| 3 | 1.65(1.13;2.17) | 2.14(2;2.29) | 0.03(-0.55;0.61) | 1.7(1.42;2.04) | 1.86(1.6;2.17) | -0.54(-1.09;0.02) | 0.16(-0.39;0.72) | -1.12(-2.75;0.51) | -0.97(-2.01;0.07) |
| 4 | 2.47(2.04;2.9) | 1.84(1.77;1.9) | 0.5(0.22;0.78) | 2.19(1.99;2.4) | 2.4(2.23;2.59) | -1.39(-1.84;-0.93) | -0.5(-0.97;-0.03) | -1.66(-2.48;-0.83) | -1.24(-1.94;-0.53) |
| 5 | 1.91(1.42;2.39) | 2.04(1.92;2.16) | 0.27(-0.17;0.71) | 1.71(1.48;1.98) | 1.72(1.52;1.96) | -0.72(-1.25;-0.2) | 0.08(-0.45;0.62) | -1.18(-2.07;-0.29) | -1.05(-2.06;-0.04) |
| 6 | 2.76(2.06;3.46) | 1.9(1.85;1.94) | 0.56(0.19;0.94) | 1.88(1.62;2.18) | 2.04(1.88;2.21) | -1.73(-2.47;-0.99) | -0.81(-1.57;-0.05) | -1.42(-2.75;-0.09) | -0.75(-1.34;-0.17) |
| 7 | 2.26(1.76;2.76) | 1.74(1.62;1.85) | 0.53(0.18;0.88) | 2.69(2.3;3.15) | 2.94(2.58;3.34) | -1.23(-1.77;-0.69) | -0.42(-0.98;0.14) | -1.44(-2.34;-0.53) | -0.93(-1.51;-0.34) |
| 8 | 2.19(1.7;2.69) | 1.86(1.77;1.96) | 0.51(0.17;0.85) | 2.08(1.78;2.42) | 2.96(2.66;3.3) | -1.08(-1.64;-0.53) | -0.52(-1.05;0.02) | -1.31(-2.35;-0.27) | -1.83(-3.03;-0.63) |
| **Meta-analysis**<br>**(Fixed Effects)**<br>**ALL** | 2.12(1.93;2.30) | 1.88(1.85;1.91) | 0.42(0.28;0.56) | 2.04(1.94;2.15) | 2.26(2.17;2.35) | -1.05(-1.26;-0.85) | -0.29(-0.49;-0.08) | -1.35(-1.72;-0.97) | -1.01(-1.3;-0.72) |
| **Meta-analysis**<br>**(Random Effects)**<br>**ALL** | 2.11(1.86;2.36) | 1.90(1.79;2.00) | 0.42(0.28;0.56) | 1.97(1.73;2.27) | 2.20(1.88;2.59) | -1.05(-1.31;-0.79) | -0.29(-0.51;-0.06) | -1.35(-1.72;-0.97) | -1.01(-1.3;-0.72) |

Conclusions from **Table S13**:

- There are between-replicate variations in the scaling factor estimates. But after combining across different replicates, θ_SIH_ estimate is 2.20(95% CI: 1.88;2.59) and θ_Lip-1_ estimate is 1.97(95% CI: 1.73;2.27), which means that life has significantly de-accelerated under Lip-1 and SIH relative to under controls, by about half.
- **Meta-analysis estimates also shows** that the treatment effect manifests in not only simple scaling of the Weibull scale parameter, but it also affects the shape parameters with the shape parameters for Lip-1 and SIH being significantly smaller (the 95% CI for differences in log shape parameter between Lip-1 and SIH versus controls do not include zero).
- **Meta-analysis estimates also shows** there is significant heterogeneity due to unobserved factors among control worms, 95% CI for log σ^2^_c_ = 0.42(95% CI: 0.28;0.56) while the heterogeneity due to unobserved factors are significantly less in Lip-1 and SIH worms.

**Table S14:** Comparison of Models for Combined Data

| Model | log(a) | log(b) | log($\sigma^2_c$) | $\theta_{\text{Lip-1}}$ | $\theta_{\text{SIH}}$ | $\Delta\log(a_{\text{Lip-1}})$ | $\Delta\log(a_{\text{SIH}})$ | $\Delta\log(\sigma^2_{\text{Lip-1}})$ | $\Delta\log(\sigma^2_{\text{SIH}})$ | BIC value |
| --- | --- | --- | --- | --- | --- | --- | --- | --- | --- | --- |
| M1 | same | different | same | same | same | 0 | 0 | 0 | 0 | 15835.49 |
| M2 | same | different | same | different | different | 0 | 0 | 0 | 0 | 15887.84 |
| M3 | same | different | same | same | same | 0 | 0 | same | same | 15490.08 |
| M4 | same | different | same | different | different | 0 | 0 | same | same | 15511.38 |
| M5 | same | different | same | same | same | same | same | same | same | 15109.10 |
| **M6** | **same** | **different** | **same** | **different** | **different** | **same** | **same** | **same** | **same** | **15079.13** |
| M7 | different | different | different | different | different | same | same | same | same | 15250.78 |
| Saturated | different | different | different | different | different | different | different | different | different | 15346.79 |

same = assumed the same across replicates
different = allowed to be different across replicates
0 = not estimated, set to zero

**Figure S10:**
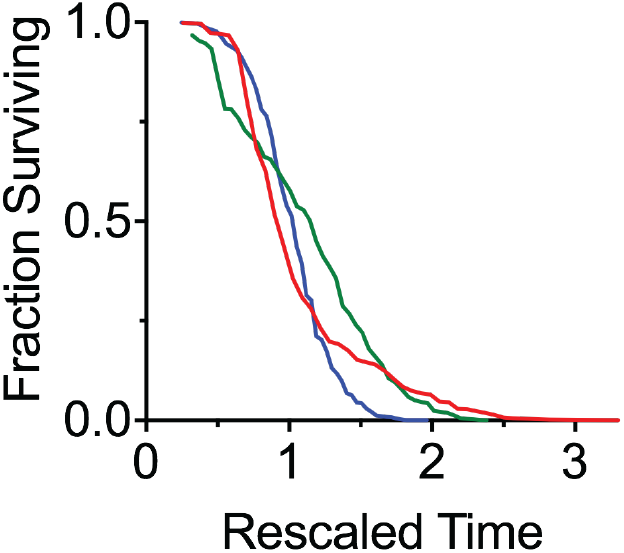
The best-fit AFT residuals from survival data meta-analysis shown in Figure S9, based the M6 model with fixed errors. Note that the survival curves cross rather than overlap, indicating temporal scaling is insufficient to explain the differences between treatments

### Investigating Temporal Rescaling by Temperature

To investigate if changing the temperature showed evidence of temporal scaling we compared worms aged at 20 °C and 25 °C during the same time frame. The effect of temperature is evaluated separately for control and SIH treated worms, population sizes for each replicate are shown in **Table S15**.

**Table S15:** Population sizes for replicates used to assess temporal rescaling by temperature

| Replicate | Temperature | control | SIH |
| --- | --- | --- | --- |
|  |  | death events | death events |
| 1 | 20 °C | 99 | 100 |
| 1 | 25 °C | 76 | 90 |
| 2 | 20 °C | 84 | 80 |
| 2 | 25 °C | 81 | 67 |

#### Checking AFT Assumption

For each of two replicates and treatment factors, the AFT model with Weibull baseline hazard was fitted with temperature as the covariate. The residuals from this model were then subjected to the K-S test. The *p*-values from the K-S test are given in **Table S16**. As can be seen, the *p*-values are generally not very small (only one *p*-value < 0.01), indicating that the evidence of departure from simple temporal rescaling is not strong. **Figure S11** shows the survival curves of the residuals and despite the discreteness, the two distributions (black for 20 °C and red for 25 °C) seem to be similar.

**Table S16:** *p*-values of KS test on Residuals of nonparametric AFT models

| Replicate | Treatment | $p$-value |
| --- | --- | --- |
| 1 | Control | $2.0 \times 10^{-2}$ |
| 1 | SIH | $3.5 \times 10^{-2}$ |
| 2 | Control | $2.7 \times 10^{-3}$ |
| 2 | SIH | $1.3 \times 10^{-1}$ |

**Figure S11:**
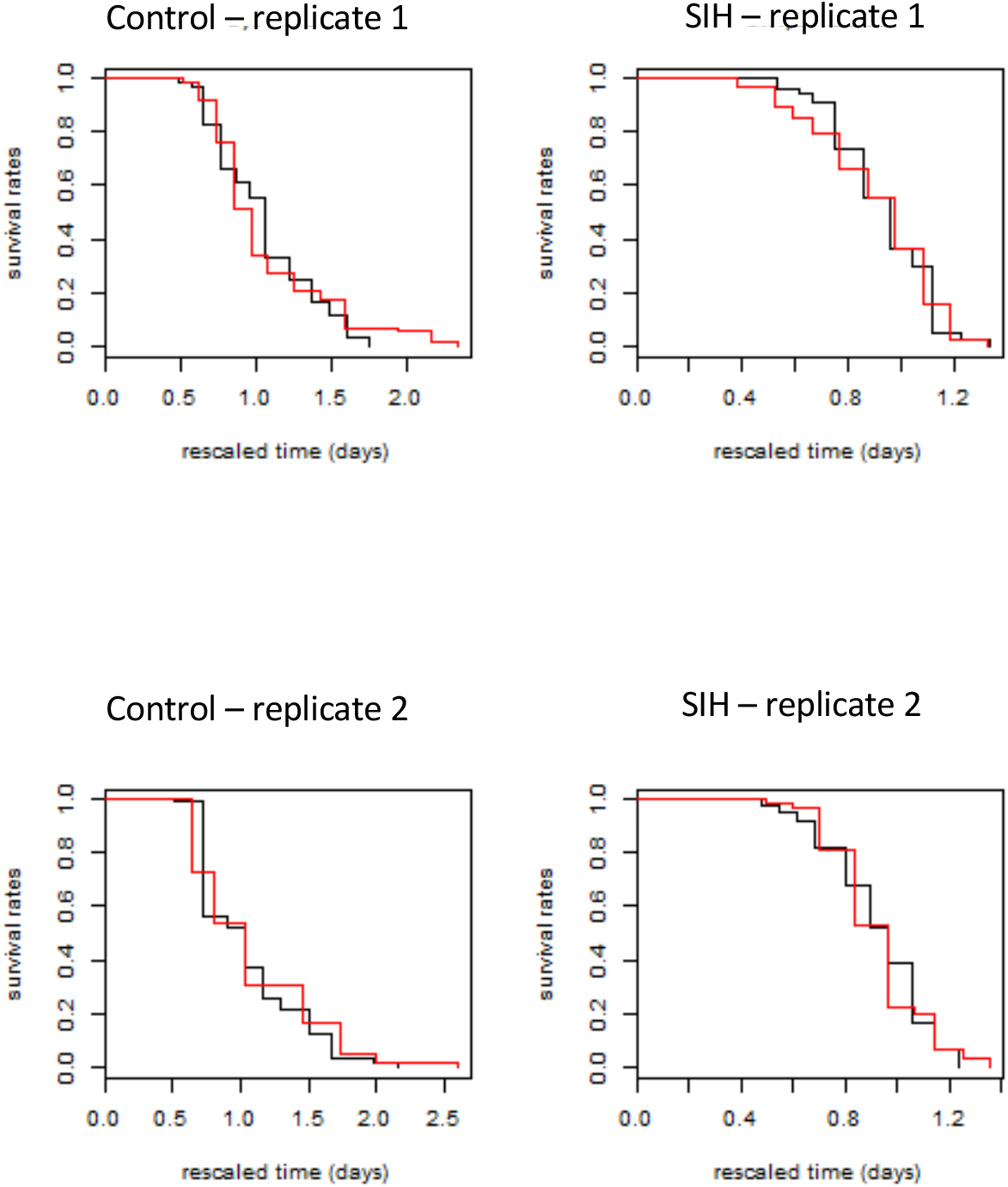
The best-fit AFT residuals from the model with Weibull baseline hazard for survival of control and SIH treated worms at 20 °C (black) and 25 °C (red)

#### Meta-Analysis

We also performed meta-analysis for each worm population and the results are given below. The log σ^2^ provides an indication of the level of heterogeneity, and it is interesting to note that the control worms exhibit greater heterogeneity than the SIH worms, consistent with that observed at 25 °C.

For both populations, being exposed to the higher temperature of 25 °C accelerates life as expected, by approximately 30% for control worms (θ_temp_ = **0.72 (95% CI: 0.68;0.77)** and 20% for SIH worms (θ_temp_ = **0.81 (95% CI: 0.78;0.84)**.

**Table S17:**
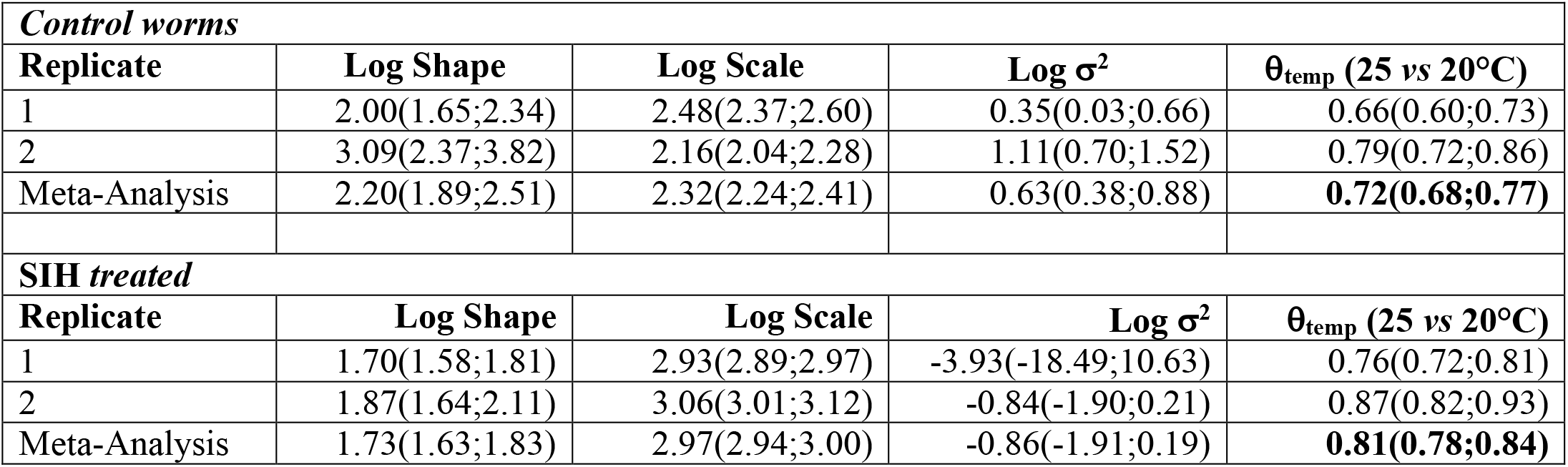
Meta-analysis results for control and SIH treated worms

### Body length analysis

Summary statistics and tests for normality of body length are included in **Table S18**. Not all length or derived data sets were normally distributed, as indicated below.

**Table S18:** Summary of body length comparisons between treatments and ages

|  | Start | Day 1<br>Control | Day 1<br>Lip-1 | Day 1<br>SIH | Day 4<br>Control | Day 4<br>Lip-1 | Day 4<br>SIH | Day 8<br>Control | Day 8<br>Lip-1 | Day 8<br>SIH |
| --- | --- | --- | --- | --- | --- | --- | --- | --- | --- | --- |
| Number of values | 125 | 82 | 98 | 90 | 71 | 49 | 68 | 17 | 10 | 21 |
| Minimum | 876.2 | 1159 | 1114 | 1024 | 934.2 | 1023 | 1377 | 1229 | 1171 | 1532 |
| 25% Percentile | 979.9 | 1341 | 1319 | 1371 | 1317 | 1277 | 1563 | 1346 | 1290 | 1651 |
| Median | 1026 | 1387 | 1364 | 1412 | 1432 | 1345 | 1614 | 1403 | 1463 | 1704 |
| 75% Percentile | 1084 | 1432 | 1403 | 1443 | 1519 | 1445 | 1648 | 1561 | 1524 | 1742 |
| Maximum | 1271 | 1527 | 1488 | 1521 | 1660 | 1589 | 1773 | 1619 | 1554 | 1815 |
| Mean | 1036 | 1387 | 1358 | 1400 | 1414 | 1344 | 1605 | 1440 | 1414 | 1696 |
| Std. Deviation | 80.61 | 67.69 | 66.66 | 80.33 | 146.2 | 128.5 | 77.57 | 122.7 | 133.9 | 63.91 |
| Std. Error of Mean | 7.21 | 7.475 | 6.734 | 8.467 | 17.35 | 18.36 | 9.406 | 29.76 | 42.33 | 13.95 |
| Lower 95% CI of mean | 1022 | 1372 | 1345 | 1383 | 1379 | 1308 | 1586 | 1376 | 1318 | 1667 |
| Upper 95% CI of mean | 1051 | 1402 | 1372 | 1417 | 1448 | 1381 | 1624 | 1503 | 1509 | 1725 |
| D'Agostino & Pearson normality test |  |  |  |  |  |  |  |  |  |  |
| K2 | 6.981 | 5.546 | 18.41 | 63.3 | 8.069 | 5.393 | 7.285 | 3.174 | 1.47 | 1.822 |
| *p* value | 0.0305 | 0.0625 | 0.0001 | <0.0001 | 0.0177 | 0.0675 | 0.0262 | 0.2045 | 0.4794 | 0.4021 |
| **Passed normality test ($\alpha=0.05$)?** | **No** | **Yes** | **No** | **No** | **No** | **Yes** | **No** | **Yes** | **Yes** | **Yes** |
| *p* value summary | * | ns | *** | **** | * | ns | * | ns | ns | ns |

To compare between age and treatment groups a Kruskal-Wallace ANOVA was performed, followed by Dunn’s multiple comparisons Post-hoc tests. There was a significant difference between body length (H(10)= 432.6, *p* < 0.0001) amongst the groups measured. The results of the pairwise comparisons, corrected for multiple comparisons, are shown in **Table S19.**

**Table S19:**
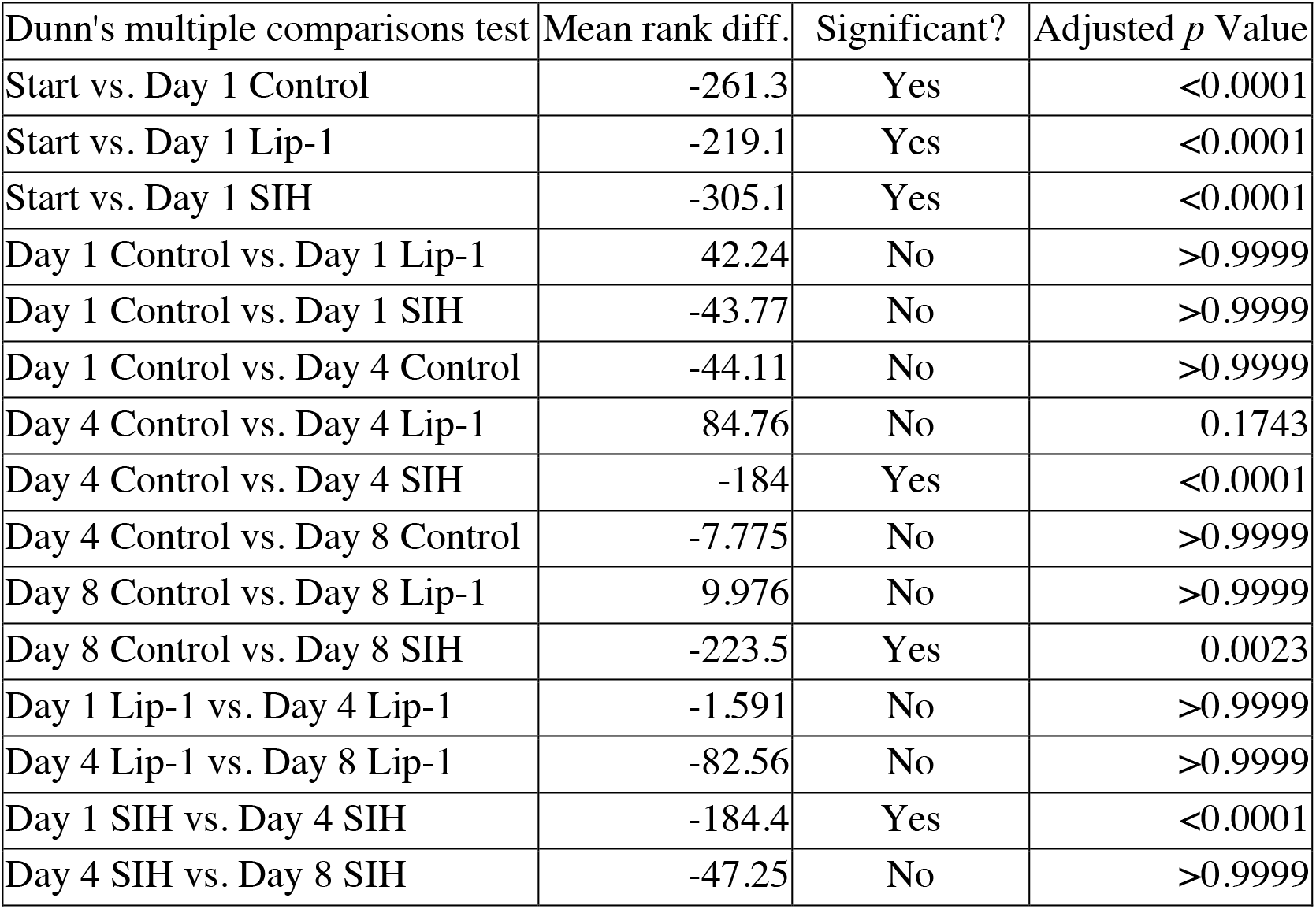
Summary of body length comparisons between ages and treatments.

| Dunn's multiple comparisons test | Mean rank diff. | Significant? | Adjusted *p* Value |
| --- | --- | --- | --- |
| Start vs. Day 1 Control | -261.3 | Yes | <0.0001 |
| Start vs. Day 1 Lip-1 | -219.1 | Yes | <0.0001 |
| Start vs. Day 1 SIH | -305.1 | Yes | <0.0001 |
| Day 1 Control vs. Day 1 Lip-1 | 42.24 | No | >0.9999 |
| Day 1 Control vs. Day 1 SIH | -43.77 | No | >0.9999 |
| Day 1 Control vs. Day 4 Control | -44.11 | No | >0.9999 |
| Day 4 Control vs. Day 4 Lip-1 | 84.76 | No | 0.1743 |
| Day 4 Control vs. Day 4 SIH | -184 | Yes | <0.0001 |
| Day 4 Control vs. Day 8 Control | -7.775 | No | >0.9999 |
| Day 8 Control vs. Day 8 Lip-1 | 9.976 | No | >0.9999 |
| Day 8 Control vs. Day 8 SIH | -223.5 | Yes | 0.0023 |
| Day 1 Lip-1 vs. Day 4 Lip-1 | -1.591 | No | >0.9999 |
| Day 4 Lip-1 vs. Day 8 Lip-1 | -82.56 | No | >0.9999 |
| Day 1 SIH vs. Day 4 SIH | -184.4 | Yes | <0.0001 |
| Day 4 SIH vs. Day 8 SIH | -47.25 | No | >0.9999 |

### Volume analysis

Summary statistics and tests for normality of total worm volume are included in **Table S20**. Not all length or derived data sets were normally distributed, as indicated below.

**Table S20:** Summary of volume comparisons between treatments and ages

|  | Start | Day 1<br>Control | Day 1 Lip-1 | Day 1 SIH | Day 4<br>Control | Day 4 Lip-1 | Day 4 SIH | Day 8<br>Control | Day 8 Lip-1 | Day 8 SIH |
| --- | --- | --- | --- | --- | --- | --- | --- | --- | --- | --- |
| Number of values | 125 | 82 | 98 | 90 | 71 | 49 | 68 | 17 | 10 | 21 |
| Minimum | 747.8 | 3115 | 2740 | 1731 | 3293 | 2258 | 4052 | 4939 | 4203 | 5461 |
| 25% Percentile | 1589 | 4051 | 3883 | 4057 | 5049 | 4251 | 6677 | 5655 | 4882 | 7280 |
| Median | 1867 | 4247 | 4293 | 4393 | 6103 | 5032 | 7183 | 6673 | 6147 | 7726 |
| 75% Percentile | 2083 | 4576 | 4598 | 4740 | 6507 | 5648 | 7738 | 7465 | 6752 | 8407 |
| Maximum | 3818 | 5249 | 5420 | 5800 | 8267 | 6712 | 8608 | 8696 | 7876 | 9210 |
| Mean | 1901 | 4289 | 4209 | 4369 | 5810 | 4970 | 7111 | 6511 | 5958 | 7731 |
| Std. Deviation | 495.1 | 417.5 | 540.8 | 643.1 | 1111 | 965.4 | 861.8 | 1043 | 1155 | 825.4 |
| Std. Error of Mean | 44.28 | 46.1 | 54.63 | 67.79 | 131.8 | 137.9 | 104.5 | 252.9 | 365.2 | 180.1 |
| Lower 95% CI of mean | 1813 | 4197 | 4101 | 4234 | 5547 | 4693 | 6903 | 5975 | 5132 | 7355 |
| Upper 95% CI of mean | 1988 | 4380 | 4318 | 4504 | 6073 | 5248 | 7320 | 7047 | 6784 | 8106 |
| D'Agostino & Pearson normality test |  |  |  |  |  |  |  |  |  |  |
| K2 | 38.42 | 0.4213 | 2.657 | 37.57 | 2.331 | 1.45 | 12.99 | 0.3558 | 0.1231 | 4.164 |
| *p* value | <0.0001 | 0.8101 | 0.2649 | <0.0001 | 0.3118 | 0.4843 | 0.0015 | 0.8370 | 0.9403 | 0.1247 |
| Passed normality test ($\alpha=0.05$)? | No | Yes | Yes | No | Yes | Yes | No | Yes | Yes | Yes |
| *p* value summary | **** | ns | ns | **** | ns | ns | ** | ns | ns | ns |

There was a significant difference between body volume (H(10)= 489, *p* < 0.0001) amongst the groups measured. Comparisons between age and treatment groups a Kruskal-Wallace ANOVA was performed, followed by Dunn’s multiple comparisons Post-hoc tests. The results of the pairwise comparisons, corrected for multiple comparisons, are shown in **Table S21**.

**Table S21:** Summary of volume comparisons between ages and treatments.

| Dunn's multiple comparisons test | Mean rank diff. | Significant? | Adjusted $p$ Value |
| --- | --- | --- | --- |
| Start vs. Day 1 Control | -214.9 | Yes | <0.0001 |
| Start vs. Day 1 Lip-1 | -205.6 | Yes | <0.0001 |
| Start vs. Day 1 SIH | -236.4 | Yes | <0.0001 |
| Day 1 Control vs. Day 1 Lip-1 | 9.301 | No | >0.9999 |
| Day 1 Control vs. Day 1 SIH | -21.45 | No | >0.9999 |
| Day 1 Control vs. Day 4 Control | -175 | Yes | <0.0001 |
| Day 4 Control vs. Day 4 Lip-1 | 80.42 | No | 0.2630 |
| Day 4 Control vs. Day 4 SIH | -102.6 | Yes | 0.0136 |
| Day 4 Control vs. Day 8 Control | -66.73 | No | >0.9999 |
| Day 8 Control vs. Day 8 Lip-1 | 54.78 | No | >0.9999 |
| Day 8 Control vs. Day 8 SIH | -70.02 | No | >0.9999 |
| Day 1 Lip-1 vs. Day 4 Lip-1 | -103.9 | Yes | 0.0169 |
| Day 4 Lip-1 vs. Day 8 Lip-1 | -92.37 | No | >0.9999 |
| Day 1 SIH vs. Day 4 SIH | -256.2 | Yes | <0.0001 |
| Day 4 SIH vs. Day 8 SIH | -34.11 | No | >0.9999 |

### Movement analysis

Movement parameters measured included maximum velocity (shown in **Figure 6e**), mean velocity and total distance travelled. The latter two datasets are represented in **Figures S12** and **S13**.

**Figure S12.**
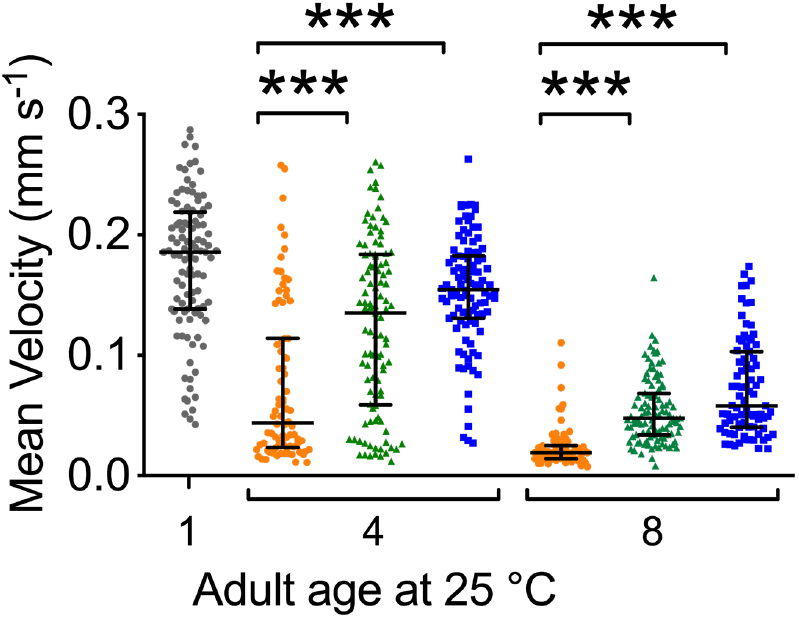
Estimates of mean velocity (mm s-1) achieved by aged and treated cohorts of *C. elegans*. (Grey=starting population, red=control, green=Lip-1 treated, blue=SIH-treated) Treatment with either Lip-1 or SIH attenuates the age-related decline in mean velocity (Kruskal-Wallis ANOVA: H(7)= 339.2, *p* < 0.0001), see **Supplemental Table S24** for summary data and **Table S25** for pair-wise comparisons).

**Figure S13.**
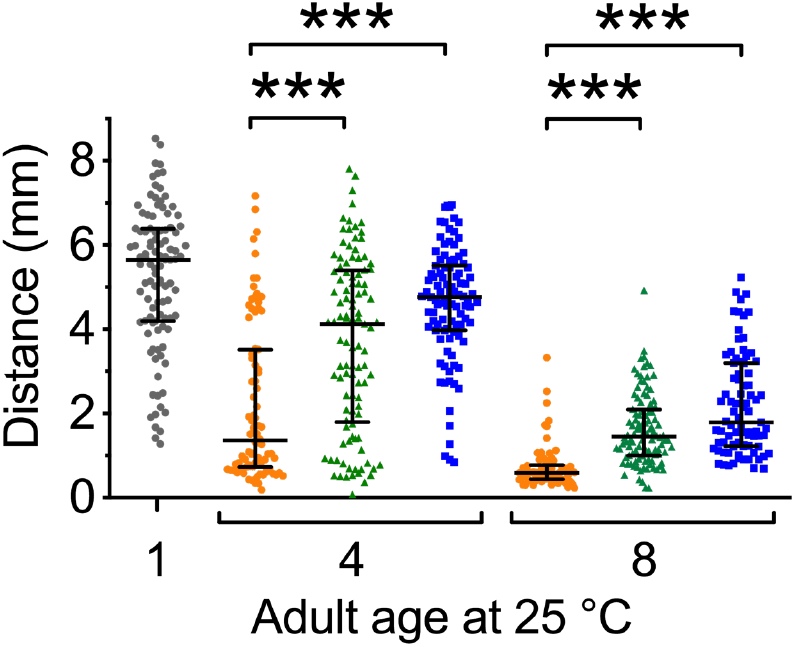
Estimates of total distance travelled (in 30 s) by aged and treated cohorts of *C. elegans*. (Grey=starting population, red=control, green=Lip-1 treated, blue=SIH-treated) Treatment with either Lip-1 or SIH attenuates the age-related decline in distance travelled (Kruskal-Wallis ANOVA: H(7)= 340.6, *p* < 0.0001)see **Supplemental Table S26** for summary data and **Table S27** for pair-wise comparisons).

Summary statistics and tests for normality of maximum velocity (mm s^−1^) are included in **Table S22**. Not all data sets were normally distributed, as indicated below.

**Table S22:** Summary of maximum velocity results across treatments and ages

|  | Day 1<br>Control | Day 4<br>Control | Day 8<br>Control | Day 4 SIH | Day 8 SIH | Day 4 Lip-1 | Day 8 Lip-1 |
| --- | --- | --- | --- | --- | --- | --- | --- |
| Number of values | 103 | 82 | 73 | 99 | 83 | 105 | 104 |
| Minimum | 0.1832 | 0.04309 | 0.03738 | 0.105 | 0.1028 | 0.06351 | 0.05622 |
| 25% Percentile | 0.3731 | 0.1294 | 0.06372 | 0.3442 | 0.174 | 0.2711 | 0.1478 |
| Median | 0.4509 | 0.2304 | 0.09115 | 0.3733 | 0.2513 | 0.3762 | 0.1922 |
| 75% Percentile | 0.5491 | 0.3364 | 0.1254 | 0.4755 | 0.3506 | 0.4861 | 0.2452 |
| Maximum | 0.8264 | 0.6134 | 0.3802 | 0.7127 | 0.6458 | 0.7204 | 0.5563 |
| Mean | 0.4692 | 0.2536 | 0.109 | 0.392 | 0.2847 | 0.37 | 0.2084 |
| Std. Deviation | 0.1457 | 0.1496 | 0.06568 | 0.1087 | 0.1356 | 0.1588 | 0.09062 |
| Std. Error of Mean | 0.01435 | 0.01653 | 0.007687 | 0.01093 | 0.01489 | 0.01549 | 0.008886 |
| Lower 95% CI of mean | 0.4408 | 0.2207 | 0.09367 | 0.3704 | 0.2551 | 0.3393 | 0.1908 |
| Upper 95% CI of mean | 0.4977 | 0.2865 | 0.1243 | 0.4137 | 0.3143 | 0.4007 | 0.226 |
| D'Agostino & Pearson normality test |  |  |  |  |  |  |  |
| K2 | 4.785 | 8.157 | 44.36 | 0.9424 | 10.76 | 3.145 | 23.3 |
| $p$ value | 0.0914 | 0.0169 | <0.0001 | 0.6242 | 0.0046 | 0.2075 | <0.0001 |
| Passed normality test ($\alpha=0.05$)? | Yes | No | No | Yes | No | Yes | No |
| $p$ value summary | ns | * | **** | ns | ** | ns | **** |

To compare between age and treatment groups a Kruskal-Wallace ANOVA was performed, followed by Dunn’s multiple comparisons post-hoc tests. There was a significant difference between maximum velocity (H(7)=298.5, *p* < 0.0001) amongst the groups measured. The results of the pairwise comparisons, corrected for multiple comparisons, are shown in **Table S23.**

**Table S23:** Summary of maximum velocity comparisons between ages and treatments.

| Dunn's multiple comparisons test | Mean rank diff. | Significant? | Adjusted $p$ Value |
| --- | --- | --- | --- |
| Day 1 Control vs. Day 4 Control | 230.5 | Yes | <0.0001 |
| Day 1 Control vs. Day 8 Control | 411.2 | Yes | <0.0001 |
| Day 4 Control vs. Day 8 Control | 180.7 | Yes | <0.0001 |
| Day 4 Control vs. Day 4 SIH | -168.4 | Yes | <0.0001 |
| Day 4 Control vs. Day 4 Lip-1 | -133.6 | Yes | <0.0001 |
| Day 8 Control vs. Day 8 SIH | -220.9 | Yes | <0.0001 |
| Day 8 Control vs. Day 8 Lip-1 | -132.7 | Yes | <0.0001 |
| Day 4 SIH vs. Day 4 Lip-1 | 34.84 | No | >0.9999 |
| Day 8 SIH vs. Day 8 Lip-1 | 88.29 | Yes | 0.0110 |

#### Mean velocity

Summary statistics and tests for normality of mean velocity (mm s^−1^) are included in **Table S24**. Not all data sets were normally distributed, as indicated below.

**Table S24:** Summary of mean velocity results across treatments and ages

|  | Day 1 Control | Day 4 Control | Day 8 Control | Day 4 SIH | Day 8 SIH | Day 4 Lip-1 | Day 8 Lip-1 |
| --- | --- | --- | --- | --- | --- | --- | --- |
| Number of values | 103 | 82 | 73 | 99 | 83 | 105 | 104 |
| Minimum | 0.04246 | 0.01112 | 0.007567 | 0.02713 | 0.02239 | 0.01219 | 0.008319 |
| 25% Percentile | 0.1382 | 0.02355 | 0.01407 | 0.1307 | 0.04028 | 0.05892 | 0.03386 |
| Median | 0.1856 | 0.04389 | 0.01907 | 0.1545 | 0.05813 | 0.1349 | 0.04763 |
| 75% Percentile | 0.2189 | 0.1139 | 0.02507 | 0.1824 | 0.1029 | 0.1839 | 0.06818 |
| Maximum | 0.2873 | 0.2578 | 0.1104 | 0.2629 | 0.1739 | 0.2606 | 0.1644 |
| Mean | 0.1756 | 0.07449 | 0.02376 | 0.1528 | 0.07262 | 0.1244 | 0.05407 |
| Std. Deviation | 0.05753 | 0.06507 | 0.01791 | 0.0464 | 0.04028 | 0.07094 | 0.02726 |
| Std. Error of Mean | 0.005669 | 0.007186 | 0.002096 | 0.004663 | 0.004421 | 0.006923 | 0.002673 |
| Lower 95% CI of mean | 0.1644 | 0.06019 | 0.01958 | 0.1435 | 0.06382 | 0.1106 | 0.04877 |
| Upper 95% CI of mean | 0.1868 | 0.08878 | 0.02793 | 0.162 | 0.08141 | 0.1381 | 0.05937 |
| D'Agostino & Pearson normality test |  |  |  |  |  |  |  |
| K2 | 3.86 | 13.86 | 66.08 | 6.268 | 8.786 | 29.36 | 21.66 |
| $p$ value | 0.1451 | 0.0010 | <0.0001 | 0.0435 | 0.0124 | <0.0001 | <0.0001 |
| Passed normality test ($\alpha=0.05$)? | Yes | No | No | No | No | No | No |
| $p$ value summary | ns | *** | **** | * | * | **** | **** |

To compare between age and treatment groups a Kruskal-Wallace ANOVA was performed, followed by Dunn’s multiple comparisons Post-hoc tests. There was a significant difference between mean velocity (H(7)= 339.2, *p* < 0.0001) amongst the groups measured. The results of the pairwise comparisons, corrected for multiple comparisons, are shown in **Table S25.**

**Table S25:** Summary of mean velocity comparisons between ages and treatments.

| Dunn's multiple comparisons test | Mean rank diff. | Significant? | Adjusted *p* Value |
| --- | --- | --- | --- |
| Day 1 Control vs. Day 4 Control | 259.2 | Yes | <0.0001 |
| Day 1 Control vs. Day 8 Control | 427.8 | Yes | <0.0001 |
| Day 4 Control vs. Day 8 Control | 168.6 | Yes | <0.0001 |
| Day 4 Control vs. Day 4 SIH | -215.1 | Yes | <0.0001 |
| Day 4 Control vs. Day 4 Lip-1 | -135.7 | Yes | <0.0001 |
| Day 8 Control vs. Day 8 SIH | -192.7 | Yes | <0.0001 |
| Day 8 Control vs. Day 8 Lip-1 | -141.2 | Yes | <0.0001 |
| Day 4 SIH vs. Day 4 Lip-1 | 79.45 | Yes | 0.0224 |
| Day 8 SIH vs. Day 8 Lip-1 | 51.53 | No | 0.5569 |

#### Total distance travelled

Summary statistics and tests for normality of total distance travelled (mm) are included in **Table S26**. Not all data were normally distributed, as indicated below.

**Table S26:** Summary of distance travelled results across treatments and ages

|  | Day 1<br>Control | Day 4<br>Control | Day 8<br>Control | Day 4<br>SIH | Day 8<br>SIH | Day 4<br>Lip-1 | Day 8<br>Lip-1 |
| --- | --- | --- | --- | --- | --- | --- | --- |
| Number of values | 103 | 82 | 73 | 99 | 83 | 105 | 104 |
| Minimum | 1.28 | 0.1883 | 0.2346 | 0.8402 | 0.6912 | 0.07633 | 0.2338 |
| 25% Percentile | 4.197 | 0.7269 | 0.4349 | 3.977 | 1.224 | 1.797 | 1.001 |
| Median | 5.649 | 1.357 | 0.5888 | 4.766 | 1.784 | 4.126 | 1.451 |
| 75% Percentile | 6.388 | 3.521 | 0.7714 | 5.513 | 3.193 | 5.397 | 2.092 |
| Maximum | 8.534 | 7.169 | 3.324 | 6.951 | 5.235 | 7.817 | 4.926 |
| Mean | 5.24 | 2.216 | 0.7247 | 4.619 | 2.192 | 3.685 | 1.627 |
| Std. Deviation | 1.68 | 1.865 | 0.5329 | 1.349 | 1.192 | 2.064 | 0.849 |
| Std. Error of Mean | 0.1656 | 0.2059 | 0.06237 | 0.1356 | 0.1308 | 0.2014 | 0.08325 |
| Lower 95% CI of mean | 4.911 | 1.806 | 0.6003 | 4.35 | 1.932 | 3.285 | 1.462 |
| Upper 95% CI of mean | 5.568 | 2.626 | 0.849 | 4.888 | 2.452 | 4.084 | 1.792 |
| D'Agostino & Pearson normality test |  |  |  |  |  |  |  |
| K2 | 4.494 | 10.94 | 63.28 | 9.719 | 8.451 | 27.91 | 16.99 |
| *p* value | 0.1057 | 0.0042 | <0.0001 | 0.0078 | 0.0146 | <0.0001 | 0.0002 |
| Passed normality test ($\alpha=0.05$)? | Yes | No | No | No | No | No | No |
| *p* value summary | ns | ** | **** | ** | * | **** | *** |

To compare between age and treatment groups a Kruskal-Wallace ANOVA was performed, followed by Dunn’s multiple comparisons Post-hoc tests. There was a significant difference between total distance travelled (H(7)= 340.6, *p*< 0.0001) amongst the groups measured. The results of the pairwise comparisons, corrected for multiple comparisons, are shown in **Table S27.**

**Table S27:**
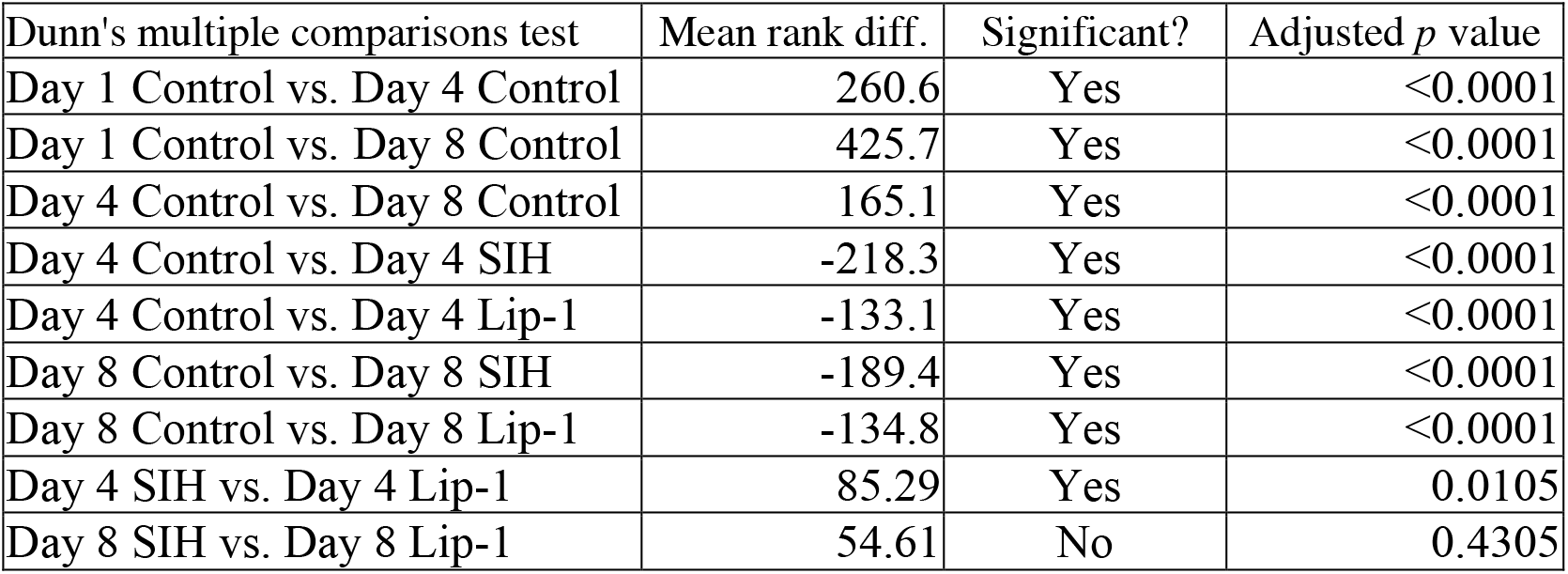
Summary of total distance travelled comparisons between ages and treatments.

| Dunn's multiple comparisons test | Mean rank diff. | Significant? | Adjusted $p$ value |
| --- | --- | --- | --- |
| Day 1 Control vs. Day 4 Control | 260.6 | Yes | <0.0001 |
| Day 1 Control vs. Day 8 Control | 425.7 | Yes | <0.0001 |
| Day 4 Control vs. Day 8 Control | 165.1 | Yes | <0.0001 |
| Day 4 Control vs. Day 4 SIH | -218.3 | Yes | <0.0001 |
| Day 4 Control vs. Day 4 Lip-1 | -133.1 | Yes | <0.0001 |
| Day 8 Control vs. Day 8 SIH | -189.4 | Yes | <0.0001 |
| Day 8 Control vs. Day 8 Lip-1 | -134.8 | Yes | <0.0001 |
| Day 4 SIH vs. Day 4 Lip-1 | 85.29 | Yes | 0.0105 |
| Day 8 SIH vs. Day 8 Lip-1 | 54.61 | No | 0.4305 |

#### Correlation of estimated movement parameters

Pooling all groups and ages reveals that all movement parameters (maximum velocity, mean velocity and distance travelled in 30s) are all positively correlated, as shown in **Figure S14**.

**Figure S14.**
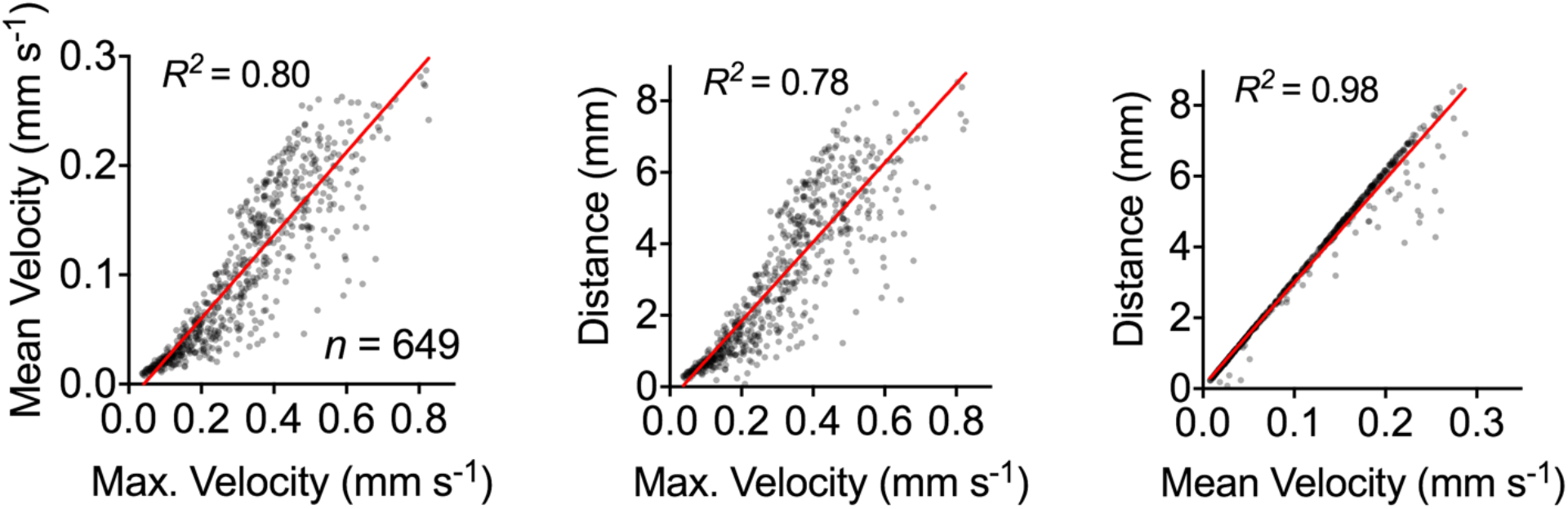
Correlation of estimated movement parameters.

### Antibiotic tests

To determine whether the increased lifespan seen with SIH treatment could be explained solely by an antibiotic effect of iron reduction, nematodes were treated with ampicillin, with and without SIH co-administration. Even in the presence of ampicillin, SIH increased median lifespan by 6 days, similar to its benefits in the absence of ampicillin (median increase of 7 days), as shown in **Figure S15**.

**Figure S15:**
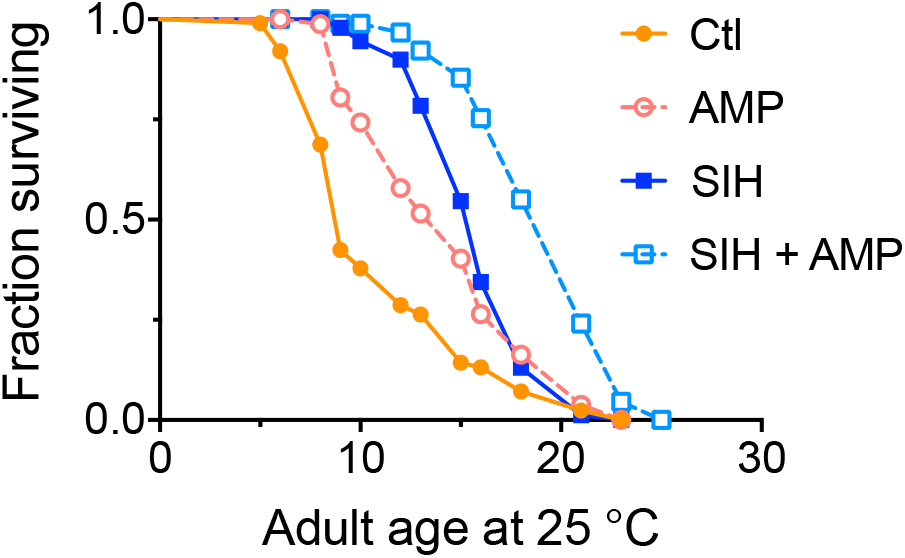
Administration of the antibiotic ampicillin and SIH. AMP= 5 µg / ml ampicillin. Median survival - Log-rank (Mantel-Cox) test: Control (Ctl) =9 days; AMP=15 days (*p*<0.0001); 250 µM SIH=16 days (*p*<0.0001); 250 µM SIH + AMP=21 days (*p*<0.0001).

